# Reconstruction of networks with direct and indirect genetic effects

**DOI:** 10.1101/646208

**Authors:** Willem Kruijer, Pariya Behrouzi, Daniela Bustos-Korts, María Xosé Rodríguez-Álvarez, Seyed Mahdi Mahmoudi, Brian Yandell, Ernst Wit, Fred A. van Eeuwijk

## Abstract

Genetic variance of a phenotypic trait can originate from direct genetic effects, or from indirect effects, i.e., through genetic effects on other traits, affecting the trait of interest. This distinction is often of great importance, for example when trying to improve crop yield and simultaneously controlling plant height. As suggested by Sewall Wright, assessing contributions of direct and indirect effects requires knowledge of (1) the presence or absence of direct genetic effects on each trait, and (2) the functional relationships between the traits. Because experimental validation of such relationships is often unfeasible, it is increasingly common to reconstruct them using causal inference methods. However, most of the current methods require all genetic variance to be explained by a small number of QTLs with fixed effects. Only few authors considered the ‘missing heritability’ case, where contributions of many undetectable QTLs are modelled with random effects. Usually, these are treated as nuisance terms, that need to be eliminated by taking residuals from a multi-trait mixed model (MTM). But fitting such MTM is challenging, and it is impossible to infer the presence of direct genetic effects. Here we propose an alternative strategy, where genetic effects are formally included in the graph. This has important advantages: (1) genetic effects can be directly incorporated in causal inference, implemented via our PCgen algorithm, which can analyze many more traits and (2) we can test the existence of direct genetic effects and improve the orientation of edges between traits. Finally, we show that reconstruction is much more accurate if individual plant or plot data are used, instead of genotypic means. We have implemented the PCgen-algorithm in the R-package pcgen.

## Introduction

To attain higher genetic gains, modern plant and animal breeders increasingly scale up their programs via the implementation of genomic prediction technologies (Cooper *et al.* 2014). Most of the genomic prediction applications are based on linear mixed- or Bayesian models that predict the phenotype for the target trait (yield) as a function of a multivariate distribution for SNP effects. In these models, the physiological mechanisms and traits that modulate the genotypic response to the environment over time are modeled implicitly via the SNP effects on the target trait (Zhou and Stephens 2014; Calus and Veerkamp 2011). The availability of high throughput phenotyping technologies has enabled breeders to characterize additional traits and monitor growth and development during the season. This opens new opportunities in breeding strategies, in which better-adapted genotypes result from combining loci that regulate complementary physiological mechanisms. This kind of breeding strategy is called physiological breeding (Reynolds and Langridge 2016). In physiological breeding, prediction accuracy for the target trait benefits from modeling the target trait and its underlying traits simultaneously (van Eeuwijk *et al.* 2018) because of a larger power to estimate its underlying effects (Stephens 2013). The challenge of seeking complementary traits is that many observed traits will be highly correlated to each other. Therefore, it is hard to disentangle the causal mechanisms modulating the genotypic response over time, which could potentially lead to larger prediction accuracy and to a transgressive segregation for the target trait. Structural equation models (SEMs), proposed by Wright (Wright 1921), and extended with random genetic effects in Gianola and Sorensen (2004), are a promising approach to deal with this problem (Rosa *et al.* 2011). In SEMs each trait is modelled explicitly as a function of the other traits and a noise term. Therefore, SEMs are a useful tool to identify which are the key traits that could be selection targets, or be incorporated in multi-trait genomic prediction models to improve the prediction accuracy for the target trait (van Eeuwijk *et al.* 2018).

A first advantage of SEMs, compared to regression models, is that one can predict the behavior of the system when one or more of the structural equations are modified by some kind of intervention. Figure 1 shows an illustration of an intervention. For example, a question that could be interesting to plant breeders is: which would be the contribution of a trait (say, radiation use efficiency) to yield, if flowering time is fixed for all genotypes at a particular value?

**Figure. 1.**
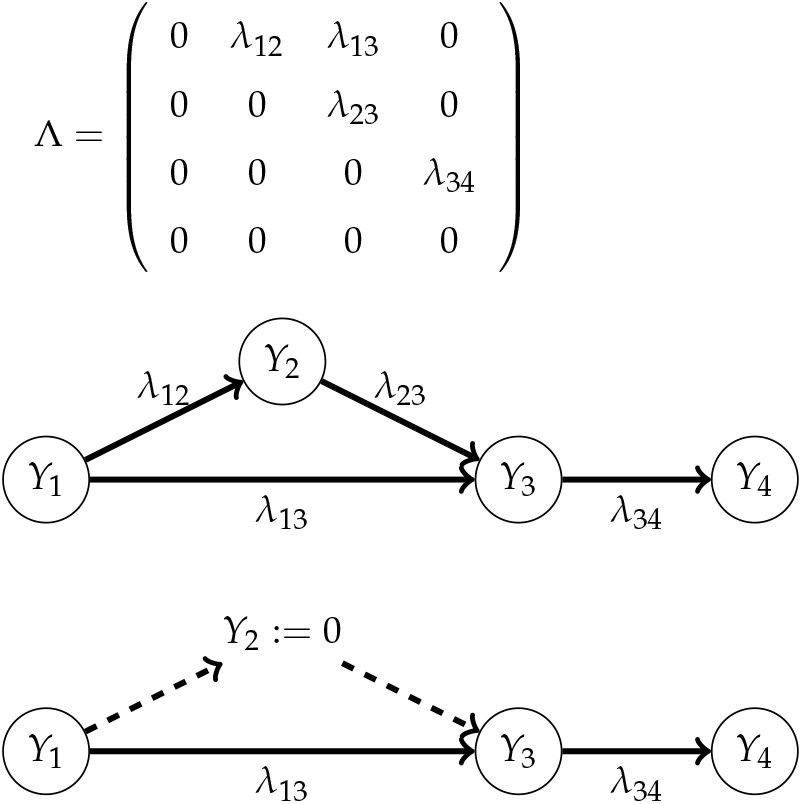
An example of a linear SEM. The SEM can be represented by a graph (middle), which is defined by the nonzero elements of Λ, the matrix containing the path coefficients (top). The total effect of *ϒ*_1_ on *ϒ*_4_ can be obtained by summing the contributions of the directed paths *ϒ*_1_ ϒ *ϒ*_3_ ϒ *ϒ*_4_ and *ϒ*_1_ ϒ *ϒ*_2_ ϒ *ϒ*_3_ ϒ *ϒ*_4_, where each contribution is the product of the corresponding path coefficients. After the intervention *ϒ*_2_ := 0 (bottom), the effect changes from (*λ*_13_ *λ*_34_ + *λ*_12_*λ*_23_*λ*_34_) to *λ*_13_*λ*_34_.

Second, SEMs make possible to distinguish between direct and indirect effects of one trait on another, and similarly between direct and indirect genetic effects. For example, let plant height (trait **ϒ_1_**) be modelled as **ϒ_1_**= **G_1_**+ **E_1_**, i.e., as the sum of a random genetic and residual effect, where all terms are *n*× 1 vectors, containing the values for a population of *n* individuals. Suppose plant height has a linear effect on yield (**ϒ_2_**), with additional random effects **G_2_** and **E_2_**:

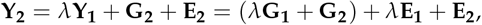

where *λ* is the path (or structural) coefficient associated with the effect of **ϒ_1_** on **ϒ_2_**. On the one hand, we have the *direct* genetic effects **G_1_** and **G_2_**; on the other hand, we have the *total* (or marginal) genetic effects **U_1_**= **G_1_** and **U_2_**= *λ***G_1_** + **G_2_**, the indirect effects being **U_1_** − **G_1_**= 0 and **U_2_** − **G_2_**= *λ***G_1_**. Similarly, we can distinguish between the 2 × 2 matrices Σ_*G*_, containing the (co)variances of **G_1_** and **G_2_**, and *V_G_*, with the (co)variances of **U_1_** and **U_2_**. The latter is the matrix of genetic (co)variances, appearing in the usual MTM (multi-trait mixed model) for **ϒ_1_** and **ϒ_2_**; here it is a function of Σ_*G*_ and *λ*.

Knowledge of the direct genetic effects is in principle of great interest to breeders (Valente *et al.* 2013, 2015). However, routine use of these models is currently difficult for two reasons. First, for a given SEM, not all parameters may be identifiable, i.e., because of overparameterization different sets of parameter values can lead to the same model, making estimation infeasible. Gianola and Sorensen (2004) provided criteria for identifiability and suggested putting constraints on some of the parameters, although automatic generation of interpretable and meaningful constraints remains difficult, especially in high-dimensional settings.

A second (and more fundamental) obstacle for the use of SEMs with genetic effects is that the underlying structure is often unknown. In such cases causal inference methods (Pearl 2009; Spirtes *et al.* 2001; Maathuis and Nandy 2016) can be used, that reconstruct causal models that are in some sense most compatible with the observed data. Most causal inference methods however require independent samples, and cannot account for genetic relatedness. For this reason, genotypic differences are most often modelled using a small number of QTLs with fixed effects (Neto *et al.* 2008, 2013; Scutari *et al.* 2014), but when part of the genetic variance is not explained by QTLs (missing heritability), the use of random genetic effects seems inevitable. Only a few works have studied reconstruction in the presence of such effects. Valente *et al.* (2010) and Töpner *et al.* (2017) proposed to perform causal inference after subtracting genomic predictions obtained from an MTM. Similarly, Gao and Cui (2015) applied the PC-algorithm (Spirtes *et al.* 2001) to the residuals of multiSNP models. The difficulty with these approaches is that the MTM is limited to small numbers of traits, and that the existence of direct genetic effects cannot be tested. For example, if the causal graph among three traits is *ϒ*_1_ → *ϒ*_2_ → *ϒ*_3_, and there are direct genetic effects on *ϒ*_1_ and *ϒ*_3_, then the absence of a direct genetic effect on *ϒ*_2_ cannot be inferred from MTM residuals.

Inspired by these problems we define a framework in which direct genetic effects are part of the causal graph, and a single node *G* represents all direct genetic effects. For each trait *ϒ*_*j*_ an arrow *G* → *ϒ_j_* is present if and only if the direct genetic effect on *ϒ*_*j*_ is nonzero, i.e., if the *j*th diagonal element of Σ_*G*_ is positive. See Figure 3 below for an example. Although our causal interpretation of genetic effects is not new (Stephens 2013; Valente *et al.* 2013, 2015), this work appears to be the first that formalizes it. In particular, we show that the Markov property holds for the graph extended with genetic effects (Theorem 1 below). Informally speaking, this means that there is a one-to-one correspondence between edges in the causal graph and conditional dependencies in the distribution of the traits and genetic effects. This means that edges (either between two traits, or between a trait and *G*) can in principle be inferred from multi-trait data. Consequently, while some of the covariances between direct genetic effects (contained in Σ_*G*_) may still be unidentifiable, we *can* identify which rows and columns in Σ_*G*_ are zero.

Based on the Markov-property we propose the PCgen algorithm. PCgen stands for PC with genetic effects, and is an adaptation of the general PC-algorithm (named after its inventors Peter Spirtes and Clark Glymour). Briefly, PCgen assesses the existence of a direct genetic effect on a given trait by testing whether its genetic variance is zero, conditional on various sets of other traits. For the existence of an edge between traits *ϒ*_1_ and *ϒ*_2_ we test whether in a bivariate MTM the residual covariance between *ϒ*_1_ and *ϒ*_2_ is zero, again conditional on sets of other traits. Under the usual assumptions of independent errors, recursiveness and faithfulness, we show that PCgen can recover the underlying partially directed graph (Theorem 2). Because fitting an MTM for all traits simultaneously is no longer necessary, PCgen can handle a considerably larger number of traits.

While our approach is generally applicable to any species and relatedness matrix, our implementation of PCgen is currently limited to the specific (but important) case of populations where observations on genetically identical replicates are available, assuming independent genetic effects (i.e., as in the classical estimation of broad-sense heritability). This is partly for pragmatic reasons (e.g., the lower computational requirements), and partly for statistical reasons. In particular, successful reconstruction requires sufficient power in the tests for direct genetic effects (*G* → *ϒ*_*j*_) and those for the between traits relations (*ϒ*_*j*_ → *ϒ*_*k*_). Given the availability of replicates, this power is likely to be highest when the original observations are used, instead of genotypic means and a marker based genetic relatedness matrix (GRM), modelling additive effects (Kruijer *et al.* 2015). Although mixed models with *both* replicates and a GRM may further increase power, the increase is often modest, and statistical inference can become biased under model misspecification (e.g., when the GRM models additive effects, and the true architecture is partly epistatic; see Kruijer (2016)). By contrast, using only replicates, unbiased estimation of broad-sense heritability is always possible, regardless of the population structure and genetic architecture. The downside is that the contributions of different types of genetic effects cannot be distinguished. On the positive side, PCgen appears to be the first algorithm that can infer the presence of direct genetic effects based on phenotypic data alone.

Our approach is related to that of Stephens (2013), who inferred the sets of traits being directly and indirectly affected by a given locus, assuming unrelated individuals and using only summary statistics. Here we instead consider sums of individual locus effects, for possibly related individuals. Moreover, PCgen also aims to reconstruct the causal structure *within* the sets of traits with direct and indirect genetic effects, and can deal with larger numbers of traits.

The paper is organized as follows. After introducing SEMs with genetic effects, we define their graphical structure, and from this perspective review existing approaches. We then describe the general form of the PCgen-algorithm for estimation of the graphical structure, followed by various proposals for the required conditional independence tests. Next we test PCgen performance in data simulated with a statistical as well as a crop-growth model, and analyze a maize and a rice dataset. Finally, we state several results regarding PCgen’s statistical properties. Table S1 provides an overview of the notation, and Appendix A.1 contains the necessary graph-theoretic definitions. Figure 2 provides a graphical summary of our theory and methodology.

**Figure. 2.**
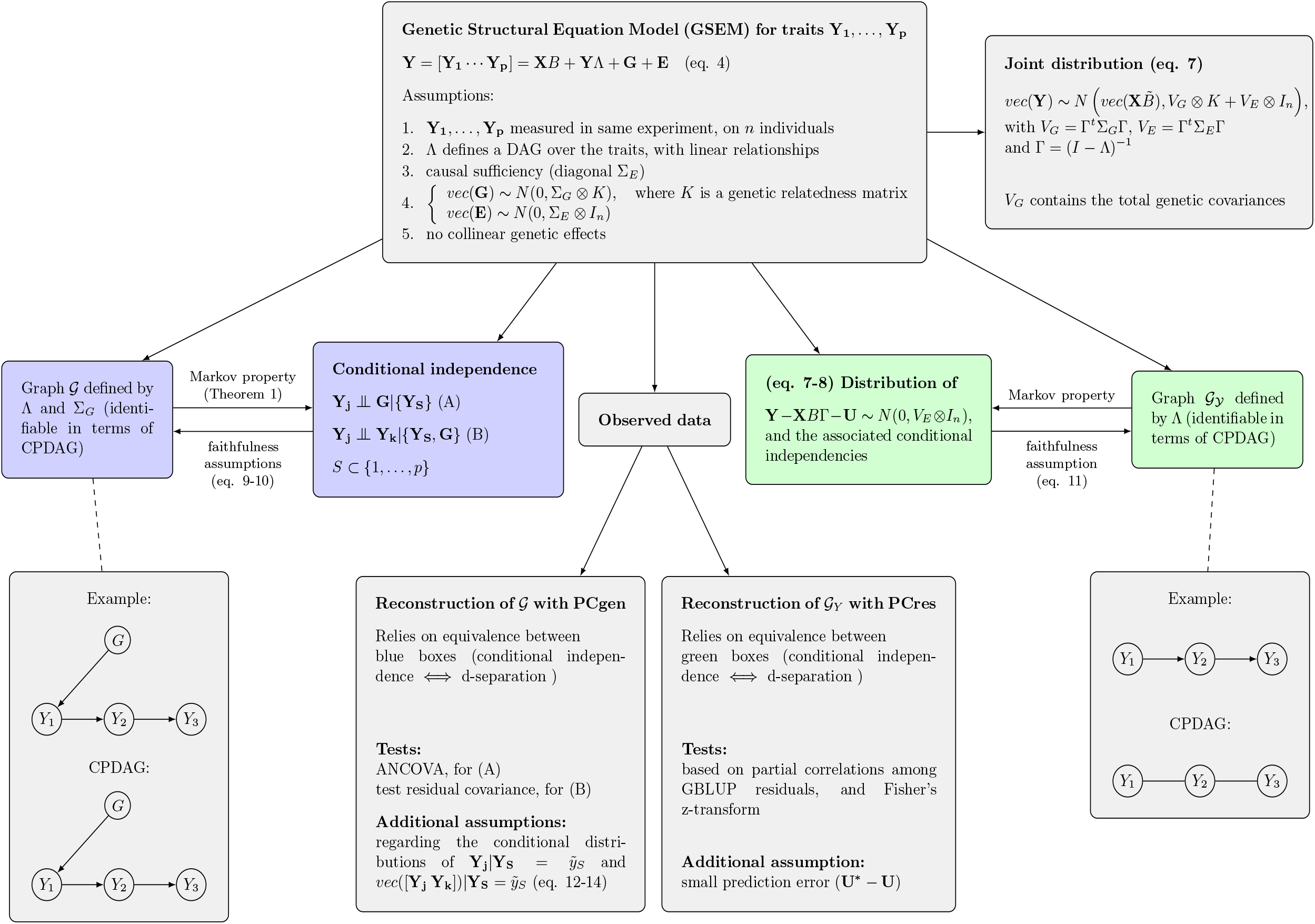
Graphical summary of the theory and methodology. The Markov property on the right (green; for the residuals) is well known from the literature, while the Markov property on the left (blue, for the conditional distributions of **ϒ**_**1**_, …, **ϒ**_**p**_, **G**) is established in Theorem 1. Table S1 contains an overview of the notation, and Appendix A.1 provides the necessary graph-theoretic definitions.

## Materials and methods

### Structural Equation Models

To introduce structural models, we first consider a simple linear SEM without genetic effects:

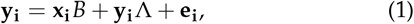

where **y_i_** is a 1 × *p* vector of phenotypic values for *p* traits measured on the *i*th individual, **e_i_** a vector of random errors, and Λ is a *p* × *p* matrix of structural coefficients. The *q* × *p* matrix *B* = [*β*^(1)^ … *β*^(*p*)^] contains intercepts and trait specific fixed effects of (exogenous) covariates, whose values are contained in the 1 ×*q* vector **x_i_**.

To write (1) in matrix-form, we define the *n* × *q* design matrix **X** with rows **x_i_**. Similarly we define *n* ×*p* matrices **ϒ**= [**ϒ**_**1**_ ··· **ϒ**_**p**_] and **E**= [**E**_**1**_ ··· **E**_**p**_], with rows **y**_**i**_ and **e**_**i**_, and columns **ϒ**_**j**_ and **E**_**j**_. We can then write

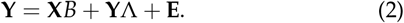

Λ has zeros on the diagonal and defines a directed graph 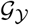 over the traits *ϒ*_1_, … , *ϒ*_*p*_, containing the edge *ϒ*_*j*_ → *ϒ_k_* if an only if the (*j*, *k*)*th* entry of Λ is nonzero. The columns in (2) correspond to *p* linear *structural equations*, one for each trait. These are determined by the *path coefficients*, the nonzero elements in Λ. For example, in Figure 1, if **X**= 1_*n*_ is the *n* × 1 vector of ones and *B* = [*μ*_1_ *μ*_2_ *μ*_3_], the third trait has values **ϒ**_**3**_ = *μ*_3_1_*n*_ + *λ*_13_ **ϒ**_**1**_ + *λ*_23_ **ϒ**_**2**_ + **E**_**3**_. The equality sign here can be understood as an assignment, i.e., **ϒ**_**3**_ is determined by the values of **ϒ_1_** and **ϒ_2_** (its *parents* in the graph 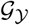) and an error. If the directed graph does not contain any cycle (i.e., a directed path from a trait to itself), it is a directed acyclic graph (*DAG*), and the SEM is said to be *recursive*. In the notation we will distinguish between the nodes *ϒ*_1_, … , *ϒ*_*p*_ in the graph 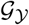 (normal type), and the random vectors **ϒ**_**1**_, … , **ϒ**_**p**_ that these nodes represent (bold face).

As mentioned above, SEMs can be used to predict the effects of *interventions*, which mathematically correspond to changes in the structural equations. For example, suppose that in Figure 1, *ϒ*_1_, *ϒ*_2_ and *ϒ*_3_ are the expression levels of three genes, and *ϒ*_4_ is plant height. Then after forcing *ϒ*_2_ to be zero (e.g., by knocking out the gene), the total effect of *ϒ*_1_ on *ϒ*_4_ changes from (*λ*_13_*λ*_34_ + *λ*_12_*λ*_23_*λ*_34_) to *λ*_13_*λ*_34_ (File S7.3 and File S7.4 provide other examples, involving genomic prediction). More generally, the new joint distribution of **ϒ**_**1**_, … , **ϒ**_**p**_ after an intervention can be obtained from the manipulation or truncated factorization theorem (Pearl 2009), *without* observations from the new distribution. For the consequences for genomic prediction, see Valente *et al.* (2013) and the Discussion below.

### GSEM: Structural Equation Models with genetic effects

Gianola and Sorensen (2004) extended model (1) with random genetic effects **g**_**i**_: for individuals *i* = 1, … , *n*, it is then assumed that

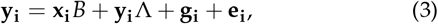

where the 1 × *p* vectors **g**_**i**_ contain the direct genetic effects for individuals *i* = 1, … , *n*. We will refer to model (3) as a linear GSEM (genetic structural equation model), or simply GSEM. While the genetic effects introduce relatedness between individuals, there is no form of social interaction (as in Moore *et al.* (1997), and Bijma (2014)). Each **g**_**i**_^*t*^ follows a *N*(0, Σ_*G*_) distribution, where Σ_*G*_ is a *p* × *p* matrix of genetic variances and covariances. The vectors **g**_**i**_ are independent of the **e**_**i**_’s, but not independent among themselves. Defining a *n* × *p* matrix **G** = [**G_1_** ··· **G_p_**] with rows **g**_**i**_ and columns **G**_**j**_ (*j* = 1, … , *p*), we can extend (2) as follows:

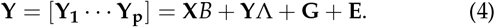

Each vector **G**_**j**_ is the vector of direct genetic effects on the *j*th trait. We make the following assumptions about the GSEM defined in (4):

1. *all traits are measured in the same experiment:* the rows **y**_**i**_ of **ϒ** may be either observations at plot or plant level or genotypic means across plots or plants, but the observations should always come from the same experiment. In addition, the residual errors originate from biological variation, i.e., measurement errors are negligible (this in contrast to related work on Mendelian Randomization (Hemani *et al.* 2017)).
2. *recursiveness:* the graph 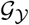 defined by Λ is a DAG. Consequently, there should be no feedback loops.
3. *causal sufficiency:* the covariance matrix Σ_*E*_ of the error vectors **e**_**i**_ is diagonal, i.e., there are no latent variables. This means that all nonzero (non-genetic) correlations between traits must be the consequence of causal relations between the traits. We also assume the diagonal elements of Σ_*E*_ to be strictly non-zero.
4. *Genetic relatedness among individuals:* **G** is independent from **E**, and has a matrix-variate normal distribution with row-covariance *K* and column covariance Σ_*G*_, where *K* is a *n* × *n* relatedness matrix, which we describe in more detail below (section ‘Genetic relatedness’). Equivalent to this, the *np* × 1 vector *vec*(**G**) = (**G**_**1**_^*t*^, … , **G**_**p**_^*t*^)^*t*^ is multivariate normal with covariance Σ_*G*_ ⭙ *K*, where *vec* denotes the operation of creating a column vector by stacking the columns of a matrix. Consequently, each **G**_**j**_ is multivariate normal with covariance 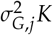, where the variances 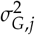 form the diagonal of Σ _*G*_. Using the same notation, we can write that **E** is matrix-variate normal with row-covariance *I_n_* and column covariance Σ_*E*_, and that *vec*(**E**) ~ *N*(0, Σ_*E*_ ⊗ *In*).
5. *No collinear genetic effects:* the diagonal elements of Σ_*G*_ do not need to be strictly positive, but for all nonzero elements the corresponding correlation should not be 1 or −1.

Assumptions 1-4 were also made in related work on structural models with random genetic effects (Valente *et al.* 2010; Töpner *et al.* 2017), and 1-3 are commonly made for structural models without such effects. Assumption 1 is implicit in the GSEM model (4) itself, as it is assumed that the structural equations propagate all errors to traits further down in the graph. Network reconstruction with traits from different experiments would rely completely on the genetic effects, requiring Σ_*G*_ to be diagonal, which is a rather unrealistic assumption (see the Discussion, section ‘Data from different experiments’). A small amount of measurement error does not seem to pose problems for our PCgen algorithm. Larger amounts of measurement error will decrease power, which can however be avoided by increasing the number of genotypes or replicates (see Table S4, discussed below). Assumption 1 does not require traits to be measured at the same time. In particular, it is possible to include the same trait measured at different time-points, which of course puts contraints on the causality. Such contraints can in principle be incorporated in our model, just as other biological contraints (see e.g., Peters *et al.* (2017)), although we will not explore this here. What is also implicit in the GSEM model (4) is that all causal relations between traits are linear. Our PCgen algorithm relies on this rather heavily, and we discuss the consequences of nonlinearity in the Results section below (specifically, in the APSIM simulations, and the example just before the Discussion). In specific cases it may be possible to obtain linearity by certain transformations of the data, but this requires prior knowledge that is typically unavailable. In the Discussion we suggest various directions of future work to deal with nonlinear relationships, as well as non-Gaussian errors. In any case, as long as the other assumptions hold, the core of our framework (the graphical representation of genetic effects with a single node *G*, and the Markov property in Theorem 1 below) is still valid for nonlinear GSEMs.

Assumption 2 (no cycles) is essential given the type of data considered here, as the reconstruction of feedback loops requires time-course data (Peters *et al.* 2017), typically with high-resolution. Without such data (or only a few time-points) it is impossible to verify this assumption, but Maathuis *et al.* (2010) provide examples of yeast data, where cycles are likely to exist, but structural models still outperform nonstructural models.

Assumption 3 (no latent variables) is important for the orientation of the edges, and has been studied in detail by many authors. In particular, Spirtes *et al.* (2001) and Colombo *et al.* (2012) proposed the FCI and RFCI algorithms, which are extensions of the PC-algorithm, and allow for latent variables. These algorithms could be extended with genetic effects, like we do here for the PC-algorithm (see the Discussion). Apart from non-linear trait-to-trait relations, the APSIM simulations below also contain latent variables.

As in related work (Valente *et al.* 2010; Töpner *et al.* 2017) as well as in much of the literature on multi-trait genomic prediction and GWAS (see e.g., Zhou and Stephens (2014) and Calus and Veerkamp (2011)), the relatedness matrix *K* is the same for all traits (Assumption 4). This may not hold if traits have very different genetic architectures, but seems a good approximation if most of the underlying QTLs are small. Large QTLs may be added as additional fixed effects.

Assumption 5 implies that for each pair of traits with direct genetic effects, these effects should not be the result of exactly the same set of QTLs, with exactly the same effect sizes. This seems a reasonable assumption whenever the underlying biological structures or processes are really different; see the section ‘Dealing with derived traits’ in the Discussion. Of course, reconstruction of direct genetic effects will be more difficult under strong correlations, similar to for example the reduced power in GWAS when two causal loci are in strong LD.

Finally, there are a few additional assumptions which are required for the PCgen-algorithm, and are not essential for the definition of GSEM; see the overview in Figure 2 and the discussion in section ‘Statistical properties of PCgen’ in the Results. In particular, we require the *faithfulness* assumptions defined by equations (9) and (10) below, and assumptions about the conditional distributions. Appendices A.5 and A.6 provide additional examples and results about faithfulness.

### Graphical representation of GSEM: extending 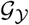 with genetic effects

In contrast to previous work, we will explicitly take into account the possibility that there are no direct genetic effects on some of the traits. In this case, the corresponding rows and columns in Σ_*G*_ are zero. Following the notation of Stephens (2013) (in the context of individual loci), we use *D* ⊆ {1, … , *p*} to denote the index set of the traits with direct genetic effects, and write Σ_*G*_ [*D*, *D*] for the submatrix with rows and columns restricted to *D*. From assumption 5 above, it follows that Σ_*G*_ [*D*, *D*] is non-singular, i.e., there can be no perfect correlations between direct genetic effects.

We graphically represent model (4) by a graph 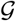 with nodes *ϒ*_1_, … , *ϒ*_*p*_ and a node *G*, which represent respectively **ϒ**_**1**_, … , **ϒ**_**p**_ and the matrix **G** = [**G**_**1**_ ··· **G**_**p**_]. 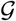 contains an edge *ϒ*_*j*_ → *ϒ*_*k*_ if the (*j*, *k*)th entry of Λ is nonzero, and an edge *G* → *ϒ*_*j*_ if **G**_**j**_ is nonzero with probability one, i.e., if 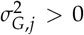. See Figure 3 for an example. In words, 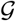 is defined as the original graph 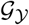 over the traits, extended with the node *G* and arrows *G* → *ϒ*_*j*_ for traits with a direct genetic effect, i.e., for all *j* ∊ *D*. Consequently, our main objective of reconstructing trait-to-trait relationships and direct genetic effects translates as reconstructing 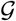.

**Figure. 3.**
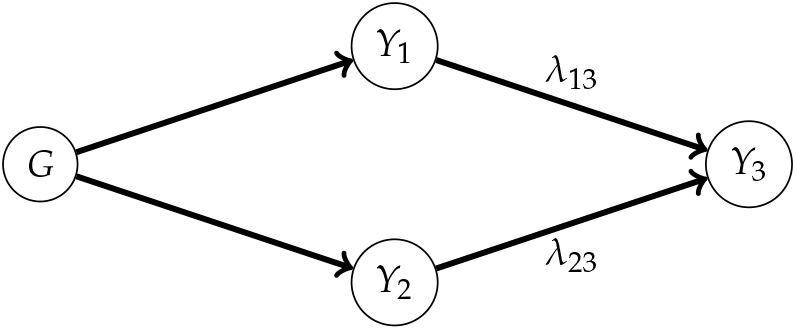
An example of a graph 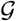 representing a genetic structural equation model (GSEM), with path-coefficients *λ*_13_ and *λ*_23_. There is no direct genetic effect on *ϒ*_3_, and therefore no edge *G* → *ϒ*_3_.

As for the **ϒ**_**j**_’s, we distinguish between the node *G* in the graph (normal type) and the random matrix **G** it represents (bold face). **G** is represented by a *single* node *G*, instead of multiple nodes *G*1, … , *Gp*. This choice is related to our assumption that *K* is the same for all traits; see File S7.1 for a motivating example. The orientation of any edge between *G* and *ϒ*_*j*_ is restricted to *G* → *ϒ*_*j*_, because the opposite orientation would be biologically nonsensical. Because of our assumption that 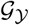 is a DAG, it follows that 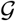 is a DAG as well, as a cycle would require at least one edge pointing into *G*.

We emphasize that 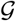 is just a mathematical object and not a complete visualization of all model terms and their distribution, as is common in the SEM-literature. In particular, 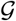 does not contain nodes for the residual errors, path coefficients, or information about the off-diagonal elements of Σ_*G*_. While in general^3^ Σ_*G*_ is not entirely identifiable (Gianola and Sorensen 2004), we will see that 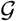 is identifiable in terms of its skeleton (the undirected graph obtained when removing the arrowheads) and some of the orientations. The skeleton is generally not equal to the conditional independence graph, which is the undirected graph associated with the inverse covariance or precision matrix (Spirtes *et al.* 2001; Kalisch and Bühlmann 2007). See File S6.2 for an example.

### Direct and indirect genetic effects

As pointed out by various authors (Gianola and Sorensen 2004; Valente *et al.* 2010, 2013; Töpner *et al.* 2017), the genetic variance of a trait is not only driven by its direct genetic effect (**g**_**i**_), but also by direct genetic effects on traits affecting it, i.e., its parents in the graph 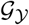. Assuming that the inverse Γ = (*I* − Λ)^−1^ exists ^4^, it follows from equation (3) that

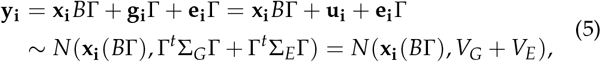

where the 1 × *p* vector **u**_**i**_ = **g**_**i**_Γ is the *total* genetic effect for the *i*th individual. The *n* × 1 vector **U**_**j**_ = **G***γ*_*j*_ contains the total genetic effects for the *j*th trait, where *γ*_*j*_ is defined as the *j*th column of Γ. The vector of *indirect* genetic effects is the difference **U**_**j**_ – **G**_**j**_. In Figure 3 for example ^5^, **G3**= (0, … , 0)^*t*^ and **U**_**3**_ = *λ*_13_**G_1_** + *λ*_23_**G**_**2**_.

Likewise, we can distinguish between the contribution of direct and indirect genetic effects to the genetic covariance. The (*j*, *k*)th element of *V_G_* = Γ^*t*^ Σ_*G*_ Γ in (5) is the *total* genetic covariance between **ϒ**_**j**_and **ϒ**_**k**_. This is what is usually meant with genetic covariance. Most often, this is different from the covariance between the direct genetic effects **G**_**j**_ and **G**_**k**_, given by Σ_*G*_ [*j*, *k*]. Indeed, Σ_*G*_ [*j*, *k*] affects the total genetic covariance, but the latter is also driven by causal relationships between traits, as defined by Γ = (*I* – Λ)^−1^. If no such relations exist, then Λ contains only zeros, and *V_G_* = Σ_*G*_. In general however these matrices are different, and depending on the structure of the graph and the path coeffecients, the correlation 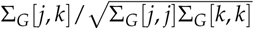 may be much larger than 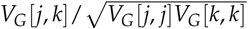, or vice-versa. For example, given direct effects **G_1_** and **G_2_** with equal variance and correlation 0.9, and an effect *ϒ*_1_ → *ϒ*_2_ of size −1, the total genetic correlation is −0.22. Regarding the diagonal of *V_G_*, we note that traits without a direct genetic effect may still have positive genetic variance.

### Genetic relatedness

The genetic relatedness matrix *K* introduced in assumption 4 determines the covariance between the rows of **G**. In principle our approach allows for any type of GRM, but for simplicity we focus on the following types. In all cases, *K* has dimension *n* × *n*.

- *K* = *ZZ^t^*, *Z* being the *n* × *m* incidence matrix assigning *n* = *mr* plants (or plots) to *m* genotypes, in a balanced design with *r* replicates for each genotype. This *K* is obtained when each genotype has an independent effect, as in the classical estimation of broad-sense heritability (or repeatability). Since no marker-information is included, the model cannot be directly used for genomic prediction, but we will see that for the reconstruction of 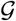 (using the training genotypes) it has considerable computational and statistical advantages.
- given only a single individual per genotype (or genotypic means) we assume *K* = *A*, *A* being a (*n* × *n*) GRM estimated from a dense set of markers, assuming additive infinitesimal effects.
- given both *r* replicates of *m* genotypes and a GRM *A* of dimension *m* × *m*, we assume that *K* = *ZAZ^t^*. In absence of non-additive effects, this covariance structure uses all available information. However, for computational reasons it is usually easier to work with either the replicates or with genotypic means and the GRM *A*. We further explore this issue in the simulations below and in the Discussion.

The balance required when *K* = *ZZ^t^* is necessary in Theorems 5 and 6 below, but is not a general requirement for our models, nor for the PCgen algorithm.

### The joint distribution implied by the GSEM

The sum **G** + **E** does in general not have a matrix-variate normal distribution, but from our assumption 4 it still follows that *vec*(**G** + **E**) is multivariate normal with covariance Σ_*G*_ ⊗ *K* + Σ_*E*_ ⊗ *In*. We can therefore rewrite equation (4) as

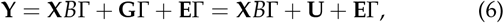

where **U** = **G**Γ is the *n × p* matrix of total genetic effects, with columns **U_j_**. Equation (5) now generalizes to

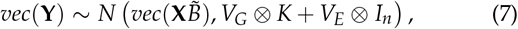

where 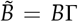 is the matrix of fixed effects transformed by Γ. This is a common model for multi-trait GWAS and genomic prediction (see among others Stephens (2013), Zhou and Stephens (2014) and Korte *et al.* (2012)). In those applications, however, *V_G_* and *V_E_* are usually arbitrary covariance matrices, whereas here they are modelled as functions of Σ_*G*_, Σ_*E*_ and Γ = (*I* Λ)^−1^.

Under Assumption 3 (Σ_*E*_ diagonal), Σ_*G*_, Σ_*E*_ and Λ together have at most *p*(*p* + 1)/2 + *p* + *p*(*p* − 1)/2 = *p*(*p* + 1) parameters, as many as *V_G_* and *V_E_* together. This suggests that Σ_*G*_, Σ_*E*_ and Λ might be identifiable from the distribution (7). In Appendix A.2 we show how Σ_*G*_, Σ_*E*_ and Λ can be obtained from *V_G_* and *V_E_*. Apart from Assumption 3, this requires knowledge of the graph, and the faithfulness assumptions (9)-(10) given below. Expressions (17)-(18) in Appendix A.2 can in principle be used to derive estimates 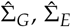 and 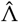 from estimates 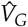 and 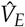, although the development of good estimators of Σ_*G*_, Σ_*E*_ and Λ is beyond the scope of this work. Such estimators should account for the structure of the GSEM, as defined by Λ and Σ_*G*_.

Using the results of Spirtes *et al.* (2001) (p. 371), it turns out that Γ can be written directly in terms of sums of products of path coefficients (see Appendix A.3). Consequently, there is no need to invert (*I* − Λ), although it still holds that Γ = (*I* − Λ)^−1^, provided the inverse exists. Recalling that *γ_j_* is the *j*th column of Γ, we can express the *j*th trait as

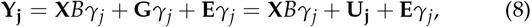

which is equation (6) restricted to the *j*th column. Similarly, for any non-empty index-set *S* ⊂ {1, … , *p*}, the *n* × |*S*| matrix **ϒ_S_** of traits in *S* (i.e., **ϒ** with columns restricted to *S*) equals **X***B*Γ_*S*_ + **G**Γ_*S*_ + **E**Γ_*S*_, where Γ_*S*_ is the *p* × |*S*| matrix with columns *γ_j_* (*j* ∈ *S*). Equation (26) (Appendix A.7) provides an expression for the covariance of *vec*(**ϒ_S_**). For the corresponding nodes in the graph, we write *ϒ_S_* = {*ϒ_m_* : *m* ∈ *S*}.

### Causal inference without genetic effects

So far we have assumed that 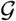 is known: given sufficient restrictions on Λ, Σ_*G*_ and Σ_*E*_, it may then be possible to estimate these matrices (Gianola and Sorensen 2004). In this work however, we aim to reconstruct an unknown 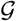, based on observations from a GSEM of the form (4). We will do this with the PCgen algorithm introduced below, but first briefly review the necessary concepts, as well as existing methods. Appendix A.1 contains a more detailed introduction.

Suppose for the moment we have observations generated by an acyclic SEM without latent variables, and without genetic effects. From the pioneering work of Judea Pearl and others in the 1980s it is known that we can recover the skeleton of the DAG and some of the orientations, i.e., those given by the *v*-structures. A *v*-structure is any triple of nodes *ϒ_j_*, *ϒ_k_*, *ϒ_l_* such that *ϒ_j_* → *ϒ_5_* ← *ϒ_l_*, without an edge between *ϒ_j_* and *ϒ_l_*. All DAGs with the same skeleton and *v*-structures form an *equivalence class*, which can be represented by a completed partially directed acyclic graph (CPDAG). DAGs from the same equivalence class cannot be distinguished using observational data, at least not under the assumptions we make here. For the reconstruction of the CPDAG, constraint-based and score-based methods have been developed (for an overview, see Peters *et al.* (2017)).

Here we focus on constraint-based methods, which rely on the equivalence of conditional independence (a property of the distribution) and directed separation (d-separation; a property of the graph). An important result is that an edge *ϒ_j_* − *ϒ_k_* is missing in the skeleton of the DAG if and only if *ϒ_j_* and *ϒ_k_* are d-separated by at least one (possibly empty) set of nodes *ϒ_S_*. Such *ϒ_S_* is called a *separating set* for *ϒ_j_* and *ϒ_k_*. Given the equivalence of d-separation and conditional independence, this means that we can infer the presence of the edge *ϒ_j_* − *ϒ_k_* in the skeleton by testing **ϒ_j_** ᚇ **ϒ_k_**|**ϒ_S_** for all **ϒ_S_**. The PC- and related algorithms therefore start with a fully connected undirected graph, and remove the edge *ϒ_j_* − *ϒ_k_* whenever **ϒ_j_** and **ϒ_k_** are found to be conditionally independent for some **ϒ_S_**. While the first constraint-based algorithms such as IC (Pearl 2014) exhaustively tested all possible subsets, the PC-algorithm (Spirtes *et al.* 2001) can often greatly reduce the number of subsets to be considered. Although this is not essential for the equivalence of d-separation and conditional independence, most constraint-based algorithms assume that observations be indendently and identically distributed, and structural equations with additional random effects are usually not considered.

### Existing approaches for estimating 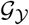, given genetic effects

To deal with the dependence introduced by the genetic effects, Valente *et al.* (2010) and Töpner *et al.* (2017) proposed to predict the total genetic effects (i.e., the term **U** in (6)), and perform causal inference on the residuals. These methods are flexible, in the sense that any genomic prediction method can be used, and combined with any causal inference method. A disadvantage however is that the presence of direct genetic effects cannot be tested. Suppose for example that *G* → *ϒ*_1_ → *ϒ*_2_ → *ϒ*_3_, and we subtract the total genetic effects. Then given only the residuals, we can never know if part of the genetic variance of *ϒ*_2_ was due to a direct effect *G* → *ϒ*_2_. Another disadvantage is that fewer of the between-trait edges can be oriented. Technically this is because in the CPDAG (showing which orientations can be recovered from data), typically more edges are undirected; see Appendix A.1 for more details. In the preceding example, the CPDAG associated with 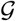 is *G* → *ϒ*_1_ → *ϒ*_2_ → *ϒ*_3_, i.e., all orientations can be recovered (see also Figure 2). By contrast, the CPDAG associated with 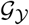 is *ϒ*_1_ − *ϒ*_2_ − *ϒ*_3_, and we only know that the orientation is *not ϒ*_1_ → *ϒ*_2_ ← *ϒ*_3_.

To use the causal information associated with the genetic effects, Töpner *et al.* (2017) estimated ‘genomic networks’, based on the predictions themselves. These however seem to require additional assumptions, which are not required for the residual networks (in particular, diagonal Σ_*G*_). Moreover, it seems difficult to relate edges in such a network to direct genetic effects (see the section ‘Data from different experiments’ in the Discussion, and File S7.2). In summary, residual and genomic networks only estimate the (CPDAG associated with the) subgraph 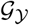 of trait to trait relations, instead of the complete graph 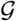.

Another disadvantage is that without specific models putting restrictions on *V_G_* and *V_E_*, the MTM (7) can only be fitted for a handful of traits (Zhou and Stephens 2014) for statistical as well as computational reasons. For example, Zwiernik *et al.* (2014) showed that for general Gaussian covariance models, (residual) ML-estimation behaves like a convex optimization problem only when *n* ≳ 14*p*. Similar problems are likely to occur for Bayesian approaches. The problem with fitting the MTM to data from the GSEM model (4) is that one cannot exploit the possible sparseness of *G*. Even for sparse graphs with few direct genetic effects, the matrices *V_G_* = Γ^*t*^ Σ_*G*_ Γ and *V_E_* = Γ^*t*^ Σ_*E*_ Γ may still be dense, requiring a total of *p*(*p* + 1) parameters. To overcome these limitations, we explicitly consider the presence or absence of direct genetic effects to be part of the causal structure, and develop PCgen, a causal inference approach directly on 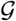.

### The PCgen algorithm

The main idea behind PCgen is that the PC-algorithm is applicable to any system in which d-separation and conditional independence are equivalent, and where conditional independence can be tested. We first describe the algorithm and propose the various independence tests; the equivalence is addressed in Theorem 1 below. If we define **ϒ_p+1_**:= **G** and temporarily rename the node *G* as *ϒ*_*p*+1_, PCgen is essentially the PC-algorithm applied to *ϒ*_1_, … , *ϒ*_*p*+1_:

1. **skeleton-stage.** Start with the fully connected undirected graph over {*ϒ*_1_, … , *ϒ*_*p*+1_}, and an empty list of separation sets. Then test the conditional independence between all pairs **ϒ_j_** and **ϒ_k_**, given subsets of other variables **ϒ_S_**. Whenever a p-value is larger than the pre-specified significance threshold *α*, update the skeleton by removing the edge *ϒ_j_* − *ϒ_k_*, and add *ϒ_S_* to the list of separation sets for *ϒ_j_* and *ϒ_k_*. This is done for conditioning sets of increasing size, starting with the empty set (*S* = ∅; marginal independence between **ϒ_j_** and **ϒ_k_**). Only consider *S* that, in the current skeleton, are adjacent to *ϒ_j_* or *ϒ_k_*.
2. **orientation-stage.** Apply the orientation rules given in File S1 (R1-R3 in Algorithm 1) to the skeleton and separating sets found in the skeleton-stage. For example, if the skeleton is *ϒ*_1_ − *ϒ*_2_ − *ϒ*_3_ and {*ϒ*_2_} is *not* a separating set for *ϒ*_1_ and *ϒ*_3_, the skeleton is oriented *ϒ*_1_ → *ϒ*_2_ ← *ϒ*_3_; otherwise, none of the two edges can be oriented.

In order to obtain PCgen, we need to make a few refinements to these steps. First, in the skeleton stage we need to specify *how* to test conditional independence statements. Clearly, independence between two traits requires a different test than independence between a trait (**ϒ_j_**) and **G**(i.e., **ϒ_p+1_**), in particular because the latter is not directly observed. Second, we need to modify the orientation rules, in order to avoid edges pointing into *G*. The usual rules give the correct orientations when given perfect conditional independence information, but statistical errors in the tests may lead to edges of the form *ϒ_j_* → *G*. Third, statistical errors can also make the output of PC(gen) order-dependent, i.e., putting the columns (traits) in a different order may lead to a different reconstruction. We therefore adopt the PC-stable algorithm of Colombo and Maathuis (2014), who proposed to perform all operations in skeleton- and orientation-stage list-wise (details given in File S1). Apart from eliminating the order-dependence, this has the advantage that all conditional independence tests of a given size |*S*| = *s* can be performed in parallel.

In summary, PCgen is the PC-stable algorithm with: (1) specific conditional independence tests (described shortly below) and (2) modified orientation rules, in order to avoid edges pointing into *G* (File S1.2). As in the original PC-algorithm, the number of type-I and type-II errors occurring in the tests is determined by the choice of the significance threshold *α*, which is discussed in section ‘Assessing uncertainty’ below and the Discussion.

### Skeleton stage: conditional independence tests

We can distinguish between the following types of conditional independence statements in the skeleton stage:

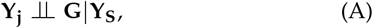

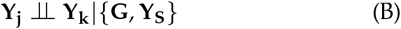

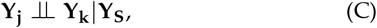

where *j*, *k* ∈ {1, … , *p*} (*j* ≠ *k*) and *S* ⊆ 1, … , *p j*, *k* (or *S* ⊆ 1, … , *p j*, in statement (A)). In words, (A) means that the trait **ϒ_j_** is independent of all genetic effects (**G**), conditional on the traits **ϒ_m_**(*m ∈ S*). If *S* is the empty set, this is understood as marginal independence of **ϒ_j_** and **G**. Similarly, (B) and (C) express conditional independence of traits **ϒ_j_** and **ϒ_k_** given **G** and **ϒ_S_**, or given **ϒ_S_** alone.

We now propose statistical tests for statements (A) and (B), which rely on the linearity of our GSEM as well as some additional assumptions, which we discuss in more detail below (section ‘Statistical properties of PCgen’, and Figure 2). Statement (C) can be tested using standard partial correlations and Fisher’s z-transform. However, as we show in File S6, this test is redundant, since for any set *ϒ_S_* that d-separates *ϒ_j_* and *ϒ_k_*, the set *ϒ_S_* ∪ {*G*} will also d-separate them. We therefore skip any test for **ϒ_j_** ╨ **ϒ_k_**|**ϒ_S_**, and instead test the corresponding statement including **G**, i.e., **ϒ_j_** ╨ **ϒ_k_**|{**ϒ_S_**, **G**}.

### Testing ϒ_j_ ⫫ G|ϒ_S_

Our test for statement (A) is based on the intuition that **ϒ_j_** is independent of **G**= [**G_1_** ··· **G_p_**] given **ϒ_S_**, whenever there is no direct genetic effect on **ϒ_j_** (i.e., **G_j_**= **0**), and all directed paths from *G* to *ϒ_j_* are blocked by the set *ϒ_S_* = {*ϒ_m_* : *m* ∈ *S*}. In particular, if *S* is the empty set, there should not be any directed path from *G* to *ϒ_j_*. Because directed paths from *G* to *ϒ_j_* will generally introduce some genetic variance in **ϒ_j_**, the idea is to test whether there is significant genetic variance in the conditional distribution of **ϒ_j_** given 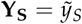. This is done as follows:

- When *K* = *ZZ^t^*, we use the classical F-test in a 1-way ANOVA, with **X** and 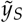 as covariates. Technically, this is an ANCOVA (analysis of covariance), where the treatment factor genotype is tested conditional on the covariates being in the model.
- For other *K* one can use a likelihood ratio test (LRT). The asymptotic distribution under the null-hypothesis is a mixture of a point mass at zero and a chi-square.

In both cases, it is assumed that the conditional distribution of **ϒ_j_** given 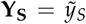 is that of a single trait mixed model, the mean being a linear regression over the conditioning traits. This assumption is made mathematically precise below in equations (12) and (14).

### Testing ϒ_j_ ╨ ϒ_k_ | {G, ϒ_S_}

For statement (B) we mostly use the residual covariance (RC) test, which is based on the conditional distribution of *vec*([**ϒ_j_ ϒ_k_**]) given the observed 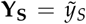. It is assumed that this distribution is that of a bivariate MTM, again with the mean being a linear regression over the conditioning traits; see equations (13) and (14) below. Assuming the bivariate MTM, we test whether the residual covariance^6^ is zero, using the LRT described in File S1.3. The underlying idea is that a nonzero residual covariance must be the consequence of an edge *ϒ_j_* → *ϒ_k_* or *ϒ_k_* → *ϒ_j_*, because of the assumed normality and causal sufficiency. On the other hand, a nonzero *genetic* covariance may also be due to covariance between direct genetic effects on these variables, or due to a genetic effect on a common ancestor. The RC-test therefore compares the full bivariate mixed model with the submodel with diagonal residual covariance, while accounting for all genetic (co)variances. The RC-test is not to be confused with a test for zero *genetic* covariance. The latter is often useful for data exploration, but has no role in PCgen (although in File S1.4 we describe a LRT test, which is implemented in our software).

An alternative to the RC-test is the RG-test (*R*esiduals of *G*BLUP). Fitting the MTM (7), we obtain the BLUP **U**^∗^ of the total genetic effects **U**= **G**Γ, and the BLUE 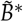 of the fixed effects. We then test the significance of partial correlations among residuals, i.e., the columns of 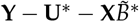. When **U**^∗^ is close enough to **U**, it follows from (6)-(7) that the covariance of *vec*(**ϒ** − **U**^∗^) is approximately (Γ^*t*^Σ_*E*_Γ) ⊗ *I_n_*, i.e., that of independent samples, without any genetic relatedness. This approach is very similar to the work of Valente *et al.* (2010) and Töpner *et al.* (2017), who instead took a fully Bayesian approach to predict **U**. In either case, the performance of the RG-test critically depends on the prediction error (**U**^∗^ − **U**). As mentioned before, fitting an MTM is usually challenging for more than 5-10 traits; we therefore also consider residuals of single trait GBLUP, as an approximation.

### PCres: reconstructing only trait-to-trait relationships

Testing only conditional independencies of the form (B), one can reconstruct the graph 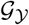 of trait-to-trait relations (see the green boxes in Figure 2). Moreover, if this is done with the RG-test, the algorithm is very similar to the residual approaches of Valente *et al.* (2010) and Töpner *et al.* (2017). Staying within the context of the PC-algorithm and using residuals from GBLUP, we will call this PCres. As for the RG-test in PCgen, PCres can be based on residuals of either single or multi-trait mixed models.

### Software

In our R-package pcgen we implemented PCgen for the case *K* = *ZZ^t^*. PCres is implemented for *K* = *ZZ^t^*, *K* = *A* as well as *K* = *ZAZ^t^*. Moreover, PCres can be based on either residuals of the full MTM (7) (only for small numbers of traits), or from univariate models (the default). Tables 2 and 3 in File S2 provide a complete overview of the options, with the required R-commands. The package is freely available at https://cran.r-project.org/web/packages/PCgen/index.html. pcgen is built on the pcalg package (Kalisch *et al.* 2012; Hauser and Bühlmann 2012), in which we modified the orientation rules and the default conditional independence test.

### Assessing uncertainty

The PC-algorithm is asymptotically correct, in the sense that the underlying CPDAG is recovered if conditional independence can be tested without error (Spirtes *et al.* 2001). In Theorem 2 below we provide a similar consistency result for PCgen. In practice however, type-I or type-II errors are likely to occur, leading to incorrect edges in the graph. Depending on the significance level *α* used in each test, there may be more type-I errors (large *α*) or rather more type-II errors (small *α*). Reliable control of the (expected) false positive rate or total number of false positives remains challenging; see the Discussion (’Assessing uncertainty’). We will therefore just consider the p-values as they are, and analyze the real datasets for different significance thresholds. Following Kalisch and Bühlmann (2007) and Kalisch *et al.* (2012), we report, for each remaining edge, the largest p-value found across all conditioning sets for which the edge was tested.

### Extensions of PCgen

File S3 describes several extensions of PCgen, which are partly implemented in our software. Among others, the causal graph and PCgen could be extended with fixed effect QTLs, and PCgen can be speed up by starting with a skeleton obtained from PCres (’prior screening’). As in the pcalg-package, it is possible to restrict the maximum size of the conditioning sets, also to speed up computation.

### Data availability

The maize and rice data used below can be accessed at respectively https://doi.org/10.15454/IASSTN and https://doi.org/10.6084/m9.figshare.7964357.v1.

## Results

### Simulations with randomly drawn graphs

To compare the different algorithms we simulated random GSEMs, by randomly sampling the sets *D* (defining the traits with direct genetic effects) and the covariance matrices Σ_*G*_, combined with randomly drawn DAGs over the traits 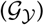. Traits were simulated for an existing population of 256 maize hybrids (Millet *et al.* 2016). Two replicates of each genotype were simulated. Given the 256 × 256 additive relatedness matrix *A* based on 50k SNPs, genetic effects were simulated such that *vec*(**G**) ~ Σ*G* ⊗ (*ZAZ^t^*) (i.e., the vector of genetic effects for all the replicates, and all traits). File S4.1 provides further details, such as the magnitude of genetic (co)variances. We focus here on the comparison of

- PCgen based on the replicates, assuming *K* = *ZZ^t^* (i.e., ignoring *A*). By default, we apply the prior-screening with PCres.
- PCres (replicates): PCres based on residuals from univariate GBLUP, again using only the replicates.
- PCres (means): PCres based on residuals from multivariate GBLUP, using genotypic means and the relatedness matrix *A* that was used to simulate the data.

Table S2 provides results for variations on these algorithms, including PCgen without prior screening. In all simulations the significance threshold was *α* = 0.01. The effect of sample size and the trade-off between power and false positives as function of *α* was already investigated by Kalisch and Bühlmann (2007) for the standard PC-algorithm, and is likely to be similar for PCgen.

We separately evaluated the reconstruction of 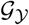 and the edges *G* → *ϒ_j_*, as the latter is only possible with PCgen. To assess the difference between estimated and true skeleton of 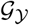, we considered the true positive rate (TPR), the true discovery rate (TDR) and the false positive rate (FPR). Additionally we used the Structural Hamming Distance (SHD), which also takes into account the orientation of the edges. File S4.2 provides definitions of these criteria. Reconstruction of *G* → *ϒ_j_* is only assessed in terms of TPR, TDR and FPR, as these edges can have only one orientation.

#### Simulation results

We first performed simulations with *p* = 4 traits (scenario 1), with each potential edge between traits occurring in the true graph with probability *p_t_* = 1/3. Hence, for any given trait, the expected number of adjacent traits was (*p* − 1)*p_t_* = 1. The edges *G* → *ϒ_j_* were included in the true graph with probability *p_g_* = 1/2. In a related set of simulations (scenario 2), *p_t_* was increased to 0.5, giving denser graphs. In both scenarios, PCgen reconstructed the edges *G* → *ϒ_j_* with little error, the average TPR being above 0.97 and FPR around 0.03 (Table 1). In the first scenario, about a third of the actual edges between traits was not detected with PCgen (TPR ≈ 0.65, i.e., the proportion of true edges that was discovered). At the same time the number of false edges in 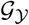 was very low, which is also reflected in high TDR values (the proportion of edges in the reconstruction that is true). In scenario 2, the TPR, FPR and TDR all increased. Hence, for denser graphs, more of the true edges were found, at the expense of a somewhat higher number of false edges.

**Table 1.**
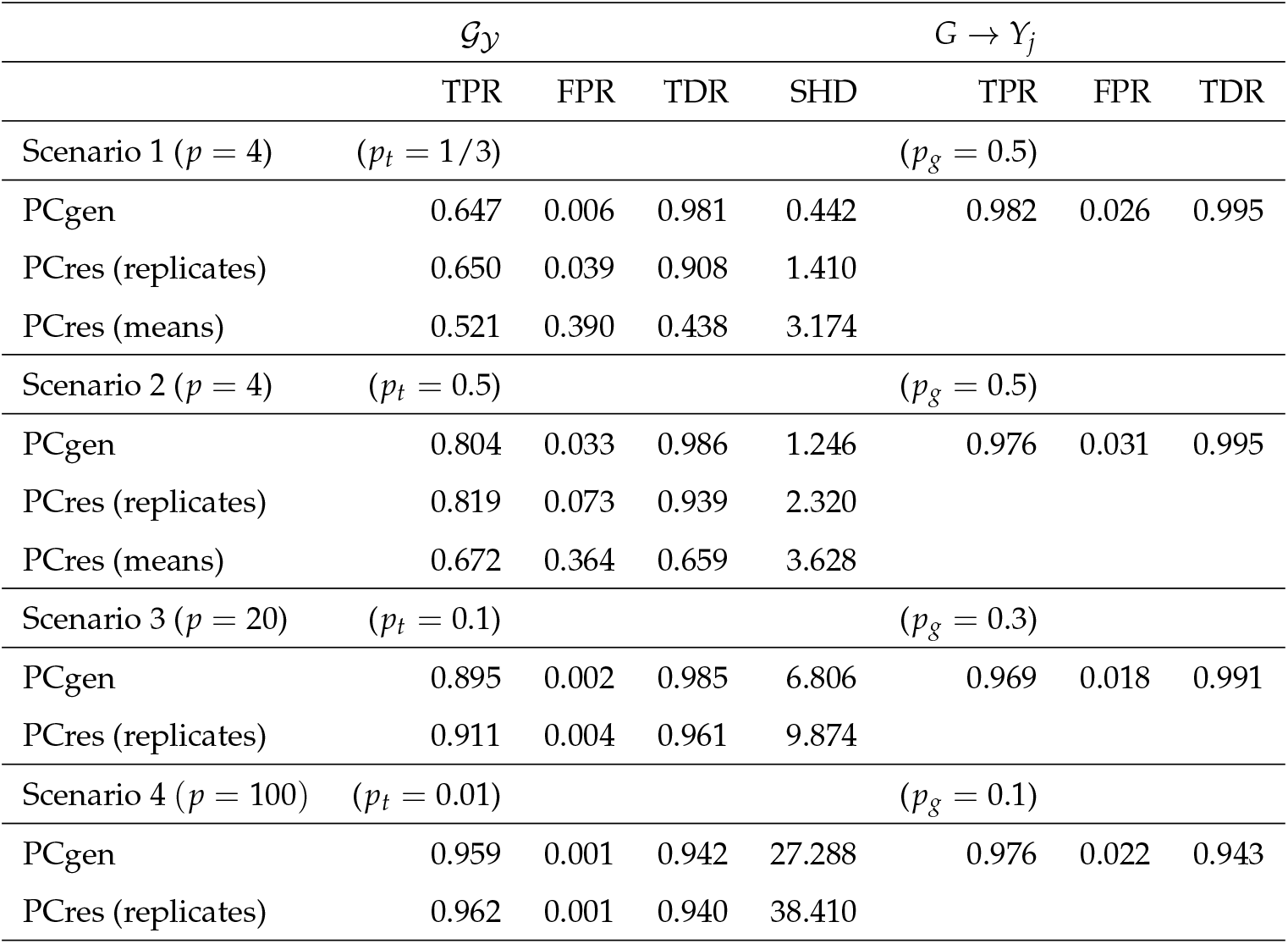
Performance of PCgen and residuals-based approaches, averaged over 500 simulated datasets per scenario. Standard errors for the TPR, FPR and TDR were between 0.001 and 0.015. Standard errors for the SHD were around 0.06 (scenarios 1 and 2), 0.18 (scenario 3) and 0.28 (scenario 4). For the performance of other variants of PCgen and PCres in scenarios 1 and 2, see Table S2. In scenario 4, we used PCgen with the RG-test (PCgen-RG-uni); in the other scenarios we used the RC-test, with prior screening (PCgen-RC-screening). All acronyms are explained in Table 2 in File S2. PCres (replicates) and PCres (means) refer to PCres-uni-R and PC-multi-A.

PCres (replicates) outperformed PCres (means), in spite of the use of univariate GBLUP, and ignoring the actual relatedness matrix. Hence, the information contained in the replicates appears much more important than the precise form of the relatedness matrix, or unbiased estimation of genetic correlations. The performance of PCres strongly depends on the prediction error of the GBLUP, and, in line with the results of Kruijer *et al.* (2015), this error appeared lowest when using the replicates. The use of both the replicates and the marker-based GRM (i.e., assuming *K* = *ZAZ^t^*, as the data were generated), further improved performance, but only slightly (Table S1, ‘PCres-uni-RA’). Unsurprisingly, the MTM required for PCres (means) was computationally more demanding, and often not estimable for more than 4 traits. Motivated by this computational advantage and the statistical advantages mentioned in the Discussion, all analyses in the remainder will only consider PCgen and PCres based on replicates.

For the trait-to-trait relations, PCgen and PCres (replicates) had very similar performance in terms of TPR, TDR and FPR. However, PCgen substantially improved the orientation of these edges, as shown by the reduced SHD. This is not because the algorithm as such is better, but rather because more of the edges can be oriented from conditional independence tests, which in turn is a consequence of the addition of the edges *G* → *ϒ_j_* to the graph over the traits (see again the example in Figure 2).

To assess performance in higher dimensions, we simulated data sets with *p* = 20 traits, *p_g_* = 0.3 and *p_t_* = 0.1 (scenario 3) and with *p* = 100, *p_g_* = 0.1 and *p_t_* = 0.01 (scenario 4). Both scenarios consider sparse graphs; denser graphs can be analyzed as well, but, for *p* larger than 20-30, require several hours or even days, unless the size of the conditioning sets is restricted or PCgen would be parallelized. Here we limited the size of conditioning sets to 3 (scenario 3) and 2 (scenario 4). As in the first two scenarios, PCgen achieved a strong reduction in SHD, and reliable reconstruction of the direct genetic effects (Table 1).

To assess the effect of thresholding the size of conditioning sets, we simulated 200 datasets with *p* = 10 traits and a relatively dense graph (*p_g_* = 0.4 and *p_t_* = 4/9), and used PCgen with various thresholds (Table S3). The restricted maximum size means that a certain number of conditional independence tests is skipped, which may lead to extra false positives. However, the thresholding is only done in PCgen itself and not in the prior screening with PCres (which is much faster, and already removes most false edges). Consequently, thresholding had very little effect on the reconstruction of trait-to-trait relations 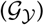, but did lead to a higher FPR in the reconstruction of the direct genetic effects (0.07 without thresholding, 0.08 with *m* = 3, and 0.48 with *m* = 1). Also the accuracy in the orientations of 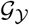 slightly decreased (the SHD increasing from 15.9 to 16.2).

In another set of simulations, we explored the effect of measurement error. As expected, increasing amounts of measurement error decreased the power to detect between-trait edges as well as direct genetic effects (Table S4). However, the loss in power could largely be compensated by increasing the number of replicates, or the number of genotypes. The latter was most effective for the between-trait edges, while increased replication gave the highest power for the edges *G* → *ϒ_j_*.

In our final set of simulations (Table S5) we explored the effect of strong correlations in Σ_*G*_, i.e., when Assumption 5 is close to being violated. We simulated an example with two traits whose direct genetic effects had unit variance, and increasing covariance (0, 0.5 and 0.95). The corresponding TPR values for the genetic effects were respectively 0.94, 0.85 and 0.60. Consequently, even in the presence of strong correlations, PCgen still had some power to detect direct genetic effects.

### Simulations using a crop-growth model

We also simulated data using the popular crop growth model APSIM-wheat (Keating *et al.* 2003; Holzworth *et al.* 2014). Compared to the preceding simulations this represents a more challenging scenario, as several of the underlying assumptions are violated. In particular, the data-generating process introduces nonlinearities and latent variables. We simulated 12 traits for an existing wheat population of 199 genotypes, with three replicates each. The traits include seven primary traits, four secondary traits and yield (*ϒ*). File S5 provides further details, and trait acronyms are given in Table S6. Traits were simulated by running a discrete dynamic model from the beginning (*t* = 0) to the end of the growing season (*t* = *T*). Motivated by the fact that some trait measurements are destructive, observations are only taken at *t* = *T*. Figure S1 (A) shows the summary graph ^7^, defining the causal effects from one time-step to the next (Peters *et al.* 2017). There are direct genetic effects on all of the primary traits, which have heritability 0.9. The genetic effects originate from 300 trait-specific QTLs, with randomly drawn effect sizes. There are no direct genetic effects on the secondary traits and yield.

Compared to the simulations above, it turned out to be much harder to detect the absence of direct genetic effects: in the PCgen reconstruction, all 12 traits had such effects (Figure S1 (B); highest p-value: 1.7 · 10^−4^). These false positives seemed to be a consequence of the nonlinearities in the data-generating distribution, which are not accounted for in our tests. The reconstructed trait-to-trait relations were mostly correct, except for the missing edge *GN → ϒ*, and one incorrect orientation (*ϒ → GW*). PCres made the same errors (Figure S1 (C)), with an additional false arrow (*MGS − SP*). The standard PC-stable algorithm applied to all traits and QTLs led to many more errors (Figure S1 (D)), such as the false edge between *GW* and *RUE*, the missing edge *TFI → FT* and some incorrect orientations. These errors occurred because for various traits **ϒ_j_** many QTL-effects were removed from the graph, i.e., for some set of traits *ϒ_S_*, the conditional independence **ϒ_j_** ⫫ *QTL*|**ϒ_S_** was mistakenly accepted. This in turn led to problems in the remaining tests, where part of the genetic variance was not taken into account. We emphasize that all 300 QTLs were available to the PC-algorithm, and no other markers were provided. Hence, the poor performance in this case is really a consequence of the small effects, rather than the difficulty of QTL *detection*.

### Two case-studies

We now use PCgen to analyze real data from four field trials and one experiment in a phenotyping platform. In all network reconstructions we used a significance threshold of *α* = 0.01. Reconstructions with *α* = 0.001 are shown in Figures S2 and S4. Table S9 and Figure S5 contain p-values for the remaining edges. In all datasets we removed traits that were derived as sums or ratios of other traits, rather than being directly measured. In particular, the maize data do not contain grain number, which was defined as the ratio of yield over grain weight. We return to this issue in the Discussion.

#### Maize

First we analyze the field trials described in Millet *et al.* (2016) and Millet *et al.* (2019), involving 254 hybrids of maize (*Zea mays*). We consider a subset of four trials, representing four (out of a total of five) different environmental scenarios described in Millet *et al.* (2016). See Table S8 for an overview. The scenarios were derived from physiological knowledge, crop-growth models and environmental sensors in the fields. Scenarios were defined as a combination of well-watered or water-deficient conditions (WW versus WD) and temperature. The latter was classified as “Cool” (average maximum and night temperature below respectively 33 and 20 degrees Celcius), “Hot” (above 33 and 20 degrees) or “Hot(days)” (max. temperature above 33, night temperature below 20). Most trials included 7 traits:

- three height traits, i.e., tassel height (*TH*), ear height (*EH*) and plant height (*PH*); the latter is missing in the Ner12R trial.
- two flowering time traits: anthesis (*A*) and silking (*Sk*), which are respectively male and female flowering.
- two yield related traits: grain weight (*GW*) and yield (*ϒ*).

Table S7 provides an overview of trait acronyms. Each trial was laid out as an alpha-lattice design, with either two or three replicates. Spatial trends and (in)complete block effects were estimated using the mixed model of Rodríguez-Álvarez *et al.* (2018) (R-package SpATS), and subtracted from the original data; PCgen was then applied to the detrended data, assuming a completely randomized design. Residuals from SpATS appeared approximately Gaussian, and no further transformation was applied.

All traits have direct genetic effects, and traits mostly cluster according to their biological category (height, flowering and yield related), especially in the WW scenarios (Figure 4; panels A and B). In the Ner13W and Ner12R trials (B and C) there are edges between yield and respectively tassel- and plant-height, but these (conditional) dependencies are weak and disappear in the reconstruction with *α* = 0.001 (Figure S2). Much stronger are the edges between yield and one or both of the flowering traits, in the water-deficit trials (C and D); the corresponding conditional independence tests gave highly significant p-values for all of the considered conditioning sets (Table S9). By contrast, in the trial without heat or drought stress (Kar12W), the *ϒ − Sk* and *ϒ − A* edges were already removed in the test conditioning only on the genetic effects; Figure S3 provides an illustration. The relation between yield and delay in silking in maize is well known (see e.g., Borrás *et al.* (2007) and Araus *et al.* (2012)). In the most stressed environment (Bol12R), there is an additional edge between plant height and silking. This may relate to the fact that the timing of anthesis determines the number of phytomeres (number of internodes and leaves) that a plant will generate, which in turns affects plant height (McMaster *et al.* 2005). The strong correlation between anthesis (*A*) and silking (*Sk*) may explain the presence of the edge *PH − Sk* (rather than *PH − A*).

Finally, apart from the Bol12R trial there is never an edge between *ϒ* and *GW*, which seems due to the choice of the genetic material (giving little variation in grain weight) and the design of the trials (targeting stress around flowering time, rather than the grain filling period). See Millet *et al.* (2016) for further details. For all trials, the structure of the graphs is such that none of the between-trait edges can be oriented (technically, this is due to a lack of v-structures). However, for some of these edges, physiological knowledge clearly suggests a certain orientation, in particular for *Sk − ϒ* and *GW − ϒ*.

**Figure 4.**
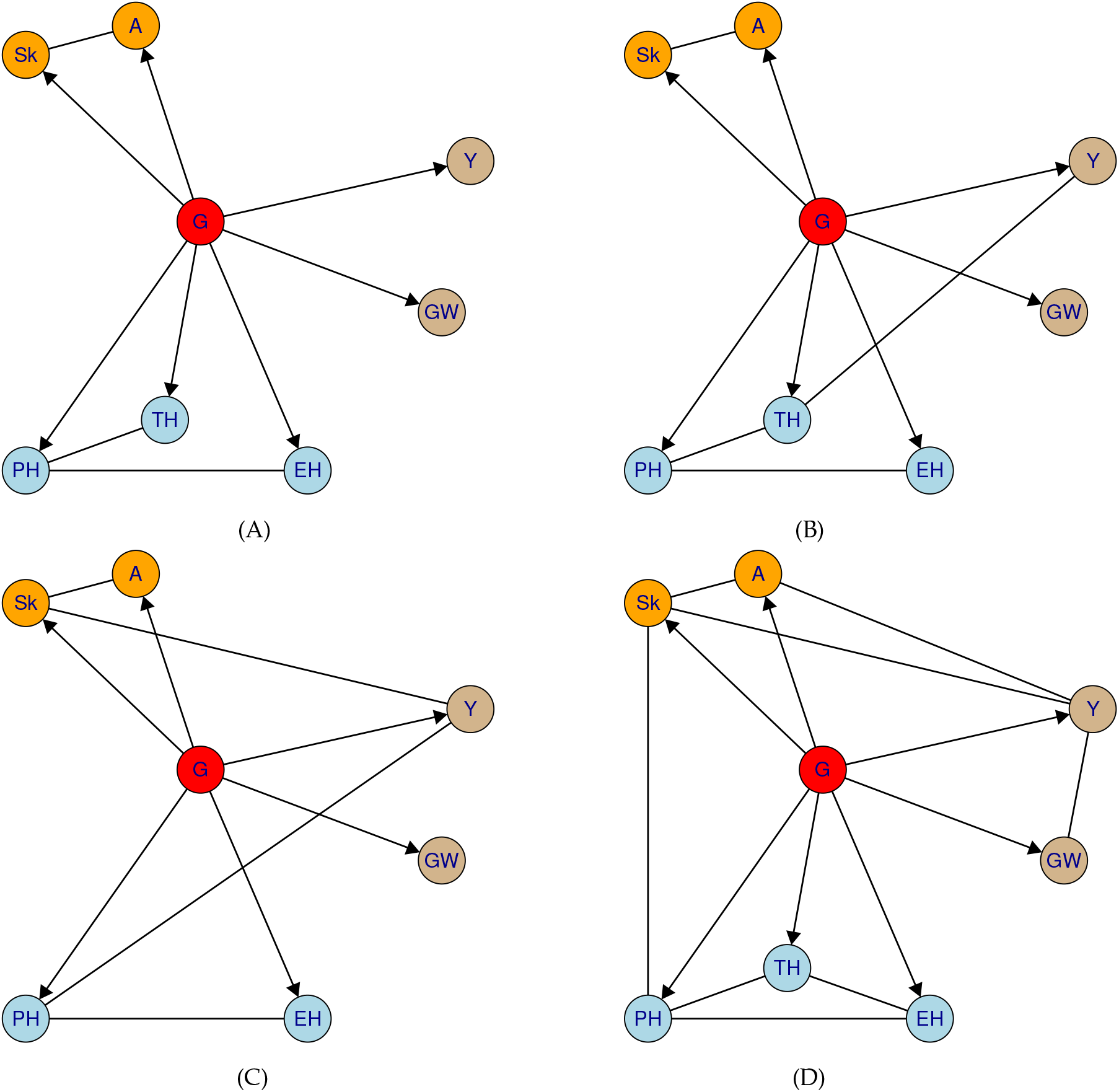
Estimated networks with *α* = 0.01, for four of the DROPS field trials. Trait categories are flowering (orange), height (blue) and yield (brown). Each trial represents a different environmental scenario, being well-watered (WW) or water-deficit (WD), and different temperatures (see text). (A) Kar12W, (WW, Cool), (B) Ner13W (WW, Hot), (C) Ner12R (WD, Hot (daY_S_)), (D) Bol12R (WD, Hot). Trait acronyms are given in Table S7.

The trials also illustrate the difference between the total genetic covariance (*V_G_*) and the covariance among direct genetic effects, as defined by Σ_*G*_. For most pairs of traits, the total genetic correlation^8^ was between *ρ_g_* = 0.3 − 0.9 (Table S10). The (total) genetic correlation between yield and silking was strongly negative in both WD trials (−0.44 and −0.61), and in the Bol12R trial also for yield and anthesis (−0.43). In all trials, genetic correlation with grain weight (*GW*) was negative for most traits, but not always significant. In the Kar12W trial for example, we found *ρ_g_* =−0.010 for *GW* and *PH*, and *ρ_g_* =−0.435 for *GW* and *Sk* (silking). In both cases, the two traits are d-separated in the graph (conditioning on {*G*}), but only for *Sk* the genetic co-variance is significant (*p* = 1.31 · 10^−9^). While this may provide information about Σ_*G*_, we recall that most often, the latter is not entirely identifiable.

As we have seen in the examples following equation (5), the existence of an edge between two traits in the graph does not necessarily imply a strong genetic correlation. In other words, having a shared genetic basis is not the same thing as the presence of a causal effect, found after conditioning on the genetic effects and other traits. In the Ner12R trial, for example, there is no edge between yield and grain weight, but a significant genetic correlation, while in the Bol12R trial it is the other way round.

#### Rice

Next we analyze 25 traits measured on 274 *indica* genotypes of rice (*Oryza sativa*) under water-deficit, reported by Kadam *et al.* (2017). Three replicates of each genotype were phenotyped in a randomized complete block design, and block was included as a covariate in all conditional independence tests. Tests were restricted to conditioning sets of at most four traits. A first run of PCgen produced several inconsistencies in the genetic effects, i.e., traits with significantly positive heritability but without a partially directed path coming from the node *G*. We therefore applied the correction described in File S3, adding edges *G* → *ϒ_j_* for all traits with this inconsistency, and then re-ran PCgen. The final reconstruction is given in Figure 5, where traits are grouped into three shoot morphological traits (blue), one physiological trait (rose), 13 root morphological traits (green), five root anatomical traits (gray) and three dry matter traits (orange).

**Figure 5.**
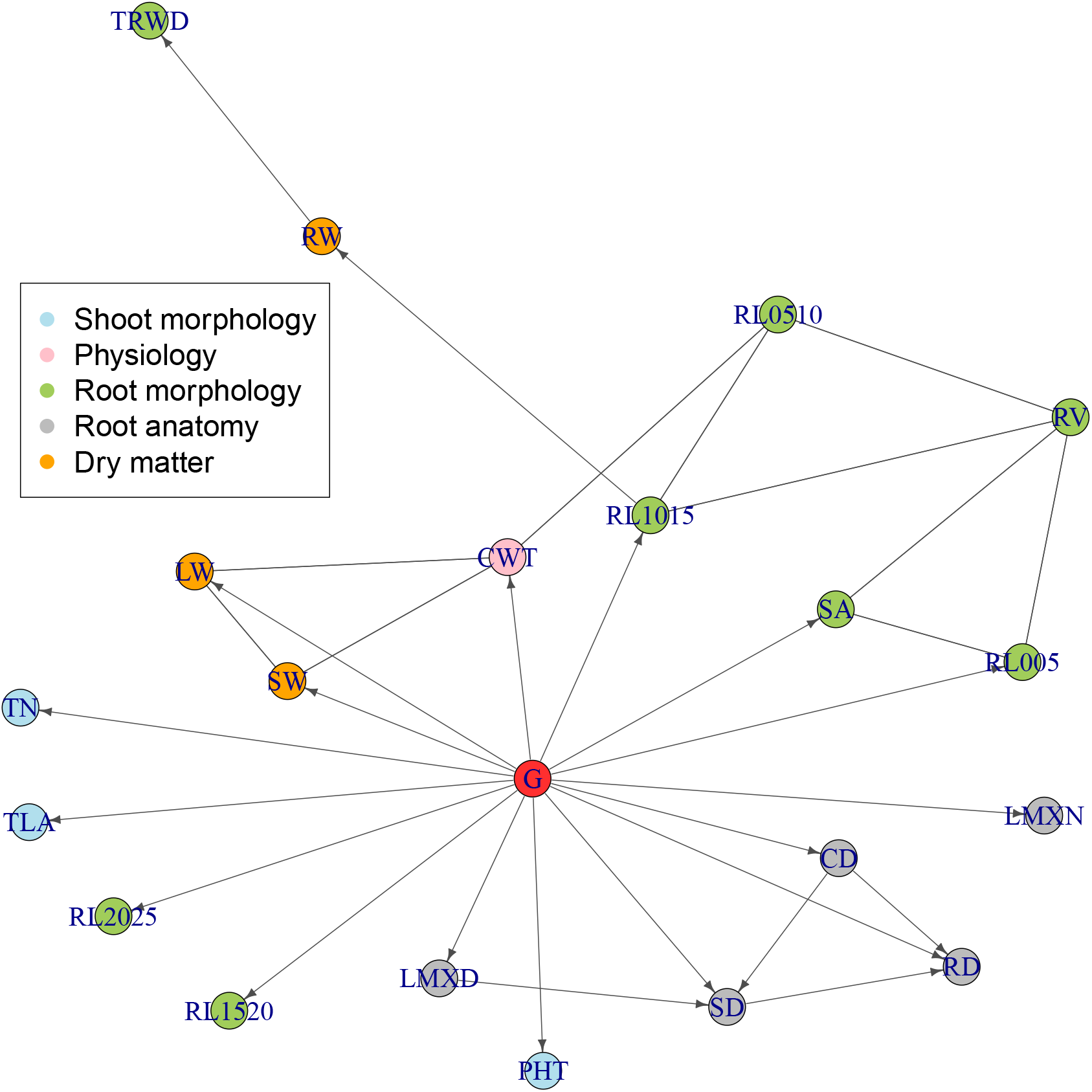
PCgen-reconstruction for the rice-data from Kadam *et al.* (2017), with *α* = 0.01. Five traits (MRL, ART, RL2530, RL3035 and RL35) are not shown, as they were completely isolated in the graph, without any connections to other traits or *G*.

After correcting the inconsistencies, there were nine traits without a direct genetic effect. Five of these (MRL, ART, RL2530, RL3035 and RL35) were completely isolated in the graph, without edges connecting to any other trait. All of these traits are related to either root length, or to the length of thicker roots, which contribute to drought adaptation under field conditions (Uga *et al.* 2013). However, as the experiment was done in pots, roots were constrained in their exploration range and therefore genotypic differences in root length would not translate into differential access to water and biomass (Poorter *et al.* 2012). Four other traits (TRWD, RW, RL0510 and RV) had at least one adjacent trait in the graph, but no direct genetic effect. At a lower significance level (*α* = 0.001, Figure S4), direct genetic effects disappeared also for cumulative water transpiration (CWT), and for three root anatomical traits (RD, CD and SD). For RV (root volume), a direct genetic effect was only present with *α* = 0.001, which was an artefact of the way the initial consistencies were resolved.

Traits related to root surface area (SA), root volume (RV) and roots with small diameter class (RL005, RL1015) had direct genetic effects and were connected amongst each other. As expected, traits related to root volume and area influenced root weight and total root weight density (RW, TRWD). In the recontruction with *α* = 0.001, cumulative water transpired (CWT) was affected by stem and leaf weight (SW, LW) and by RL0510, in agreement with physiological knowledge that water transpiration is influenced by water demand (related to the above-ground biomass) and water supply (related to the roots’ water uptake capacity). The corresponding edges were also present in the recontruction with *α* = 0.01, where however they could not be oriented because of the denser network (in particular, the presence of *G → CWT*). Root anatomical traits (LMXD, SD, CD and RD) appeared as a separate module, not related to the plant water dynamics, suggesting that root anatomy had a smaller impact on water uptake, compared to root biomass.

### Statistical properties of PCgen

We now investigate a number of statistical issues: the assumptions required for asymptotic consistency of PCgen, the assumptions required for faithfulness, and properties of the conditional independence tests. Readers primarily interested in the application of PCgen could skip this section and continue with the Discussion. Proofs of Theorems 1-6 are given in Appendix A.

#### Consistency

Asymptotic consistency holds if, for increasing sample size, the probability of finding the correct network converges to 1. Correct in this context means that we recover the class of partially directed graphs (CPDAG) that contains the underlying DAG. Consistency of the PC-algorithm was shown by Spirtes *et al.* (2001) (for low dimensions) and Kalisch and Bühlmann (2007) (for high dimensions). These authors distinguished between consistency of the oracle version of PC, where conditional independence information is available without error, and the sample version, where conditional independence is obtained from statistical tests. For PCgen we will focus on the oracle version and consistency of the skeleton, leaving the sample version and the correctness of the orientations for future work.

As for the standard PC-algorithm, consistency of PCgen requires the equivalence between conditional independence and d-separation in the graph. Part of this is the Markov property, which states that d-separation of two nodes in the graph given a set of other nodes implies conditional independence of the corresponding random variables. The converse (conditional independence implying d-separation) is known as faithfulness. The following result provides the Markov property for SEM with genetic effects. The proof (Appendix A.9) is a straightforward adaptation of Pearl’s proof for general SEMs (Pearl 2009).

##### Theorem 1

*Suppose we have a GSEM as defined by* *equation* (4), *with a graph 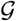 as defined in the Materials and Methods section, and satisfying assumptions 1-4. Then the global Markov condition holds for* 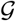 *and the joint distribution of* **G**, **ϒ_1_**, … , **ϒ_p_**. *In particular, d-separation of ϒ_j_ and G given ϒ_S_ implies* **ϒ_j_** ╨ **G**|**{ϒ_S_**}, *and d-separation of ϒ*_*j*_ *and ϒ*_*k*_ *given* {ϒ_S_, *G*} *implies* **ϒ_j_** ╨ **ϒ_k_** | {**ϒ_S_**, **G**}, *for all traits* **ϒ_j_** *and* **ϒ_k_** *and subsets* **ϒ_S_**.

If we now assume faithfulness, the preceding result directly gives the equivalence between conditional independence and d-separation. This in turn implies that PCgen will recover the correct skeleton:

##### Theorem 2

*Let* 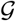 *denote d-separation in the graph* 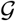. *Suppose we have a GSEM as in Theorem 1, and we make the additional assumptions of faithfulness*:

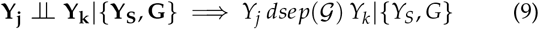

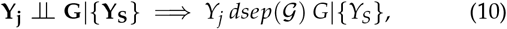

*for all traits* **ϒ_j_** *and* **ϒ_k_** *and subsets* **ϒ_S_**. *Then the oracle version of PCgen gives the correct skeleton*.

#### Faithfulness

For our first faithfulness condition (9), it suffices to have faithfulness for the graph without genetic effects. A necessary (but not sufficient) condition for this is that contributions from different paths do not cancel out (Appendix A.5):

##### Theorem 3

*Let P*_**ϒ**|**U**_ *denote the joint distribution of* **ϒ_1_**, … , **ϒ_p_** *conditional on* **U** = **GΓ**, *the matrix of total genetic effects. Then* ϒ_j_ ╨ ϒ_k_|{ϒ_S_,G} *is equivalent with* ϒ_j_ ╨ _P__Y|U_ ϒ_k_, and 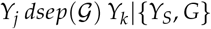 *is equivalent with* 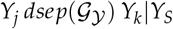. *Therefore* (9) *holds if*

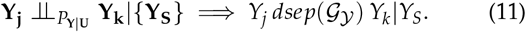

Consequently, we can rephrase (9) in terms of a faithfulness assumption for the analogous SEM without genetic effects.

The second faithfulness statement (equation (10)) involves d-separation of *ϒ*_*j*_ and *G*, and requries that the genetic effects are not collinear. If for example we have **ϒ_3_** = **ϒ_1_** + **ϒ_2_** + **E_3_**, with **ϒ_1_** = **G_1_** + **E_1_**, **ϒ_2_** = **G_2_** + **E_2_**, and **G_2_** = −**G_1_** = **G**, it follows that **ϒ_3_** = **E_1_** + **E_2_** + **E_3_**. Consequently, because **G_3_** = (0, … , 0)^*t*^, we find that **ϒ_3_** and **G** = [**G_1_ G_2_ G_3_**] are marginally independent, but in the graph 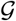, the nodes *ϒ*_*j*_ and *G* are not d-separated by the empty set, as there are directed paths *G* → *Y*_2_ → *Y*_3_ and *G* → *Y*_1_ → *Y*_3_. Conversely, if **G_1_** and **G_2_** are not perfectly correlated, this violation of faithfulness cannot occur. The following theorem shows that marginal independence always implies d-separation. We conjecture (but could not prove) that (10) also holds for non-empty conditioning sets.

##### Theorem 4

*Suppose we have a GSEM sastifying Assumptions 1-5, and faithfulness for the graph without genetic effects, given by* (11). *Then* (10) *holds for S* = *Ø, i.e., marginal independence implies d-separation of ϒ_j_ and G*.

Hence, faithfulness involving **ϒ_j_** and **G** requires (at least) absence of collinearities between genetic effects, as well as faithfulness for the corresponding SEM without genetic effects.

#### Properties of the tests

Theorem 2 provides consistency of the oracle version of PCgen, where conditional independence information is available without error. Proving consistency of the sample version is challenging for two reasons. First, the assumptions made for our conditional independence tests may not always hold, introducing approximation errors. Second, even without these errors, the probabilities of type-I and type-II errors still need to converge to zero with increasing sample size. This is well known for the PC-algorithm with independent Gaussian data (Kalisch and Bühlmann 2007), but more difficult to establish in the presence of genetic effects. Here we address the first issue, leaving the second for future work.

Our tests ^9^ for conditional independence statements (A) and (B) (i.e., **ϒ_j_ ╨ G|{Y_S_**, **G**} and **ϒ_j_** ╨ **ϒ_k_**|{**ϒ_S_**, **G**) rely on the conditional distributions of respectively **ϒ_j_** and *vec*([**ϒ_j_**, **ϒ_k_**]), given the observed **ϒ_S_**:

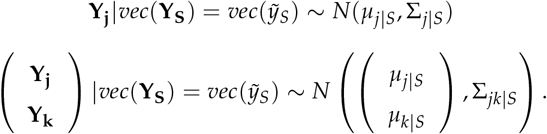

The normality of these distributions directly follows from the assumed normality of the genetic and residual effects. We made the following assumptions about the form of their covariance and mean:

- The covariance matrix Σ_*j|S*_ is that of a single trait mixed model with the same relatedness matrix *K* assumed in the GSEM, i.e.,

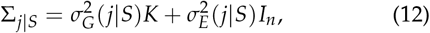

for some variance components 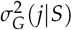 and 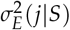.
- The covariance matrix Σ_*jk|S*_ is that of a bivariate MTM, again with the same *K* assumed in the GSEM:

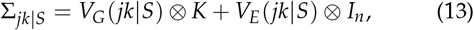

for some 2 × 2 matrices *V*_*G*_ (*jk|S*) and *V*_*E*_(*jk|S*).
- The conditional means *μ*_*j*|*S*_ and *μ*_*k*|*S*_ are linear regressions over the conditioning traits:

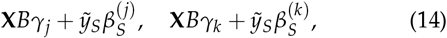

where **X***Bγ*_*j*_ is the marginal mean of **ϒ_*j*_** (see (8)), and 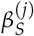 and 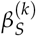 are |*S*| × 1 vectors of regression coefficients.

In the following theorems we show that when *K* = *ZZ*^*t*^, assumptions (12) and (13) always hold, i.e., they directly follow from our GSEM model.

##### Theorem 5

*When K* = *ZZ*^*t*^, *the distribution of vec*([**ϒ_j_**, **ϒ_k_**]) *vec*(**ϒ_S_**) *has covariance of the form given by* (13), *i.e., that of a bivariate MTM. Moreover, under faithfulness condition* (9), *the residual covariance in the MTM is zero if and only if* **ϒ_j_** ╨ **ϒ_k_**|{**ϒ_S_**, **G**}.

##### Theorem 6

*Suppose we have a GSEM as described in Theorem 1, with K = ZZ^t^*. *Then the covariance of* **ϒ_j_**|**ϒ_S_** *is of the form 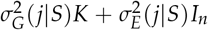, for any conditioning set *S*. *Moreover, assuming the faithfulness condition* (10) *and* Σ_*G*_[*D*, *D*] *of full rank (Assumption 5)*, 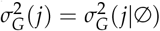 is zero if and only if* **ϒ_j_** ╨ **G**.

Apart from the covariance structure, these theorems address the correctness of our tests. In particular, Theorem 5 shows that the residual covariance in the distribution of {**ϒ_j_**, **ϒ_k_**}| **ϒ_S_** is indeed the right quantity to test statement (B). Similarly, the genetic variance in the conditional distribution of **ϒ_j_**|**ϒ_S_** is the relevant thing for testing (A). This appears to be true for any conditioning set *S*, although we could prove it only for the empty conditioning set, because faithfulness is required (which we also established only for *S* = Ø see Theorem 4).

The situation is different for assumption (14), regarding the conditional means: even when *K* = *ZZ*^*t*^, it holds for certain conditioning sets and not for others. We illustrate this with the following example. Suppose that **ϒ_1_** = **G_1_** + **E_1_** and **ϒ_2_** = *λ***ϒ_1_ + E_2_**, with independent vectors 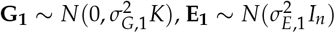 and 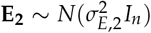. Then the graph 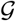 given by *G* → *ϒ*_1_ → *ϒ*_2_. There is no edge *G* → *ϒ*_2_, although this is not essential for the example. The distributions are given by

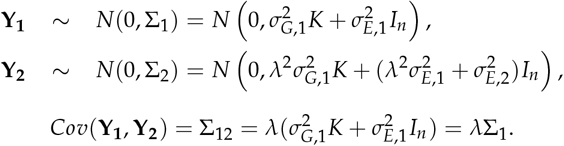

The conditional mean of **ϒ_2_** given **ϒ_1_** = *y*_1_ is 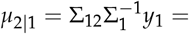 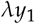. As expected given the graph, the conditional mean is a simple linear regression on *ϒ*_1_. However, the conditional mean of **ϒ_1_** given **ϒ_2_** = *y*_2_ equals

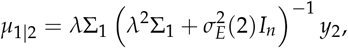

which is a linear transformation, but not a multiple of *y*_2_. In summary, our models for **ϒ_j_|ϒ_S_** and {**ϒ_j_**, **ϒ_k_}ϒ_S_** are sometimes misspecified in terms of the mean, although still correct in terms of covariance, provided *K* = *ZZ*^*t*^ (Theorems 5 and 6). Despite the approximation error occurring sometimes for the conditional means, our tests still seem to perform reasonable, as shown in the simulations above. Assumption (14) is more problematic if relations between traits are nonlinear. Suppose for example that for each individual *i*, **ϒ_2_**[*i*] := (**ϒ_1_**[*i*])^2^ and 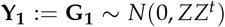, where for the sake of the argument we assume absence of residual errors. Then the factor genotype will generally be significant in the ANCOVA with *ϒ*_1_ as covariate. For example, there could be two replicates of three genotypes, with genetic effects (−1, −1, 0, 0, 1, 1). Then clearly there is some unexplained genetic variance when regressing **ϒ_2_** = (1, 1, 0, 0, 1, 1)^*t*^ on **ϒ_1_**.

Finally, we briefly discuss how the approximation could be improved. In general, the conditional mean is a function of the genetic and residual covariances between **ϒ_j_** and **ϒ_S_**. In Appendix A.7 (equation (23)) we derive that 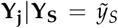 has mean 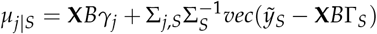. Defining *η*_*j*|*S*_ = 0 for *S* = Ø, we can write *μ*_*j*|*S*_ = **X***B*_*γj*_ + *η*_*j|S*_. Consequently, our approximation of the conditional mean models *η*_*j|S*_ as a linear regression on 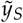. This approximation could probably be improved if we have good estimates of 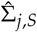 and 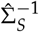, and set 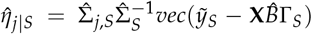. Such estimates however require fitting an MTM for |*S*| + 1 traits, which for large *S* is statistically and computationally challenging, unless pairwise or other approximations are applied (Furlotte and Eskin 2015; Joo *et al.* 2016). Moreover, it seems unclear how 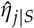 should be incorporated in the tests.

## Discussion

Causal inference for data with random genetic effects is challenging because of the covariance between these effects, and because the usual assumption of independent observations is violated. To address these problems we have proposed a model where random genetic effects are part of the causal graph, rather than a nuisance factor that first needs to be eliminated. The resulting distributions and graphs were shown to satisfy the Markov property. This led us to develop the PCgen algorithm, which tests conditional independence between traits in the presence of genetic effects, and also conditional independence between traits and genetic effects. We showed that the presence of a direct genetic effect can in principle be tested, just like the direct (fixed) effect of a QTL can be tested. This is of course relative to the observed traits, i.e., for any effect *G* → *Y_j_* there may always be an unmeasured trait *Z* such that *G* → *Z* → *Y*_*j*_.

In the linear simulations as well as in the rice data, our tests could indeed identify the absence of many direct genetic effects. By contrast, in the APSIM simulations and maize data all traits had such effects. In the latter case, this could be for biological reasons, i.e., the genetic variance of each trait might really be ‘unique’ to some degree. However, the APSIM results showed that nonlinearities could increase the false positive rate in the edges *G* → *Y*_*j*_, which may be avoided in future versions with better conditional independence tests. Such tests might also allow for non-Gaussian data.

In our simulations PCgen outperformed existing approaches, regarding the reconstruction of between trait relations. Part of this improvement is due to phenotypic information on replicates, reducing the number of errors in the tests. Another part is due to the improved orientation, which is a consequence of the additional edges *G* → *Y*_*j*_. Compared to previous algorithms, PCgen also appeared to be computationally more efficient: depending on the choice of independence tests and the sparseness of the network, it can analyze around 10-50 traits on a single core, and many more if we limit the maximum size of the conditioning sets, or would parallelize the conditional independence tests.

As for the original PC-algorithm, PCgen is most efficient for sparse graphs, i.e., when each trait is connected to only a few other traits, and when there are few direct genetic effects. But even if this is not the case PCgen still has an advantage over existing approaches: by incorporating the genetic effects in the PC-algorithm, we do not need to fit an MTM for all traits simultaneously, but only bivariate models. Our approach also makes genetic network reconstruction feasible with just two traits, and in absence of QTLs, or even no genotypic data at all.

As any causal inference method, PCgen only suggests causal models that are in some sense compatible with the data, and cannot validate the existence of a functional relationship, which is only possible through additional experiments. Because of the required assumptions, the identifiability issues and the uncertainty in the estimated networks, it may be better to speak of algorithms for causal *exploration* than causal *discovery*. At the same time, analysis of one trait conditional on other traits (e.g., yield given plant height) is a common and natural thing to do (Stephens 2013). From that perspective, PCgen could be seen simply as a tool that performs such analyses systematically, compares them and visualizes the results. PCgen results for different significance levels could then be reported alongside other ‘descriptive’ statistics like heritability estimates and genetic correlations, suggesting functional hypotheses interesting for future research.

### Dealing with derived traits

We analyzed the maize and rice datasets using the traits as they were measured, without adding ‘derived’ traits defined by ratios, sums or differences of the original traits. Because such derived traits are not measured themselves there is no error associated with them, apart from ‘copies’ of errors in the original trait. For example, if there is much variation in leaf weight but almost none in the total weight of a plant, the derived trait ‘leaf weight ratio’ will be essentially a copy of the original leaf weight trait. This can violate our assumption of faithfulness and lead to errors in the reconstruction; see Figure 7 in Appendix A for an example. Sometimes, derived traits are biologically highly relevant. It may then be desirable to include them in the analysis, and omit some of the original traits. Alternatively, derived traits may be added after running PCgen. For example, we may extend the reconstructed graph with the node *ϒ*_3_ := *ϒ*_1_ + *ϒ*_2_ and edges *ϒ*_1_ → *ϒ*_3_ and *ϒ*_2_ → *ϒ*_3_, provided that biologically this makes sense.

### Data from different experiments

We assumed traits to be measured on the same individuals in the same experiment, with residuals errors arising from biological variation (Assumption 1). In certain applications this assumption can indeed be restrictive, but seems to be inevitable. Suppose traits were measured in different experiments, or residual errors would only come from measurement errors. Then there would be no propagation of residual errors, and the reconstruction would rely completely on how the *genetic* effects are propagated through the network. The GSEM model (4) would need to be replaced by **ϒ** = **X***B* + **U** + **E** and 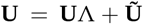, where **Ũ** in a sense models direct genetic effects, reminiscent of the genomic networks of Töpner *et al.* (2017). However, if data are actually generated by equation (4), these can only partially reconstruct the direct genetic effects, even without any type-I or type-II error in the tests (see File S7.2).

Biologically, the genomic networks have a different interpretation: for example, we would assume that *the genetic component* in high blood-pressure causes some cardiovascular disease, rather than high blood-pressure itself. The alternative model implies that the observed traits have diagonal residual covariance, instead of the matrix Γ^*t*^Σ_*E*_ Γ obtained under assumption 1 (see equation (5)). However, the latter matrix turned out to be essential for network reconstruction (see e.g., Theorems 5-6 above). Without assumption 1 we would therefore need to rely completely on the genetic effects. This in turn would require Σ_*G*_ to be diagonal, which appears rather unrealistic. A relevant alternative approach here is that of invariance causal prediction (Peters *et al.* 2016), which infers causal effects that are consistent across several experiments, but still requires all traits to be measured in each experiment (as well as low genotype-by-environment interaction).

### Replicates versus means

In principle PCgen allows for any type of genetic relatedness. We have however focused on the case of independent genetic effects, for the following reasons:

- Performance under model misspecification: different types of genetic effects could in theory be represented by introducing multiple genetic nodes, with conditional independence tests that can distinguish between these effects. But this seems difficult in practice due to the computational requirements and lack of statistical power (Uhler *et al.* 2013; Blair *et al.* 2012; Kruijer 2016). For this reason it seems, previous work on network reconstruction used genotypic means and an additive GRM. For the analysis of a single trait however, Kruijer (2016) showed that broad-sense heritability estimates (obtained with *K* = *ZZ*^*t*^) capture any type of genetic effect, while a model assuming only additive effects can produce strongly biased heritability estimates, if the actual genetic effects are for example partly epistatic. It seems plausible that this robustness extends to the multi-variate models considered here, for example when direct genetic effects are driven by different sets of QTLs, leading to trait-specific relatedness matrices.
- Higher power: estimates of (total) genetic variance based on replicates are typically more accurate than marker-based estimates based on genotypic means (Kruijer *et al.* 2015; Visscher and Goddard 2015), and the use of replicates is therefore also likely to improve hypothesis testing. For the reconstruction of trait-to-trait relations with PCres, our simulations indeed suggest that replicates give more power. Mixed models with *both* replicates and a GRM might further increase power if the true architecture is really additive (Kruijer *et al.* (2015)), but also these models lead to biased inference if the actual architecture is different (Kruijer (2016)).
- When *K* = *ZZ*^*t*^, the conditional independence statement considered in the RC-test is completely equivalent with **ϒ_j_ ╨ ϒ_k_**|{**ϒ_S_**, **G**} (Theorem 5), while for other *K* it is not, and an alternative test might be required.

Apart from these statistical issues, there is also a computational advantage: the test for **ϒ_j_** ╨ **G**|**ϒ_S_** can be based on standard ANCOVA, which is many times faster than the LRT for a mixed model. Also the tests for **ϒ_j_ ╨ Y_k_**|{**ϒ_S_**, **G**} are faster when *K* = *ZZ*^*t*^.

Finally, we have not investigated the performance of PCgen for unbalanced designs, but it seems likely that small unbalancedness has only a minor effect. A more fundamental challenge seems to be the presence of incomplete blocks or spatial trends (Rodríguez-Álvarez *et al.* 2018; Flaxman *et al.* 2015).

### Assessing uncertainty

If one mistakenly rejects the null-hypothesis of conditional independence (type-I error), PCgen leaves the corresponding edge, although it may still be removed at a later stage, with a different conditioning set. If the null-hypothesis is mistakenly not rejected (type-II error), a true edge is removed, and will not be recovered. Moreover, it may affect the remaining tests, since d-separation of *ϒ*_*j*_ and *ϒ*_*k*_ is only tested given conditioning sets contained in *adj*(*ϒ*_*j*_) or *adj*(*ϒ*_*k*_), where the adjacency sets are defined relative to the current skeleton. This is correct in the oracle version, but in the sample version of PC(gen), *adj*(*ϒ*_*j*_) or *adj*(*ϒ*_*k*_) may become smaller than the corresponding adjacency sets in the true graph, and the algorithm may therefore not perform an essential independence test. See Colombo and Maathuis (2014) for examples.

Consequently, assessing uncertainty for constraint-based algorithms is difficult, and cannot be achieved by just applying some multiple testing correction to the p-values. To obtain bounds on the expected number of false edges in the skeleton, several authors have used stability selection (Meinshausen and Bühlmann 2010; Stekhoven *et al.* 2012) or other sample-splitting techniques (Töpner *et al.* 2017), but these are often too stringent and require an additional exchangeability assumption (Bühlmann *et al.* 2013; Meinshausen *et al.* 2016). Moreover, these approaches do not provide a level of confidence for *specific* edges. For example, an edge present in 60% of all subsamples may appear to be present in the true graph with a probability of 0.6, but there is no justification for such a statement.

Alternatively, uncertainty may be assessed using Bayesian methods, which are however computationally very demanding and outside the scope of this work. Moreover, despite the recent progress in Bayesian asymptotics (Ghosal and van der Vaart 2017), there do not seem to be results regarding the correct coverage of posteriors in these models.

### Genomic prediction

PCgen can select traits with direct genetic effects, which are the most relevant in genomic selection. More generally, the usefulness of structural models for genomic selection depends on whether there is an interest in some kind of intervention (Valente *et al.* 2013, 2015). Informally speaking, an intervention is an external manipulation that forces some of the traits to have a particular distribution. For example, with a so-called hard intervention on the *j*th trait, **ϒ_j_** is forced to a constant level *c*, e.g., *c* = 0, when **ϒ_j_** is the expression of a gene that is knocked out. The manipulation or truncated factorization theorem (Pearl 2009; Spirtes *et al.* 2001) can then predict the joint distribution of the system *after* the intervention:

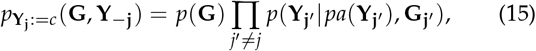

where *pa*(**ϒ_j′_**) is the set of parents of *ϒ_j′_*. This is generally different from the distribution

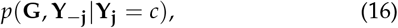

obtained from conditioning on **ϒ_j_** = *c*, *prior* to the intervention (see e.g., Peters *et al.* (2017)). In other words, *conditioning* is not the same as *doing* (intervening). File S7.3 and File S7.4 provide examples, showing that the use of (16) can lead to improved genomic prediction. When however the intervention is on a root node, (15) and (16) are the same (see again Peters *et al.* (2017), p. 110).

The example in File S7.4 is special in the sense that PCgen could correctly infer the complete graph, for most of the simulated datasets. A technical obstacle for a more general use of our networks in genomic prediction is the identifiability issue mentioned in the introduction. PCgen (and the PC-algorithm in general) typically outputs a *partially* directed graph, several DAGs being compatible with this graph. This is particularly problematic for edges between the traits in *D* (the traits with direct genetic efffects). For the traits with only indirect genetic effects, it is in principle possible to estimate how much of the genetic variance originates from a partiular trait in *D*, the result being independent of the chosen DAG. This would first require estimates of the (total) genetic covariance among traits in *D*, obtained either by fitting an MTM, or by an approximation (as in Furlotte and Eskin (2015)).

In absence of interventions on the traits, we can think of genomic prediction in terms of an intervention on the node *G*. Because the latter is a root node by definition, standard genomic prediction methods can in principle have optimal performance (Valente *et al.* 2013). More specifically, genomic prediction usually involves a regression of a target trait on a number of markers, having either fixed or random effects. In either case, it is only the total effect of genotype on the target trait that matters, not through which other traits this effect passes.

Optimal prediction accuracy however requires that the regression model contains the true distribution (or a good approximation), and a sufficiently accurate estimate of this distribution. We therefore believe that structural models may sometimes be an appealing alternative, especially if the underlying model is highly nonlinear, or when prior physiological knowledge can be incorporated. The extent to which this can really improve accuracy remains to be investigated.

### Open questions and extensions

Although we have shown the Markov property for our model and studied consistency of PCgen, there are a number of open questions left for future work. First, it may be possible to construct better tests, especially for nonlinear structural models and non-Gaussian error distributions. The recent work of Pfister *et al.* (2018) seems particularly relevant here. A second issue is the consistency of the orientations: while we have shown PCgen’s consistency in reconstructing the skeleton, we did not address this for the final CPDAG. This is well known for the PC-algorithm without genetic effects (Spirtes *et al.* 2001; Kalisch and Bühlmann 2007), but more difficult to establish here, as the class of CPDAGs needs to be restricted to those without errors pointing to *G*. More generally, orientation constraints seem to be of interest for trait-to-trait relationships as well, e.g., one may require that, *if there is an edge*, the expression of a gene can only affect a metabolite and not the other way round. To the best of our knowledge, current methodology and theory has only considered the forced absence/presence of an edge, leaving the orientation to the algorithm^10^. A final question for future work is whether Theorems 4 and 6 hold for general conditioning sets.

Apart from these open questions, we believe that the idea of explicitly modeling direct genetic effects can be applied more generally. In particular, we hope that the ideas developed here provide a first step towards the more amibitious goal of modeling multiple traits through time, simultanesouly for many environments. A first generalization would be to replace the PC-algorithm with other constraint-based algorithms, in particular FCI and RFCI (Spirtes *et al.* 2001; Colombo *et al.* 2012). These have the advantage that the causal sufficiency assumption (no latent variables) can be dropped or considerably weakened. The presence or absence of direct genetic effects could also be incorporated in (empirical) Bayesian approaches for genetic network reconstruction, or in invariant causal prediction (Peters *et al.* 2016). It might also be possible to extend the approach of Stephens (2013), and focus only on the detection of traits with direct genetic effects. Another application of GSEM might be as covariance models in multi-trait GWAS, as an alternative to unstructured (Zhou and Stephens 2014) or low-rank (Millet *et al.* 2016) models. Finally, the concept of direct and indirect genetic effects may be useful in deep-learning models for high-dimensional phenotypes, observed on genetically diverse individuals.

## Author contributions

WK developed the PCgen algorithm. PB developed the package pcgen, based on code written by WK and the EM-algorithm contributed by MR. WK wrote the paper, with input from FvE, DBK, EW, BY, PB and MR. DBK simulated data with APSIM, and analyzed the rice data. PB visualized the estimated networks for the rice, maize and APSIM data. WK proved Theorem 1-2 and WK, EW and MM proved Theorems 3-6.

## Acknowledgements

WK was funded by the Learning from Nature project of the Dutch Technology Foundation (STW), which is part of the Netherlands Organisation for Scientific Research (NWO). MX was funded by project MTM2017-82379-R (AEI/FEDER, UE), by the Basque Government through the BERC 2018-2021 program and by the Spanish Ministry of Science, Innovation and Universities (BCAM Severo Ochoa accreditation SEV-2017-0718). EW acknowledges support from the EU COST Action CA15109. We thank Niteen Kadam for providing the rice data, and Xinyou Yin for useful discussions on the interpretation of the resulting networks. Emilie Millet and François Tardieu are acknowledged for providing and interpretating the maize data.

## Appendix A. Faithfulness, conditional distributions and proofs of Theorems 1-6

## A.1. Overview of graph theoretic definitions

Definitions of for example d-separation and CPDAGs can be found in many books and articles on graphical models and causal inference; see for example Lauritzen (1996); Pearl (2009); Spirtes *et al.* (2001); Kalisch and Bühlmann (2007). The following summary was inspired by Shipley (2016) and Maathuis (2014).

- Given different nodes *ϒ_j_* and *ϒ_k_*, a *path* from *ϒ_j_* to *ϒ_k_* is a sequence of edges connecting *ϒ_j_* and *ϒ_k_*. When all edges are directed and pointing towards the same node, we have a *directed path*. A path that is not directed is an *undirected* or *non-directed* path.
- A *cycle* is a path from *ϒ_j_* to *ϒ_k_* with an additional edge between *ϒ_j_* and *ϒ_k_*. A *directed cycle* is a directed path from *ϒ_j_* to *ϒ_k_* together with a directed edge *ϒ_k_* → *ϒ_j_*.
- A *directed acyclic graph* (DAG) is a directed graph without any directed cycle. When a graph underlying a SEM is a DAG, the SEM is said to be *recursive*.
- *pa*(*ϒ_j_*) is the set of nodes *ϒ_k_* for which there is a directed edge *ϒ_k_* → *ϒ_j_*; in this case *ϒ_j_* is a *child* of *ϒ_k_* and *ϒ_k_* is a *parent* of *ϒ_j_*. The nodes *ϒ_j_* and *ϒ_k_* are *adjacent* if there is an edge *ϒ_k_* → *ϒ_j_*, *ϒ_j_* → *ϒ_k_* or *ϒ_k_* → *ϒ_j_*.
- If in a DAG 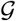 there is a directed path from *ϒ_j_* to *ϒ_k_*, then *ϒ_k_* is a *descendant* of *ϒ_j_*, and *ϒ_j_* is an *ancestor* of *ϒ_k_*.
- In a DAG with nodes *ϒ*_1_, … , *ϒ_p_*, it is always possible to relabel the nodes, such that for each node *ϒ_j_*, *k < j* for all parents *ϒ_k_* in the set *pa*(*ϒ_j_*). Such a relabeling is known as a *topological ordering* of the DAG. Using this ordering, the root nodes (without any parents) have the lowest labels, the sink nodes (without any children) have the highest labels, and the matrix of path coefficients Λ is upper triangular. Formally, a topological ordering is a function *π* : {1, … , *p*} → {1, … , *p*} such that the preceding property holds. A topological ordering always exists, and does not need to be unique (Peters *et al.* 2017).
- If, for a given path, two directed edges point into the same node, the latter is a *collider*. For example, given the DAG *A → C ← B*, *C* is a collider on the (only) path between *A* and *B*. In all other cases (*A ← C → B*, *A → C → B* and *A ← C ← B*), *C* is a *non-collider*. Several different paths can pass through a node, and being a (non-)collider is always relative to the path.
- In a DAG, a *v-structure* or *immorality* is a collection of three nodes (say *A*, *B* and *C*), such that there are directed edges *A → B* and *C → B* but no edge between *A* and *C*. In this case *B* is an *unshielded* collider. If there *is* an edge between *A* and *C*, it is a *shielded* collider. Similarly, in an undirected graph, *A*, *B* and *C* form an *unshielded triple* if there are edges *A − B* and *C − B* but no edge *A − C*.
- The *skeleton* of a (partially) directed graph is the undirected graph obtained after removing all arrowheads.
- Given a directed graph 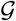, two nodes *A* and *B*, and a (possibly empty) subset of nodes *S* not containing *A* and *B*, a path between *A* and *B* is *blocked* by *S* if at least one of the following two conditions holds: (i) there exists a collider on the path which is not in *S*, and also none of its descendants are in *S*. (ii) there exists a non-collider on the path that is in *S*.
- Nodes *A* and *B* are *d-separated* by a set *S* if *S* blocks all paths from *A* to *B*.
- Given disjoint sets *U*, *V* and *S* (*U* and *V* should be non-empty), *U* and *V* are *d-separated* by *S* if *S* blocks all paths from *ϒ_j_* to *ϒ_k_*, for all nodes *ϒ_j_ ∈ U* and *ϒ_k_ ∈ V*.
- Two DAGs are *equivalent* if they have the same skeleton and the same v-structures.
- An equivalence class of DAGs is a set containing all DAGs that are equivalent to one another. For example, given a skeleton *A − B − C*, there is one equivalence class containing the three DAGs *A → B → C*, *C → B → A* and *A ← B → C*, and one equivalence class with only one DAG (*A → B ← C*). Any DAG in the class can be used to represent the class. But instead of picking an arbitrary DAG, an equivalence class can also be represented by a *completed partially directed acyclic graph* (CPDAG). A *partially directed acyclic graph* (PDAG) is ‘a graph where some edges are directed and some are undirected and one cannot trace a cycle by following the direction of directed edges and any direction for undirected edges’ (Kalisch and Bühlmann 2007). A PDAG is a CPDAG if (a) every directed edge in the PDAG exists in all DAGs in the equivalence class it represents (b) for every undirected edge *A − B* in the PDAG, the equivalence class contains at least one DAG with *A → B* and at least one with *B → A*. Chickering (2002) showed that CPDAGs represent equivalence classes uniquely.

## A.2. Identifiability

In a general GSEM, the parameters in Σ_*G*_, Σ_*E*_ and Λ are not identifiable, which was pointed out by Gianola and Sorensen (2004). However, when we know a topological ordering of the graph (defined above in Appendix A.1), and Assumption 3 (diagonal Σ_*E*_) and faithfulness assumptions (9)-(10) hold, it appears that Σ_*G*_, Σ_*E*_ and Λ are identifiable from the joint distribution of **ϒ_1_**, … , **ϒ_p_** (see equation (7)). Our approach relies on the *L^t^DL* decomposition of *V_E_*, and is probably known, although we could not find it in the literature. We could neither find a proof, but the decomposition seemed valid in all examples we considered. It works as follows:

1. Relabel the nodes (traits) *ϒ*_1_, … , *ϒ_p_* according to a topological ordering. Then both Λ and Γ = (*I* − Λ)^−1^ are upper triangular. Recall that Λ has zeros on the diagonal. Γ always has ones on the diagonal, i.e., it is *unit* upper-triangular.
2. Now we use the fact that every positive definite matrix *A* can be decomposed as *A* = *LDL^t^* = *L^t^DL*, with a diagonal matrix *D* and unit upper-triangular *L* (see e.g. Petersen *et al.* (2008), section 5.7). We apply this result to *V_E_*:

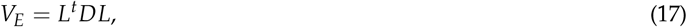

and set Σ_*E*_ equal to *D*, and Γ equal to *L*. Let Λ = *I* − Γ^−1^.
3. Finally, using *V_G_* and *L* we obtain Σ_*G*_:

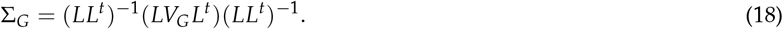

For example, consider the graph *ϒ*_3_ → *ϒ*_2_ → *ϒ*_1_, with path coefficients and unit error variances (for simplicity we ignore the genetic effects in this example). We need to relabel the graph such that *ϒ*_1_ → *ϒ*_2_ → *ϒ*_3_. After relabeling, we have

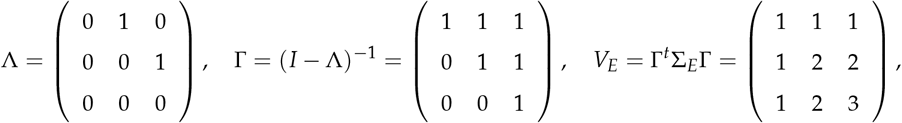

as Σ_*E*_ is the identity matrix. In R the *L^t^ DL* decomposition can be computed as follows:

~~~
b <- matrix(c(1,1,1,1,2,2,1,2,3), ncol = 3)
#b <- b[c(3,2,1), c(3,2,1)]
v <- Cholesky(Matrix(b, sparse = T), LDL=T, perm = F)@x
Gamma <- matrix(0,3,3)
Gamma[lower.tri(Gamma, diag=T)] <- v
D <- diag(diag(Gamma))
diag(Gamma) <- 1
Gamma <- t(Gamma)
Lambda <- diag(3) - (solve(Gamma))
~~~

We emphasize that the topological ordering is crucial. Without the relabeling (e.g., uncomment the second line the above R-code), a different Γ is obtained (also after interchanging the first and third row). Although in this example the topological ordering is unique, there may in general be multiple valid orderings; e.g., *ϒ*_1_, *ϒ*_3_, *ϒ*_2_ and *ϒ*_3_, *ϒ*_1_, *ϒ*_2_ in case the graph is *ϒ*_1_ → *ϒ*_2_ ← *ϒ*_3_. Based on all investigated examples we conjecture (but could not prove) that these orderings lead to the same parameter estimates.

## A.3. The matrix Γ expressed as a function of path coefficients

Let 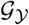 denote the DAG over the nodes *ϒ*_1_, … , *ϒ_p_*, with edges defined by Λ. For each *j* ∈ {1, … , *p*}, let *V_j_* denote the union of the set *ϒ_j_* and the set of root traits (i.e., those without parents in 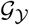) for which there is a directed path towards *ϒ_j_*. For all *j*, *k* ∈ {1, … , *p*}, let Π_*jk*_ denote the set of all directed paths from *ϒ_j_* to *ϒ_k_*. For *k* = *j*, Π_*jj*_ contains only the empty path from *ϒ_j_* to itself. For any directed path *π* from *ϒ_j_* to *ϒ_k_*, let *L*(*π*) denote the product of the corresponding path coefficients as given by Λ; for the empty path we define *L*(*π*) = 1.

Using these definitions, we can decompose the variance of a trait into contributions from different ancestors, as well as its own error variance. To this end, we follow Spirtes *et al.* (2001) and define the *p* × 1 column vector *γ_j_* with elements (*l* = 1, … , *p*)

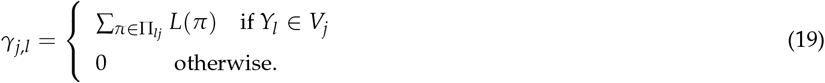

## A.4. The covariance between ϒ_j_ and ϒ_k_ as function of path coefficients

Since **ϒ_j_** = **X***Bγj* + **G***γ_j_* + **E***γ_j_* (equation (8) in the main text), the covariance between the *n* × 1 vectors **ϒ_j_** and **ϒ_k_** can be written in terms of *γj* and *γk*:

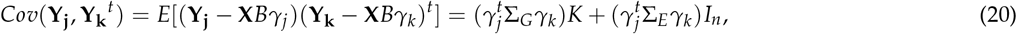

for all *j*, *k* ∈ {1, … , *p*}. Consequently, we can express the genetic and residual covariance between traits in terms of quadratic forms, involving Σ_*G*_, Σ_*E*_ and the path coefficients.

As a special case of (20), it follows that without random genetic effects,

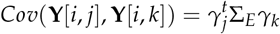

is the covariance between the *j*th and *k*th trait, for each individual *i*. See also Spirtes *et al.* (2001) (Lemma 3.1.6), or Lynch and Walsh (1998) (Appendix 2). Using standard expressions for multivariate Gaussian distributions, this implies that

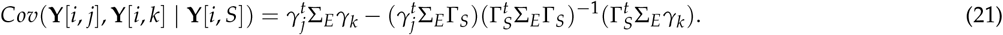

## A.5. The path coefficients condition

It is well known that faithfulness is violated when contributions from different paths cancel out. For example, in the SEM defined by *ϒ*_1_ → *ϒ*_2_, *ϒ*_1_ → *ϒ*_3_ and *ϒ*_2_ → *ϒ*_3_, with respective path coefficients 1, 1 and −1, *ϒ*_1_ and *ϒ*_3_ are marginally independent but not d-separated. Conversely, when faithfulness holds, we know that such cancellations cannot occur, and that the sum in (19) is never zero, i.e., *γ*_*j*,*l*_ = 0 only for *ϒ_l_* ∉ *V_j_*. We will refer to this as the **path coefficients condition**.

**Figure 6.**
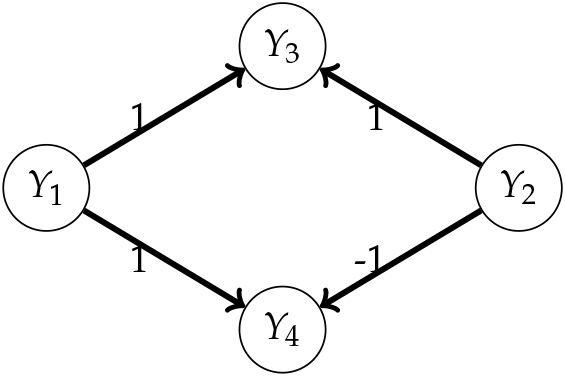
An example of a SEM where faithfulness does not hold, because the contributions to the covariance from the treks *ϒ*_3_ ← *ϒ*_1_ → *ϒ*_4_ and *ϒ*_3_ ← *ϒ*_2_ → *ϒ*_4_ cancel out. If *ϒ*_1_ and *ϒ*_2_ are Gaussian with equal error variances, it follows that for every individual *i*, *Cov*(**ϒ_3_**[*i*], **ϒ_4_**[*i*]) = *Cov*(**ϒ_1_**[*i*] + **ϒ_2_**[*i*] + **E_3_**[*i*], **ϒ_1_**[*i*] − **ϒ_2_**[*i*] + **E_4_**[*i*]) = *Cov*(**ϒ_1_**[*i*] + **ϒ_2_**[*i*], **ϒ_1_**[*i*] − **ϒ_2_**[*i*]) = 0. Consequently, **ϒ_3_** and **ϒ_4_** are marginally independent, but not d-separated by the empty set.

**Figure 7.**
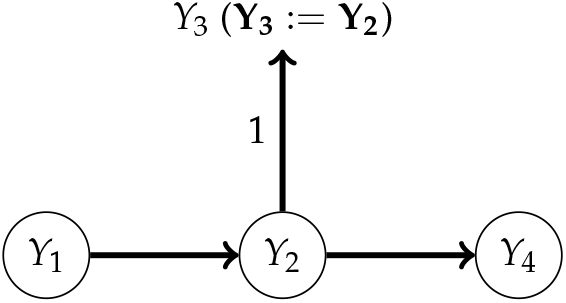
An example of a SEM where faithfulness does not hold, because the variance of the error variables in *E*_2_ is zero. The random vectors **ϒ_1_** and **ϒ_4_** are conditionally independent given **ϒ_3_**, but in the graph, the nodes *ϒ*_1_ and *ϒ*_4_ are not d-separated by *ϒ*_3_.

## A.6. The path coefficients condition and faithfulness

The path coefficients condition is a necessary but not a sufficient condition for faithfulness. First, faithfulness can also be violated when contributions from different paths cancel out when summing over a subset of all directed paths; see Example 2.10 in Peters (2012). Second, it is not only the contributions of directed paths that should not cancel out, but also those of treks. A trek between *ϒ_j_* and *ϒ_k_* is any path between these nodes without a collider (Spirtes *et al.* 2001). Every trek consists of 2 directed paths, starting at the *source* of the trek, and going towards *ϒ_k_* and *ϒ_k_*. One of these can be the empty path; hence each directed path is also a trek. Figure 6 provides an example where contributions from different treks cancel out, leading to non-faithfulness.

Another necessary condition for faithfulness is that all error variances are strictly positive. Figure 7 provides an example of non-faithfulness due to a zero error variance. An extended version of the path coefficients condition (involving sums over subset of treks) together with strictly positive error variances may be sufficient for faithfulness, but we could not find such a result in the literature. However, from (21) it follows that for Gaussian linear SEM, faithfulness is equivalent with

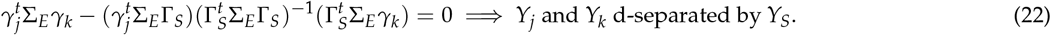

## A.7. Conditional means and covariances

Using the notation [, *S*] to select the columns corresponding to *S*, and [*S*_1_, *S*_2_] to select both rows and columns, it follows from (7) that 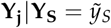 is multivariate normal with mean and covariance

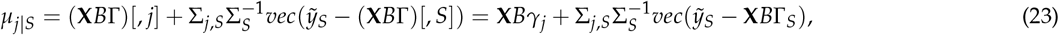

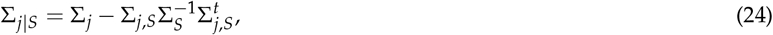

where

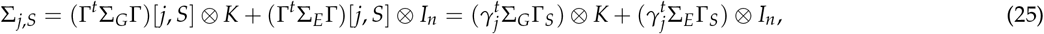

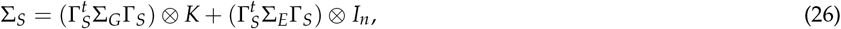

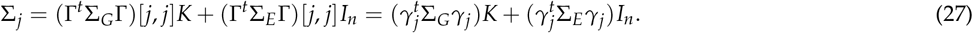

The matrices Σ_*j*_, Σ_*S*_ and Σ_*j*,*S*_ are the variance-covariance matrix of respectively *vec*(**ϒ_j_**) = **ϒ_j_** and *vec*(**ϒ_S_**), and the covariance between **ϒ_j_** and *vec*(**ϒ_S_**). From equation (7) in the main text we also obtain the conditional distribution

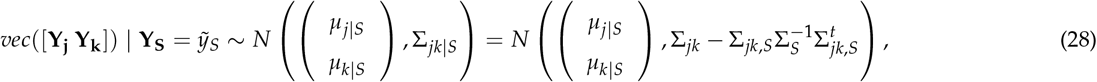

where *μ*_*j*|*S*_ and *μ*_*k*|*S*_ are as in equation (23), and Σ*jk* is the 2*n* × 2*n* block matrix with diagonal blocks Σ_*j*_ and Σ_*k*_ (defined as in (27)), and off-diagonal blocks 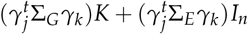. Similarly, given the *p* × 2 matrix Γ_*jk*_ with columns *γ_j_* and *γ_k_*, it follows that

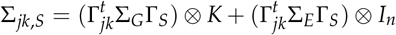

is the 2*n* × |*S*|_*n*_ covariance between *vec*([**ϒ_j_ ϒ_k_**]) and *vec*(**ϒ_S_**).

## A.8. Covariance structure of the conditional distributions

When *K* = *ZZ^t^* is block-diagonal, with *m* blocks of ones of dimension *r* × *r* on the diagonal, then for any positive constants *c* and *d*,

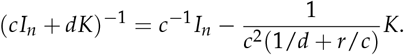

Hence, the inverse of (*cI_n_* + *dK*) is again a linear combination of *I_n_* and *K*. This follows from the Woodbury identity (Petersen *et al.* 2008; Golub and Van Loan 2012)

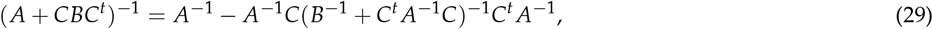

with *A* = *cI_n_*, *B* = *dI_m_* and *C* = *Z*. In addition we have *Z^t^Z* = *rI_m_*, and therefore *K*^2^ = *rK*. Consequently, any product of matrices of the form (*cI_n_* + *dK*) or their inverse is a linear combination of *I_n_* and *K*.

From this it follows that when *K* = *ZZ^t^*, Σ_*j*|*S*_ in (24) is of the form 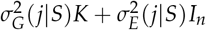. Similarly, it follows that Σ_*jk*|*S*_ (equation () in the main text) is of the form

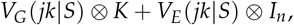

for some 2 × 2 matrices *V_G_* (*jk*|*S*) and *V_E_* (*jk*|*S*).

## A.9. Proofs of Theorems 1 and 2

Pearl (2009) (p. 51) showed that under quite general assumptions, structural equation models satisfy the global Markov property, which means that d-separation in the graph implies conditional independence. It turns out that in our case, the required assumption of independent errors applies to the *p* error variables and not to *G*. The intuition behind this is that *G* is not just an additional error node, but part of the causal graph, and we can always distinguish between residual (co)variance and genetic (co)variance. We now give the proof of Theorem 1, which only requires minor modifications of the proof given by Pearl for the case without the genetic effects.

Let 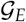 denote the graph, obtained by extending 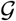 with the error variables, i.e., for traits *j* = 1, … , *p* we add the node *E_j_* and an edge *E_j_* → *ϒ_j_*. We first show that the local Markov property holds for 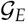, i.e., for any variable *Z* ∈ {*G*, *ϒ*_1_, … , *ϒ_p_*, *E*_1_, … , *E_p_*}, *Z* is conditionally independent of its non-descendants given its parents. This is obvious for *Z* ∈ {*G*, *E*_1_, … , *E_p_*}; we now consider *ϒ_j_*. In 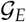, the set of parents of *ϒ_j_* is *pa*(*ϒ_j_*) ∪ {*E_j_*}, where *pa*(*ϒ_j_*) contains *G* if *j* ∈ *D*. By construction, *ϒ_j_* is entirely determined by *pa*(*ϒ_j_*) ∪ {*E_j_*}, and constant conditional on these variables. Consequently, given *pa*(*ϒ_j_*) ∪ {*E_j_*}, it is independent of any *E_k_* (*k* ≠ *j*), and of any *ϒ_k_* that it is a non-descendant of *ϒ_j_* (Note that if *G* ∉ *pa*(*ϒ_j_*), *ϒ_j_* is indeed conditionally independent of any non-descendant; if *G* ∈ *pa*(*ϒ_j_*), *G* cannot be the non-descendant because it is already in the conditioning set). Therefore the local Markov property holds for 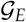. By the lemma below, we find that also the Markov factorization property holds for 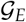, since for any distribution having a density it is equivalent with the local and global Markov properties. Given the Markov factorization property for 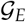 and the fact that 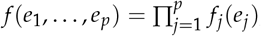 we can just integrate out the *e_j_*, and obtain the Markov factorization property for 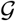. This concludes the proof of Theorem 1.

Given the result of Theorem 1 and the assumed faithulness, Theorem 2 now follows directly from the consistency for the general PC-algorithm (Spirtes *et al.* 2001); see the first part of the proof of their Theorem 5.1 (p. 407).

## Markov properties

The following lemma is taken from Lauritzen (1996) (p. 51), and reformulated with somewhat less general conditions, which however suffice for our purpose.

### Lemma

Let *P* be the joint distribution of random variables (*ϒ*_1_, … , *ϒ_p_*), having a density *f*, and let 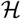 be a DAG on these variables. The following properties are equivalent:

- The Markov factorization property: given the parents *pa_j_* of each *x_j_*, the joint density (*f*) can be decomposed as

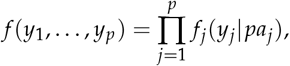

where the *f_j_* are the conditional densities.
- The local Markov property: any variable is conditionally independent of its non-descendants, given its parents.
- The global Markov property: for all disjoint sets *U*, *V*, *S* ⊂ {1, … , *ϒ_p_*}, d-separation of *U* and *V* by *S* in the graph 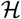 implies conditional independence of *U* and *V* given *S*. In contrast to *U* and *V*, the conditioning set *S* may be empty here. A definition of d-separation is given in Appendix A.1.

## A.10. Proof of Theorems 3 and 5

We first prove Theorem 3, by showing the equivalence of the left- and right-hand sides of equations (9) and (11) in the main text. The d-separation statements on the right-hand sides are equivalent, as *G* can never be a (descendant of a) collider. Also the left-hand sides (**ϒ_j_** ╨ **ϒ_k_**|{**ϒ_S_**, **G**} and 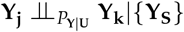) are equivalent, since

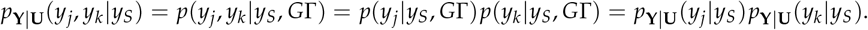

For Theorem 5 we make the additional assumption that *K* = *ZZ*^*t*^, *Z* = *I*_*m*_ ⊗ (1, … , 1)^*t*^ being the *mr* × *m* design matrix for *r* replicates of *m* genotypes in a balanced design (with *mr* = *n*). The first part of Theorem 5 then follows from the results in Appendix A.8. For the second part, we first recall the equivalence of **ϒ_j_** ╨ **ϒ_k_**|{**ϒ_S_**, **G**} and 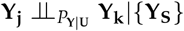. Because of the Gaussianity and the assumed faithfulness, the latter conditional independence is equivalent with

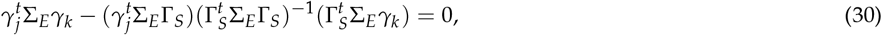

where we used (21).

Next we consider the conditional distribution of 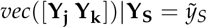 given in () (main text), whose covariance is the 2*n* × 2*n* block matrix 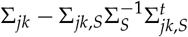. All *n* × *n* blocks are a linear combinations of *K* and *I*_*n*_, and it suffices to show that the coefficient of *I*_*n*_ in the off-diagonal blocks is zero if and only if (30) holds. We recall from (26) that

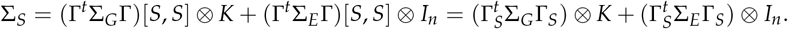

Using the Woodbury identity (equation (29)) with *A* = *V*_*E*_ ⊗ *I*_*n*_, *B* = *V*_*G*_ ⊗ *I*_*m*_ and *C* = *I*_*p*_ ⊗ *Z*, it follows that for any positive-definite *p* × *p* matrices *V*_*G*_ and *V*_*E*_, we have

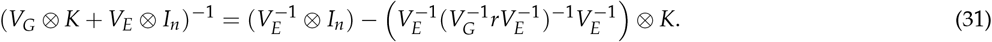

Setting 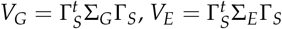 and 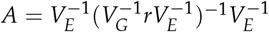, it follows that

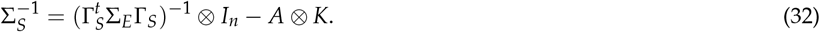

Combining this with the expressions for Σ_*jk*_ and Σ_*jk*,*S*_ given in Appendix A.7, we find that 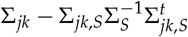 has off-diagonal blocks

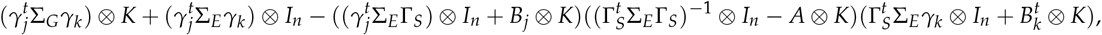

for *B*_*j*_ = *γ_j_* Σ_*G*_ Γ_*S*_ and *B*_*k*_ = *γk* Σ_*G*_ Γ_*S*_. Finally, working out the products in the last display (using that *K*^2^ = (*ZZ*^*t*^)^2^*rK*), we find that all terms involving a kronecker product with *I_n_* correspond exactly to the left-hand side of (30). Consequently, the residual covariance in the distribution 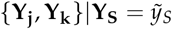 is zero if and only if **ϒ_j_ ╨ **ϒ_k_**|{**ϒ_S_**, **G**}.**

## A.11. Proof of Theorem 4

To obtain faithfulness for *S* = Ø, we need to prove that **ϒ_j_** ╨ **G** implies d-separation of *ϒ_j_* and *G* in the graph 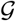. Because the conditioning set is empty, it suffices to show that there are no directed paths from *G* to *ϒ_j_*, where we can assume that *j* ∉ *D* (otherwise **G_j_** would be nonzero, and because of the non-collinearity, **ϒ_j_** and **G** would not be independent). Because of the assumed Gaussianity, the independence of **ϒ_j_** and **G** implies that

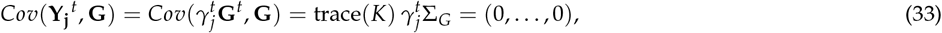

where we used that *vec*(**G**) ~ *N*(0, Σ_*G*_ ⊗ *K*), and therefore *E*(**G**[*i*, *j*]**G**[*i*, *k*]) = Σ_*G*_ [*j*, *k*]*K*[*i*, *i*], for all *i* ∈ {1, … , *n* and *j*, *k* ∈ {1, … , *p*. Since trace(*K*) is strictly positive and the submatrix Σ_*G*_ [*D*, *D*] has full rank, equation (33) implies that *γ*_*j*,*l*_ = 0 for all *l* ∈ *D*. Finally, we use that the assumed faithfulness implies the path coefficients condition (see Appendices A.3-A.6). Consequently, it follows from *γ*_*j*,*l*_ = 0 that there is no directed path from *ϒ*_*l*_ to *ϒ*_*j*_. Since this is the case for all *l* ∈ *D*, there can neither be a directed path from *G* to *ϒ*_*j*_.

## A.12. Proof of Theorem 6

Assuming *K* = *ZZ*^*t*^, the first part of theorem follows from the results in Appendix A.8. For the second part, we use that **ϒ_j_** has genetic variance 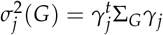 (see equation (20)). Because for traits without a direct genetic effect, rows and columns in Σ_*G*_ are zero, we can rewrite this as 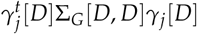. Hence, 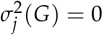 is equivalent with *γ*_*j*,*l*_ = 0 for all *l* ∈ *D*, where we used that Σ_*G*_ [*D*, *D*] is of full rank (which is a consequence of assumption 5). Using the arguments from the proof of Theorem 4 and the assumed faithfulness, it follows that this is equivalent with independence of **ϒ_j_** and **G**.

## Supplementary file 1. PCgen: implementation details

### 1.1. The PC-stable algorithm

We first state the PC-stable algorithm of Colombo and Maathuis (2014), which forms the basis of PCgen. As the original PC-algorithm, PC-stable has a skeleton and orientation stage; the former is described separately as Algorithm 2 below. Instead of updating the skeleton directly after each conditional independence test (as the original PC-algorithm), PC-stable only updates the skeleton list-wise, after doing all tests with *Y*_*S*_ of a given size |*S*| = *s*. More specifically, lines 7-8 in Algorithm 2 make an inventory of the current adjacency sets, which determines which tests of a given size *s* are to be conducted. Also the orientation rules R1-R3 in Algorithm 1 are applied listwise.

Since edges are not directly removed after finding conditional independence, multiple separation sets may be found for a given pair of variables. These may lead to conflicts in the orientation, for example when there are conflicting v-structures *ϒ*_*j*_ → *ϒ*_*k*_ ← *ϒ*_*l*_ and *ϒ*_*k*_ → *ϒ*_*l*_ ← *ϒ*_*m*_. Whenever possible these conflicts are resolved using the majority rule (line 6 of Algorithm 1). Unresolved conflicts are represented with an undirected edge^11^. In some cases this may lead to partially directed graphs that are not a CPDAG, but the advantages are that PC-stable is order-independent and easy to parallelize.

**Algorithm 1.**
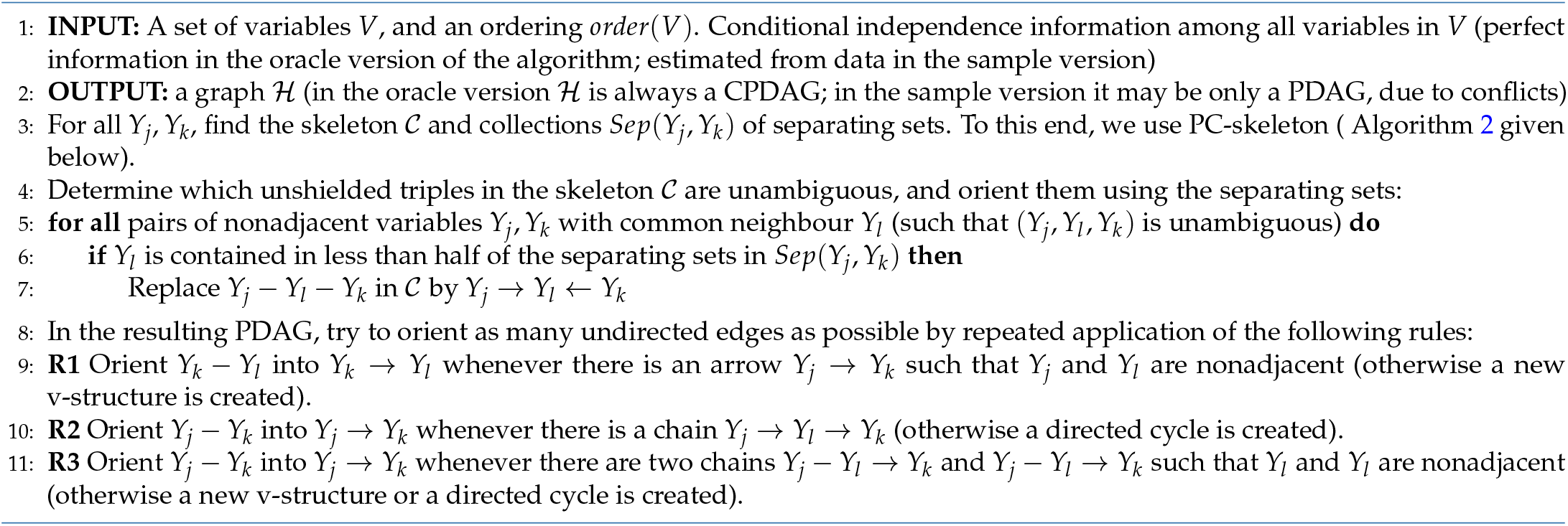
The PC-stable algorithm (following the description of Colombo and Maathuis (2014), and adapted to our notation).

**Algorithm 2.**
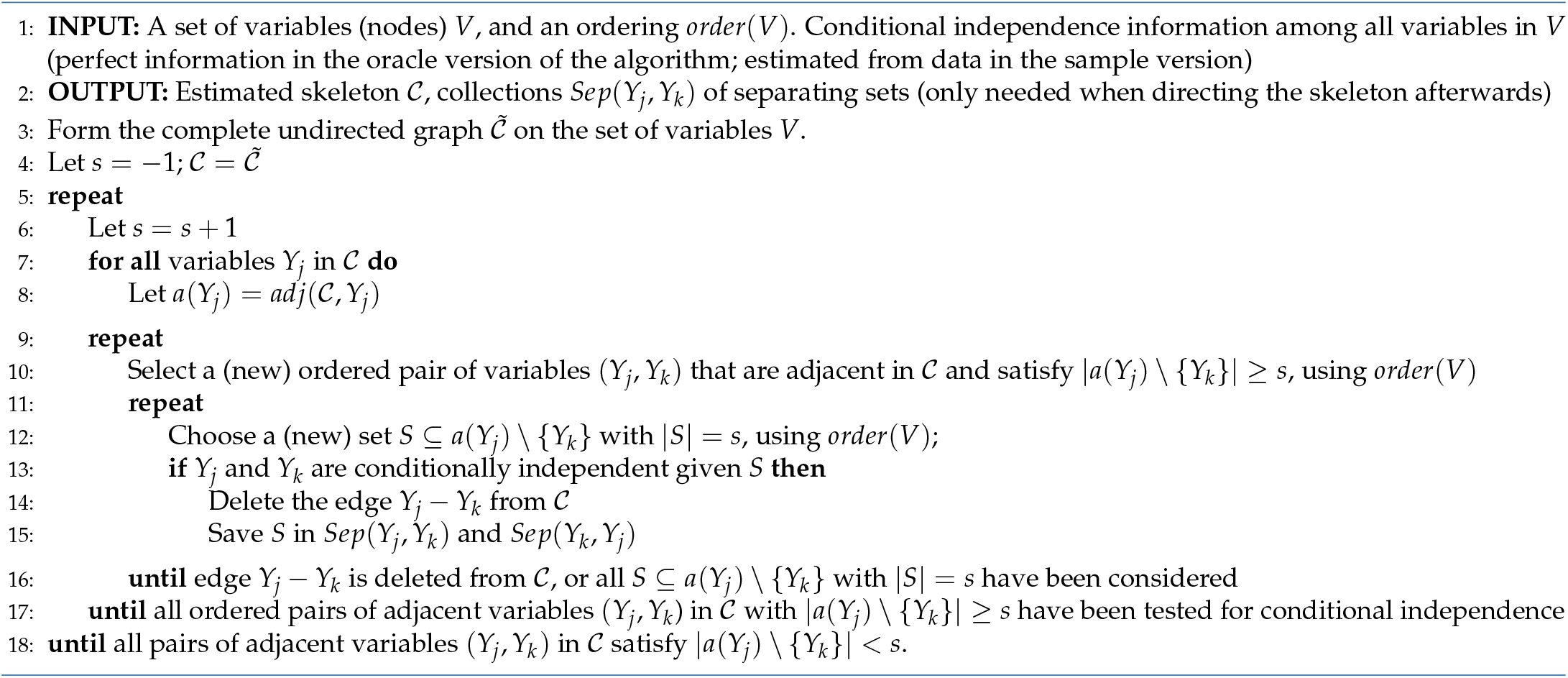
PC-skeleton (following the description of Colombo and Maathuis (2014), and adapted to our notation).

### 1.2. Modified orientation rules

PCgen follows the usual orientation rules of the PC-stable algorithm (lines 9-11 in Algorithm 1), except for the following modifications, which are required to avoid arrows pointing into *G*. Note that Algorithm 1 is written in generic notation with nodes *ϒ*_1_, … , *ϒ*_*p*_; in PCgen we have *ϒ*_1_, … , *ϒ*_*p*+1_, corresponding to *G*, *ϒ*_1_, … , *ϒ*_*p*_.

- in line 5 in Algorithm 1, we skip those triples where *ϒ*_*k*_ turns out to be *G*
- after line 7, we orient all remaining undirected edges *G* − *ϒ*_*j*_ as *G* → *ϒ*_*j*_.

These changes appear to be necessary, as edges *G ← ϒ_j_* cannot be avoided with the fixedEdges argument in the pc-function of the pcalg-package (Kalisch *et al.* 2012), where one can only enforce the presence of an edge in the skeleton, but not its orientation. pcalg also has the addBgKnowledge option to add orientations (’background knowledge’) in the estimated CPDAG, but this is only done *after* running the PC-algorithm, and is only allowed if compatible with the CPDAG. Here we intend to alwaY_S_ enforce the orientation *G* → *ϒ_j_*, and include it in the causal inference algorithm.

### 1.3. The RC-test for Y_j_ ╨ Y_k_|{G, Y_S_}

In the RC-test for **ϒ_j_** ╨ **Y_k_** |{**G**, **Y_S_**}, the null-hypothesis is that the residual covariance is zero, where the residual covariance is the off-diagonal element of *V*_*G*_ (*jk*|*S*), in equations () and (13) in the main text. The residual likelihood ratio test (RLRT) statistic for this hypothesis is defined as twice the difference in residual log-likelihood between the full and reduced model. As the null-hypothesis is not on the boundary of the parameter space, the distribution of the RLRT is approximately chi-square with 1 degree of freedom. Residual Maximum Likelihood (REML) estimates for the full and reduced model are obtained using an EM-algorithm Dempster *et al.* (1977). To improve efficiency, we use the following computational shortcuts:

1. For the reduced model, we compute starting values for the genetic (co) variances and residual variances, using multivariate analysis of variance (MANOVA), and then fit the model using the EM-algorithm.
2. For the full model, we take as starting values the estimates found for the reduced model. At each iteration of the EM-algorithm, we compute a preliminary RLRT p-value on the basis of the current restricted log-likelihood and the restricted log-likelihood of the reduced model, and stop the EM-algorithm if the p-value is below the significance threshold. This is possible because the EM-algorithm alwaY_S_ increases the likelihood at every iteration, while PCgen only requires an accept/reject decision. Of course, if exact p-values are required the EM-algorithm needs to be run until convergence.
3. We set a maximum number of 50 EM-iterations for the full model and 5 for the reduced model. Given the good starting values this usually sufficient, but occasionally EM would otherwise take very many iterations until convergence. Usually, in these cases, the RLRT statistic obtained with an unrestricted number of EM-iterations is not significant, meaning that stopping EM earlier rarely affects the outcome of the RLRT.

### 1.4. Testing the genetic covariance

Assuming that the joint distribution of *vec*([**Y_j_ Yk**]) has covariance *V*_*G*_ (*jk*) ⊗ *K* + *V*_*E*_ (*jk*) ⊗ *I*_*n*_ (i.e., (13) in the main text, with empty conditioning set), it is often of interest to test for absence of genetic covariance, i.e., whether the off-diagonal element of *V*_*G*_ (*jk*) is zero. Although not part of PCgen, this can be tested similar to the RC-test described above. Alternatively, we could test whether the total genetic *correlation* is zero, using the delta method as in Korte *et al.* (2012).

Again we define a RLRT statistic as twice the difference in residual log-likelihood between the full and reduced model, where the latter is restricted to have diagonal *V*_*G*_ (*jk*). As before the distribution of this RLRT is approximately chi-square with 1 degree of freedom, and REML-estimates for the full and reduced model are obtained with the same EM-algorithm used earlier. In the pcgen package this is implemented in the function `gencorTest`.

## Supplementary file 2. Overview of algorithms

PCgen can be run with either the RC- or the RG-test. Moreover, in PCgen as well as PCres one can use observations on replicates (if available), a GRM (typically with means), or both a GRM and replicates. Together this leads to a large number of options, which we could not all implement. Tables 2 and 3 below give an overview of the algorithms implemented in our pcgen package. We now give a brief overview, with an explanation of the acronyms used. Note that for the sake of readibility we used ‘PCgen’, ‘PCres(means)’ and ‘PCres(replicates)’ in the main text, to refer to respectively PCgen-screening, PCres-multi-A and PCres-uni-R; see the tables below.

PCgen-RC and PCgen-RG refer to PCgen with respectively the RC- or the RG-test for conditional independencies of the form (B) (the test for (A) is always the same). In case of the RG-test, either univariate or multivariate GBLUP can be used (PCgen-RG-uni versus PCgen-RG-multi). Alternatively, we can run PCgen-RC starting with a skeleton determined by PCgen-RG-uni, as described in File S3; we will refer to this variant as PCgen-RC-screening. All currently implemented variations of PCgen require data on replicates, as we did not implement the likelihood ratio test for **Y_j_** ╨ **G**|**Y_S_**(described in the main text).

PCres can be based on either univariate or multivariate GBLUP (PCres-uni versus PCres-multi). In either case, one can use an (additive) relatedness matrix (-A), replicates (-R), or both replicates and the relatedness matrix (-RA). If only a relatedness matrix is used, genotypic means are required. Alternatively, one could reconstruct 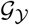 using the GBLUP itself (PC-GBLUP) itself, similar to the approach of Töpner *et al.* (2017).

Finally, note that if we have *r* replicates of *m* genotypes in a completely randomized design, with an *m × m* relatedness matrix *A*, the distribution of the data is

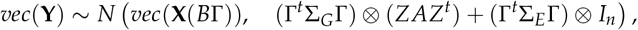

i.e., equation (7) (main text) with *K* = *ZAZ*^*t*^. Then the distribution of the *m* × *p* matrix 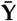 of genotypic means is then given by

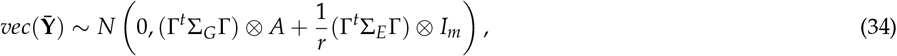

where it is assumed that the fixed effects have been accounted for in the estimation of genotypic means. Consequently, the means are distributed according to the same GSEM, but with *r* times fewer observations, and residual (co)variances that are *r* times as small.

**Table 2.**
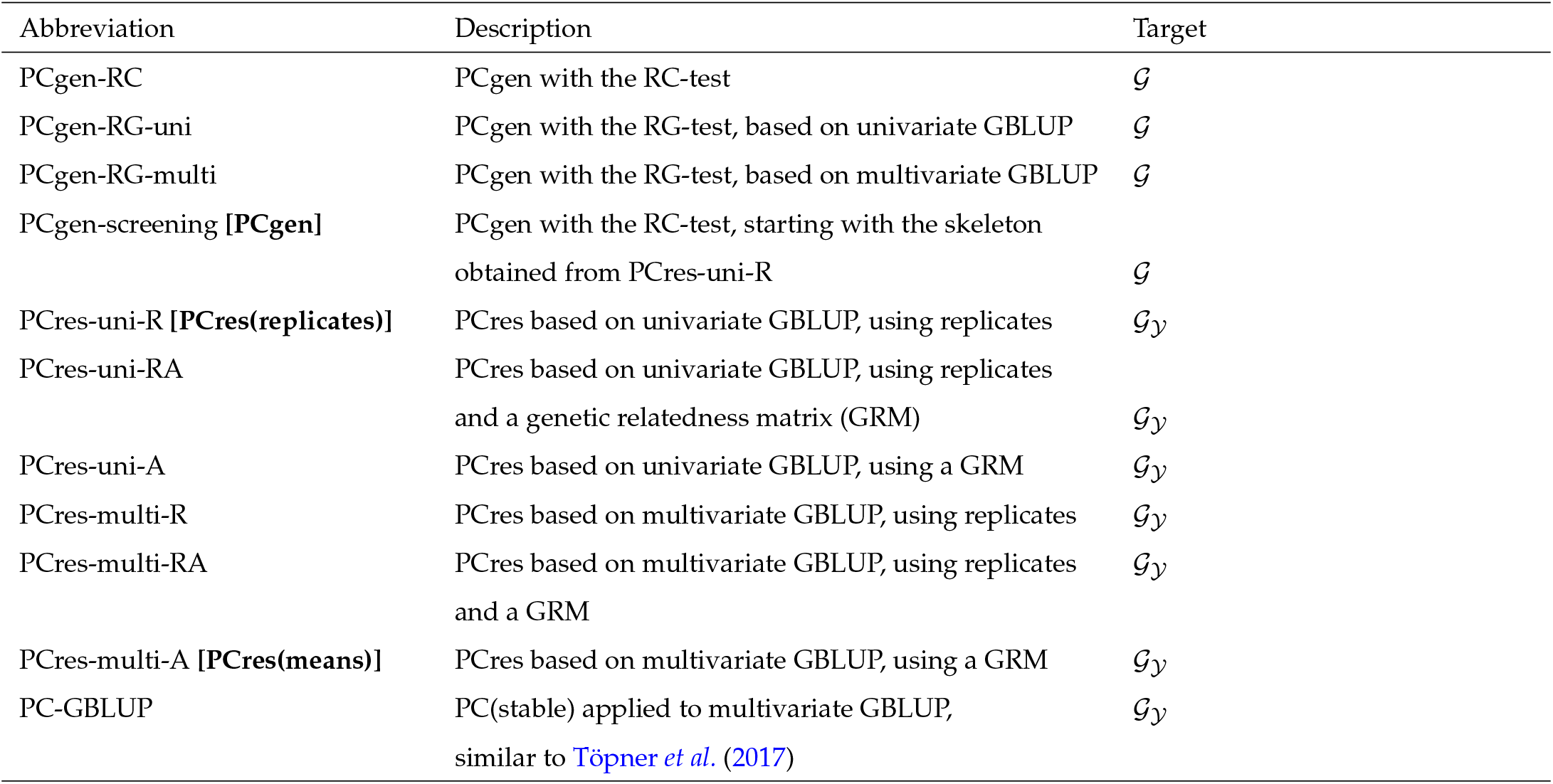
Overview of the algorithms available in the pcgen package, for reconstructing either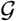 (the complete graph) or 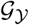 (the subgraph of trait-to-trait relations). The required commands in the pcgen package are given in Table 3 below. The abbreviations ‘PCgen’, ‘PCres(means)’ and ‘PCres(replicates)’ used in the main text refer to respectively PCgen-screening, PCres-multi-A and PCres-uni-R. All PCres approaches are based on the residuals of GBLUP, while PC-GBLUP uses the GBLUP itself.

| Abbreviation | Description | Target |
| --- | --- | --- |
| PCgen-RC | PCgen with the RC-test | $\mathcal{G}$ |
| PCgen-RG-uni | PCgen with the RG-test, based on univariate GBLUP | $\mathcal{G}$ |
| PCgen-RG-multi | PCgen with the RG-test, based on multivariate GBLUP | $\mathcal{G}$ |
| PCgen-screening [PCgen] | PCgen with the RC-test, starting with the skeleton obtained from PCres-uni-R | $\mathcal{G}$ |
| PCres-uni-R [PCres(replicates)] | PCres based on univariate GBLUP, using replicates | $\mathcal{G}_y$ |
| PCres-uni-RA | PCres based on univariate GBLUP, using replicates and a genetic relatedness matrix (GRM) | $\mathcal{G}_y$ |
| PCres-uni-A | PCres based on univariate GBLUP, using a GRM | $\mathcal{G}_y$ |
| PCres-multi-R | PCres based on multivariate GBLUP, using replicates | $\mathcal{G}_y$ |
| PCres-multi-RA | PCres based on multivariate GBLUP, using replicates and a GRM | $\mathcal{G}_y$ |
| PCres-multi-A [PCres(means)] | PCres based on multivariate GBLUP, using a GRM | $\mathcal{G}_y$ |
| PC-GBLUP | PC(stable) applied to multivariate GBLUP, similar to Töpner *et al.* (2017) | $\mathcal{G}_y$ |

## Supplementary file 3. Extensions of PCgen

We considered the following extensions of PCgen, most of which are implemented in the **pcgen**-package:

- **Prior screening (PCgen-screening):** The RC-test for statement (B) is computationally efficient, in the sense that only a bivariate MTM needs to be fitted, instead of an MTM for all traits. At the same time however, once residuals are available, the RG-test is much faster than the RC-test, as the former is based on partial correlations that can be computed recursively (see e.g Kalisch and Bühlmann (2007)). While residuals from the full MTM (for all traits) are difficult to obtain, residuals from single-trait GBLUP can be computed very easily. An appealing strategy, therefore, is to run PC-stable on univariate residuals (i.e., a form of PCres), and use the resulting skeleton as the starting point for PCgen with the RC-test. The advantage, at least for sparse graphs, is that the number of conducted RC-tests is greatly reduced. In the pcgen package this is implemented in the pcgenFast function. The skeleton based on univariate residuals typically contains somewhat more false edges, but these may be removed later on with the RC-test.
- **Inclusion of QTLs:** apart from the random genetic effects, the GSEMs considered here could include fixed effect QTLs as well. Each QTL is represented by a single node, which like the node *G* is always a root node. No edges are allowed between QTLs or between a QTL and *G*, and every edge between a QTL and a trait is oriented towards the trait. The QTLs may further improve the orientation of the graph, but require different types of conditional independence tests, which is left for future work.
- **Comparing PCgen output with genetic variance estimates:** for every trait **Y_j_** having positive genetic variance, there should be either a direct genetic effect *G* → *ϒ*_*j*_ or a partially directed path from *G* to *ϒ_j_* (with possibly undirected edges, but all directed edges pointing towards *ϒ*_*j*_). However, because of statistical errors it may happen that neither of the two exist in the CPDAG obtained from PCgen, while at the same time the genetic variance (considered in the independence test **Y_j_** ╨ **G**) is significantly different from zero. In the pcgen package, such conflicts can be detected using the checkG function. If conflicts occur, we conclude that there was insufficient evidence to remove the direct genetic effect from the graph, and re-run PCgen, skipping all tests **Y_j_** ╨ **G**|**Y_S_** for all **Y_j_** that produced conflicts in the first run. This forces the algorithm to keep the edge *G* → *ϒ*_*j*_.
- **Restricting the maximum size of *S*:** As in the original PC-algorithm, the conditioning sets *S* considered in PCgen can be restricted to a pre-specified maximum size. If the only set that d-separates two traits *ϒ*_*j*_ and *ϒ*_*k*_ is bigger than this size, the reconstructed graph will contain the false edge *ϒ*_*j*_ − *ϒ*_*k*_ (unless a statistical error occurred in one of the earlier tests, with smaller *S*); hence the graph will be somewhat denser than without the restriction. Nevertheless, setting a maximum size may have substantial computational and statistical advantages. This is especially the case when testing **Y_j_** ╨ **G**|**Y_S_**, and many traits have a direct genetic effect, making the adjacency set of the node *G* very large. Without any restriction, the number of tests would then increase exponentially with the number of traits. This problem is less severe for PCres; for large datasets we therefore propose to use the PCgen-screening approach described above, with no or little restriction in the screening step. We will do this below for the larger simulated data sets and the rice data.

**Table 3.** R-commands for the different algorithms, as implemented in the package pcgen. The first argument is the required phenotypic data-frame (suffStat = d (replicates) or suffStat = m (genotypic means)). The dots represent generic arguments (e.g., alpha and m.max, which define the significance threshold and the maximum size of the conditioning sets). cov.method determines whether univariate (uni) or multivariate (us) GBLUP is to be used (“us” stands for unstructured, as opposed to e.g., factor analytic models. All algorithms involving GBLUP use the residuals (use.GBLUP = F), except the genomic network similar to Töpner *et al.* (2017) (PC-GBLUP, with use.GBLUP = TRUE). Finally, A is a genetic relatedness matrix, which can be included by putting K = A; otherwise the default is used (K = NULL, in which case replicates are required). The abbreviations ‘PCgen’, ‘PCres(means)’ and ‘PCres(replicates)’ used in the main text refer to respectively PCgen-screening, PCres-multi-A and PCres-uni-R.

| Abbreviation | Commands in `pcgen` package |
| --- | --- |
| PCgen-RC | `pcgen(d,..., use.res = FALSE)` |
| PCgen-RG-uni | `C <- cor(getResiduals(d,..., cov.method = "uni"))`<br>`pcgen(d,..., use.res = TRUE, res.cor = C)`<br>equivalently: `pcgenFast(..., use.res = TRUE)` |
| PCgen-RG-multi | `C <- cor(getResiduals(d,..., cov.method = "us"))`<br>`pcgen(d,..., use.res = TRUE, res.cor = C)` |
| PCgen-screening [PCgen] | `pcgenFast(d,..., use.res = FALSE)` |
| PCres-uni-R [PCres(replicates)] | `pcRes(d,..., cov.method = "uni", use.GBLUP = FALSE)` |
| PCres-uni-RA | `pcRes(d,..., cov.method = "uni", use.GBLUP = FALSE, K = A)` |
| PCres-uni-A | `pcRes(m,..., cov.method = "uni", use.GBLUP = FALSE, K = A)` |
| PCres-multi-R | `pcRes(d,..., cov.method = "us", use.GBLUP = FALSE)` |
| PCres-multi-RA | `pcRes(d,..., cov.method = "us", use.GBLUP = FALSE, K = A)` |
| PCres-multi-A [PCres(means)] | `pcRes(m,..., cov.method = "us", use.GBLUP = FALSE, K = A)` |
| PC-GBLUP | `pcRes(m,..., cov.method = "us", use.GBLUP = TRUE, K = A)` |

## Supplementary file 4. Simulations

### 4.1. Simulation setup

We simulate data from model (4) (main text) using the following steps, closely resembling the simulations of Kalisch and Bühlmann (2007):

- Given the required number of traits (*p*) and sparseness of the graph (defined by the parameter *p_t_* below), we first generate the *p* × *p* matrix Λ, which determines the structural relations among the traits. Λ is simulated using the `randomDAG` function from the R-package pcalg (Kalisch *et al.* 2012), where edges (i.e., the nonzero elements of Λ) occur with probability *p*_*t*_ ∈ (0, 1) (for details see the pcalg-documentation and Kalisch and Bühlmann (2007), p. 621). The expected number of neighbors of each node is then *p*_*t*_ (*p* − 1). The values of the nonzero coefficients are drawn independently from the uniform distribution on [0.5, 1] and then given a random sign.
- The DAG defined by Λ is now extended with a genetic node *G*. For a proportion of *p*_*g*_ ∊ (0, 1) of the traits, we add an edge *G* →*ϒ*_*j*_. The subset of traits *D* for which there is a direct genetic effect then contains *p*. *p*_*g*_ traits. These are always chosen to be the traits of highest topological order (in the initial DAG defined by Λ). For example, if *p* = 4 and *p*_*g*_ = 0.5, and the initial DAG is *ϒ*_1_ → *ϒ*_2_ →*ϒ*_3_→ *ϒ*_4_, *D* will consist of *ϒ*_1_ and *ϒ*_2_.
- Next we simulate the corresponding genetic variances and covariances in Σ_*G*_. The genetic variances are drawn independently from a uniform distribution on [1, 2] and random covariances are introduced through random eigenvectors as in Joe (2006), using the `genPositiveDefMat` function from the `R-package clusterGeneration`.
- Given the relatedness matrix (*K*) and the required numbers of genotypes (*n*) and replicates (*r*), the direct genetic effects (**G**) are drawn from the matrix-variate normal distribution with column covariance *ZKZ*^*t*^ and row-covariance Σ_*G*_.
- Similarly, the residual effects (**E**) are drawn from the matrix-variate normal distribution with column covariance *I*_*nr*_ and row-covariance *I*_*p*_. Although the residual variance is 1 for all traits, the heritability of the traits still varies, as the variances of the direct genetic effects are between 1 and 2. Traits without a direct genetic effect typically have heritability below 0.5.
- Given the matrices **G**, **E** and Λ obtained in the previous steps, **Y** is computed using (4) in the main text. This is done recursively, following the topological ordering of the DAG (Kalisch and Bühlmann (2007), p. 621).

### 4.2. Performance criteria

As in Kalisch and Bühlmann (2007) we compare the true (simulated) CPDAGs and the estimated CPDAGs. Instead of considering the complete CPDAG, we evaluate separately the subgraph defined by Λ (relations among traits) and the direct genetic effects (the subgraph with edges *G*_*j*_ → *ϒ*_*j*_, whenever *j* ∈ *D*). For both subgraphs we consider the following criteria, again following Kalisch and Bühlmann (2007):

- True Positive Rate (TPR): the number of correct edges (in the estimated skeleton) divided by the total number of true edges (in the true skeleton).
- False Positive Rate (FPR): the number of incorrect edges (in the estimated skeleton) divided by the total number of true gaps (in the true skeleton).
- True Discovery Rate (TDR): the number of edges in the estimated graph that are correct (i.e., exist in the true skeleton) divided by the total number of edges in the estimated graph.
- Structural Hamming Distance (SHD): the number of edge deletions, additions and flips required to transform the estimated CPDAG into the true CPDAG. See also Tsamardinos *et al.* (2006).

All criteria were computed using functions from the pcalg-package (the TPR and FPR using the compareGraphs function, and the SHD using the function shd).

## Supplementary file 5. Simulation with APSIM

We used the crop-growth model APSIM-wheat to simulate 12 traits (**ϒ**_**1**_, … , **ϒ**_**12**_) for an existing population of 199 wheat genotypes, characterized with 3,035 SNPs with minor allele frequency larger than 0.05 (details in Bustos-Korts (2017)). APSIM simulations were carried out in Emerald, during 1993, which corresponds to a severe drought environment. Simulation settings were the same as in Casadebaig *et al.* (2016) and genotype-specific parameters had the ranges specified in Bustos-Korts (2017). The wheat panel was assumed to segregate for seven of the APSIM parameters, which we refer to as the primary traits. These are the only ones for which there are direct genetic effects. The seven primary traits (**ϒ**_**1**_, … , **ϒ**_**7**_) are a subset of those used in Bustos-Korts (2017), and were chosen because they have an important impact on grain yield, as shown by global sensitivity analysis (Casadebaig *et al.* 2016). Apart from the primary traits there are four secondary traits, each of them depending on some of the primary traits, and sometimes some of the other secondary traits. The final trait is yield, which depends on three of the secondary traits. Acronyms for all 12 traits are given in Table S6. For each genotype three replicates were simulated.

The direct genetic effects on the 7 primary traits were simulated as the sum of 300 additive QTL-effects. Different samples of 300 SNPs were used for each trait, and each effect was sampled from a trait-specific Gamma distribution. The shape and rate of this distribution were obtained by fitting a Gamma distribution to empirical additive effects estimated in a GWAS analysis of real phenotypes, observed for this population in the Australian wheat belt. For the phenology-related traits SV and SP we set *k* = 0.7 and *b* = 13.6, and *k* = 1.3 and *b* = 13.6 for the other primary traits. We then added Gaussian noise, to get a heritability of 0.9 for all primary traits.

The secondary traits (**ϒ**_**8**_, … , **ϒ**_**11**_) and yield (**ϒ**_**12**_) were simulated by running a dynamic model from time zero to time *T*, the time-point at which all traits are observed:

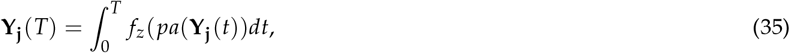

where *pa*(**ϒ**_**j**_)(*t*) are the values of the ‘parent traits’ at time-point *t*, and *z* represents a set of fixed parameters, specific for the environment under consideration. The form of *f*_*z*_ is detailed in Bustos-Korts (2017). The sets of parental traits stay the same over time, and therefore define the summary graph given in Figure S1 (main text).

## Supplementary file 6. Skipping conditional independence tests that do not involve *G*

### 6.1. Skipping tests for statements of the form **ϒ**_**j**_ ╨ **ϒ**_**k**_|{**ϒ**_**S**_}: a motivation

In Theorem 2 (main text) we provided conditions for consistency of the oracle version of PCgen. In our experience, the finite sample performance of PCgen can be improved if we skip some of the tests in the skeleton-stage. Differences between the oracle and sample version of PCgen can occur everywhere in the graph, but seem to occur most often for conditioning sets not containing *G*. This is illustrated in the example in Figure 8, in which there are genetic effects on traits *ϒ*_1_ and *ϒ*_2_, as well as a direct effect of *ϒ*_1_ on *ϒ*_2_.

**Figure. 8.**
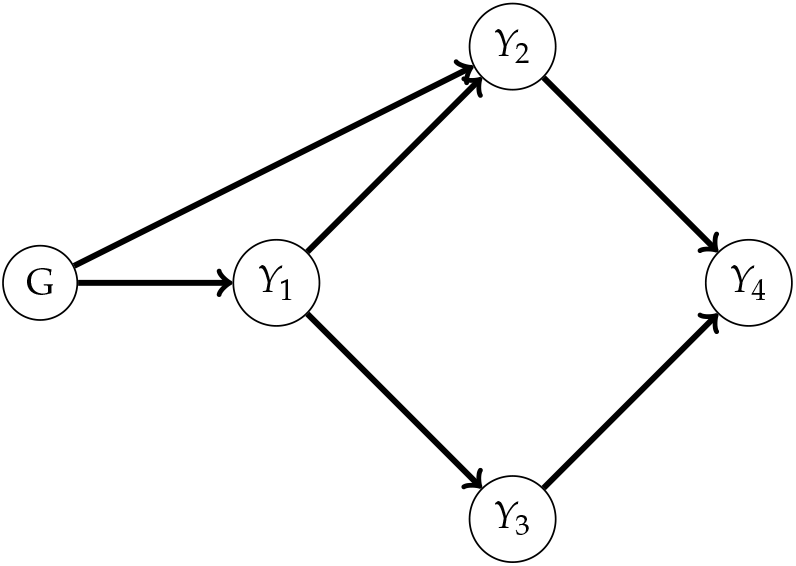
A GSEM with independent genetic effects on *ϒ*_1_ and *ϒ*_2_, and a direct effect *ϒ*_1_ → *ϒ*_2_.

**Figure. 9.**
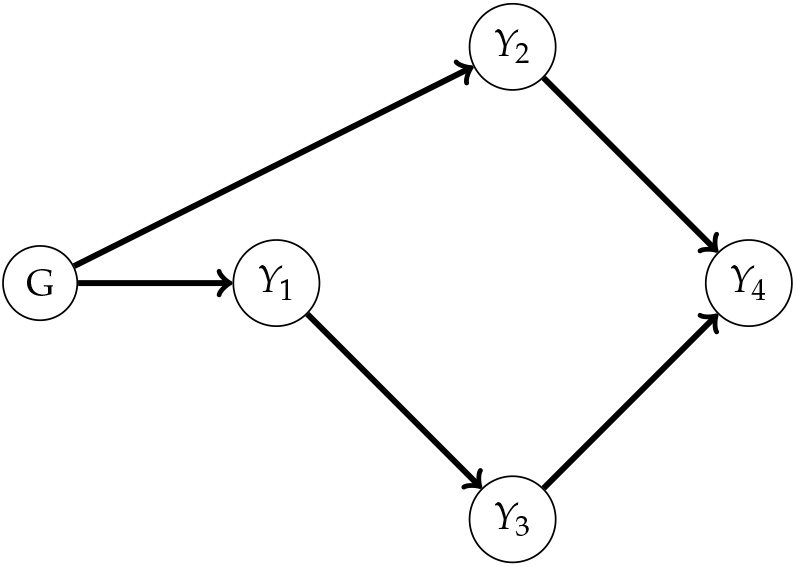
Typical output of PCgen, without skipping tests for **ϒ**_**j**_ ╨ ϒ_k_ | {ϒ_**S**_, and based on observations of (**ϒ**_**1**_, **ϒ**_**2**_, **ϒ**_**3**_, **ϒ**_**4**_) generated by the model of Figure 8, for 400 genotypes and 2 replicates. The edge between *ϒ*_1_ and *ϒ*_2_ is missing because the test for conditional independence of **ϒ**_**1**_ and **ϒ**_**2**_ given **ϒ**_**4**_ has too little power.

Then given a large number of observations and assuming faithfulness, PCgen will recover the true skeleton. However, with a small or moderate sample size, the test for conditional independence of **ϒ**_**1**_ and **ϒ**_**2**_ given **ϒ**_**4**_ has very little power. The test for conditional independence of **ϒ**_**1**_ and **ϒ**_**2**_ given {**ϒ**_**4**_, **G**} is a lot more powerful here. However, this test is not performed anymore, after the null-hypothesis of conditional independence of **ϒ**_**1**_ and **ϒ**_**2**_ given **ϒ**_**4**_ has been accepted. Therefore, a typical output of PCgen looks like Figure 9.

In order to make PCgen more powerful we therefore propose to perform only those conditional independence tests where **G** is contained in the conditioning set, at least when both variables whose conditional independence is tested have positive genetic variance. This still gives a valid algorithm, in the sense that the oracle version of PCgen gives identical output. Intuitively this is obvious, as *G* is a root node, and everything can be done conditionally on **G**. We formally show this below in section 6.3, for which we need the following well known characterization of the skeleton.

### 6.2. A characterization of the skeleton

If a distribution *P* is faithful with respect to a DAG 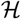, we have the following result for the skeleton of 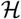: there is an edge between nodes *A* and *B* in the skeleton of a DAG 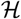

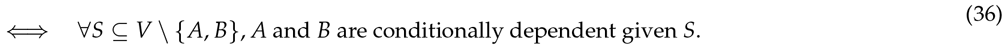

This was shown in Spirtes *et al.* (2001) (Theorem 3.4); here we adopt the formulation of Kalisch and Bühlmann (2007) (p.616). It is important to note that in general the skeleton is not equal to the so-called conditional independence graph (CIG), which is the undirected graph associated with the inverse covariance or precision matrix. The latter is characterized by an equivalence statement similar to (36), but with on the right-hand side only *S* = *V* \ {*A*, *B*}. Hence, if data are generated by a DAG 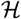 and we assume faithfulness, the skeleton of 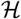 is typically a subgraph of the CIG. In case *A* → *B* ← *C* for example, the CIG also contains an edge *A* − *C* (because *S* = ∅ d-separates *A* and *C*, but *S* = *B* does not, *B* being a collider on the path *A* → *B* ← *C*).

### 6.3. *Skipping the test for* **ϒ**_**j**_ ╨ **ϒ**_**k**_|{**ϒ**_**S**_} does not affect PCgen (oracle)

In view of (36), our modification is correct in the sense that the oracle version of PCgen still recovers the true skeleton. This correctness follows from the facts that

1. like the PC-skeleton algorithm, PCgen starts with the complete undirected graph, and then tests conditional independencies, removing edges when a conditional independence relation is found. When in fact there *i*_*s*_ an edge *ϒ*_*j*_ – *ϒ*_*k*_ in the true graph 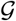, then of course skipping some of these tests still produces the correct result (and in the sample version of PCgen even with higher probability).
2. if there is *no* edge *ϒ*_*j*_ – *ϒ*_*k*_ and a set *S not* containing *G* is blocking a path between *ϒ*_*j*_ and *ϒ*_*k*_, then *{G}* ⋃ *S*} is also blocking it (since *G* can never be a collider, and neither a descendant of any node). In other words, it can not happen that a set *S* not containing *G* is blocking all paths between *ϒ*_*j*_ and *ϒ*_*k*_, and that the addition of *G* would ‘unblock’ one of these paths. Consequently, it can not happen that such a set *S* is the *only* set separating *ϒ*_*j*_ and *ϒ*_*k*_, and that we would miss the only opportunity to remove the edge between *ϒ*_*j*_ and *ϒ*_*k*_.

## Supplementary file 7. Miscelleneous

### 7.1. Representing **G** *by a single node: a motivating example*

The main reason for representing **G**_**1**_, … , **G**^**p**^ with a single node *G* (instead of multiple nodes *G*_1_, … , *G*_*p*_, sometimes used in the literature) is that the relatedness matrix *K* is the same for all traits. If for example *G*_1_ → *ϒ*_1_ → *ϒ*_3_ ← *ϒ*_2_ *G*_2_ (as in Figure 3 in the main text), it follows from (7) (also in the main text) that the marginal distribution of **ϒ**_**3**_has covariance *c*_1_ *K* + *c*_2_ *I*_*n*_, for some nonnegative constants *c*_1_, *c*_2_. However, based on this distribution alone, we cannot distinguish the contributions of **G**_**1**_ and **G**_**2**_. This differs from the scenario *QTL*_*A*_ → *ϒ*_*1*_ → *ϒ*_3_ ← *ϒ*_2_ ← *QTL*_*B*_, where the total (fixed) effects of QTLs *A* and *B* on *ϒ*_3_ can be estimated from the marginal distribution of *ϒ*_3_. If we condition on **ϒ**_**1**_and **ϒ**_**2**_, **ϒ**_**3**_becomes independent of **G**_**1**_ and **G**_**2**_. In fact it is also independent of **G**_**3**_, since the latter is zero. Consequently, given **ϒ**_**1**_ and **ϒ**_**2**_, **ϒ**_**3**_ is independent of **G**= [**G**_**1**_ **G**_**2**_ **G**_**3**_], which illustrates that the conditional independencies correspond to a property of the graph 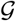, with [**G**_**1**_ **G**_**2**_ **G**_**3**_] represented by a single node *G*.

### 7.2. Genomic networks

Töpner *et al.* (2017) recently proposed to estimate a directed network based on the predicted genetic effects themselves, rather than the residuals. As mentioned in the Discussion (section ‘Data from different experiments’), the implicit assumption is that the GSEM model (equation (4) in the main text) is replaced by a model defined by **ϒ**= **X***B* + **U**+ **E** and **U** = **U**Λ + **Ũ**. This is however problematic if in fact the multi-trait data follow equation (4). For example, consider again the graph from Figure 3 in the main text (*ϒ*_1_ → *ϒ*_3_ ← *ϒ*_2_), with direct genetic effects on *ϒ*_1_ and *ϒ*_2_. For the sake of the argument, assume also that the total genetic effects can be predicted without error, i.e., we can observe the matrix **U**= **G**Γ (see equation (6) in the main text). Because *ϒ*_1_ and *ϒ*_2_ are root nodes in 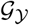, **U**_**1**_ = **G**_**1**_ and **U**_**2**_ = **G**_**2**_. We can make the following observations:

- To get the correct skeleton, the PC-algorithm must find that **Y**_**1**_ and **Y**_**2**_ are marginally independent. Using residuals, we indeed find that **Y**_**1**_ – **U**_**1**_ and **Y**_**2**_ – **U**_**2**_ are independent. However, **U**_**1**_ and **U**_**2**_ are only independent if Σ_*G*_ is diagonal, which is a rather strong and restrictive additional assumption.
- We may infer some of the edges *G* → *ϒ*_*j*_, but not all of them. Moreover, this can be done only indirectly, through comparison with the trait-to-trait network estimated by the residuals-based approach, and requires additional testing if the **U**_**j**_ are zero (similar to the tests for **ϒ**_**j**_ ╨ **G**|∅ in PCgen). In particular, if **U**_**j**_ = 0 for some *j*, this clearly excludes *G* → *ϒ*_*j*_. But in our example we find that **U**_**1**_, **U**_**2**_, **U**_**3**_ are all nonzero, and can only conclude (by comparing with the already known trait-to-trait network *ϒ*_1_ → *ϒ*_3_ ← *ϒ*_2_) that *G* → *ϒ*_1_ and *G* → *ϒ*_2_. It is impossible to make any inference about *G* → *ϒ*_3_, which PCgen can in principle do.

### 7.3. Intervening versus conditioning

Before considering genomic prediction in the next section, we first look at the difference between intervening and conditioning in an example without genetic effects. See the SEM in Figure 10, where there are no fixed effects, all path-coefficients are 1, and all error variables are standard normal and independent. Prior to the intervention **Y**_**2**_:= (1, … , 1)^*t*^, the joint distribution of (**Y**_**1**_[*i*], **Y**_**2**_[*i*], **Y**_**3**_[*i*])^*t*^ (for each individual *i*) is multivariate normal with mean zero and covariance Σ given by

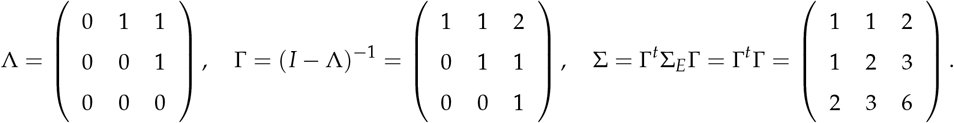

From well-known results for the multivariate normal distribution, it follows that the conditional distribution of **Y**_**3**_[*i*] given **Y**_**2**_[*i*] = 1 is normal with mean and variance both equal to 2. The conditional mean being larger than 1 may seem counterintuitive, but can be understood by noting that the information **Y**_**2**_[*i*] = 1 suggests a larger than exptected value for **Y**_**1**_[*i*], which in turn gives an additional (expected) effect on **Y**_**3**_[*i*]. In case we intervene and **Y**_**2**_[*i*] is forced to have a value 1, there is no effect *ϒ*_1_ → *ϒ*_2_, and the distribution of **Y**_**3**_[*i*] is normal with variance 2 and mean 1.

**Figure. 10.**
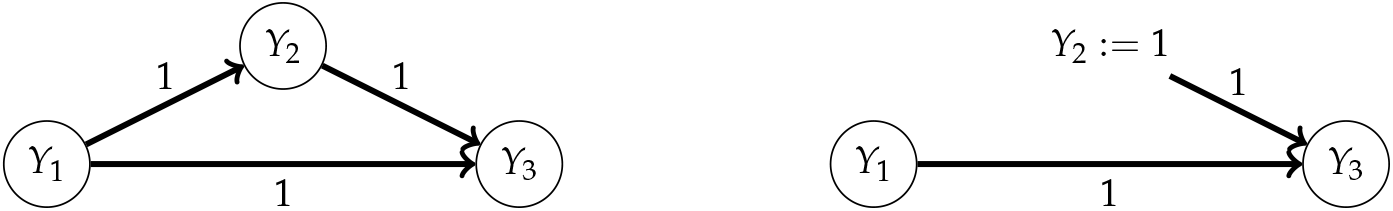
An example of a linear SEM, before and after the intervention *ϒ*_2_ := 1.

### 7.4. Intervening versus conditioning: genomic prediction

#### Simulation setup

Next we modify the preceding example and add genetic effects: see the SEM in Figure 11. For example, *ϒ*_1_, *ϒ*_2_ and *ϒ*_3_ could respectively be the sensitivity to photoperiod, flowering time and yield. There is no effect *ϒ*_1_ → *ϒ*_3_, and there are independent genetic effects on *ϒ*_1_ and *ϒ*_3_ with variance 1.5 and 4, respectively. The error variances of *ϒ*_1_, *ϒ*_2_ and *ϒ*_3_ (i.e., the diagonal of Σ_*E*_) are respectively 2, 1.5 and 1. Prior to the intervention **Y**_**2**_:= (1, … , 1)^*t*^, the joint distribution of (**Y**_**1**_, **Y**_**2**_, **Y**_**3**_)^*t*^ is defined by equation (7) in the main text, with mean zero and genetic and residual covariance defined by

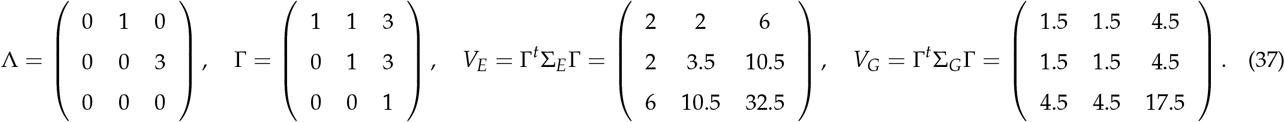

**Figure. 11.**
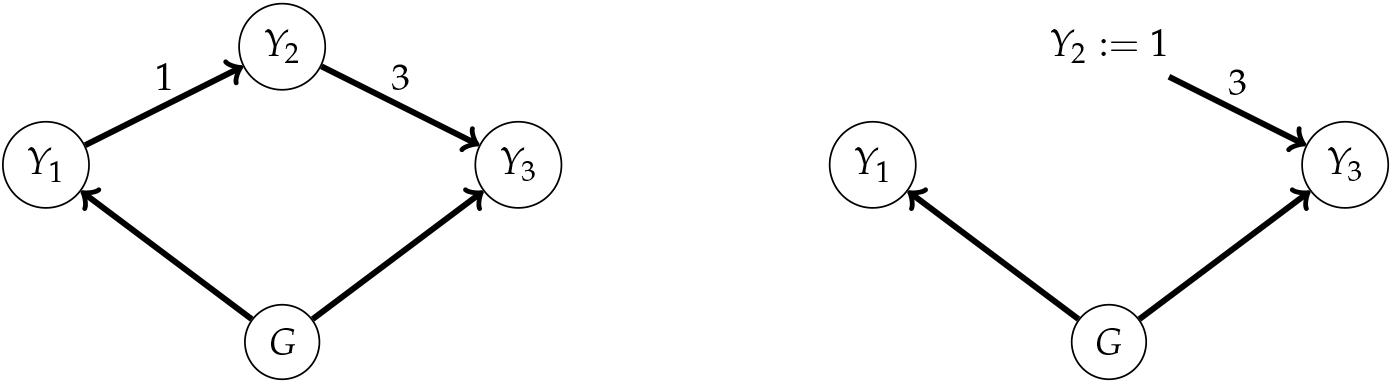
A linear GSEM, before and after the intervention *ϒ*_2_ := 1.

- We simulated data for the RegMap population (Horton *et al.* 2012), containing 1307 accessions of *Arabidopsis thaliana*. Two replicates were simulated for each accession.
- 200 datasets were simulated according to the GSEM prior to the intervention **Y**_**2**_:= 1, and 200 according to the GSEM after the intervention (see Figure 11). Additive genetic effects for *ϒ*_1_ and *ϒ*_3_ were simulated with variance given above, based on *q* = 100 randomly selected SNPs with minor allele frequency of at least 0.1. The same genetic effects were used before and after the intervention, whereas the residual errors were (of course) simulated separately. We used the RegMap in order to have a large population and used only 100 SNPs to speed up computation, but this is not essential.
- All genomic predictions were computed at the level of genotypic means. Note that, being the average of two replicates, the accession means have residual (co)variances that are twice as small as for the original observations (see equation (34) in File S2). For example, *ϒ*_1_ has a (line-) heritability of 1.5/(1.5 + 2/2) = 0.6.
- The 1307 accessions were split into a training set of 1000 and a test set of *n*_test_ = 307 accessions. Although (for convenience) we simulated data for all 1307 accessions before and after the intervention, the training data are observed prior to the intervention, and the test data afterwards. In other words, data for the test-set prior to the intervention and the training set after intervention are not used; the objective is to predict the target trait *ϒ*_3_ for the test accessions *after* the intervention, based on observations on the training set *prior* to the intervention.
- In the main set of simulations, the training set was further split into a set of 500 accessions used for inferring the GSEM with PCgen, and the remaining *n*_est_ = 500 accessions were used to estimate Λ, Σ_*G*_ and Σ_*E*_, and perform genomic prediction. We will refer to these sets as the PCgen and the estimation set, respectively. We made this distinction in order to avoid using the same data twice, which might lead to overfitting. Alternatively, we considered just a single training set of 500 and test set of 807 accessions; in this case we used the training set both for inferring the graph, parameter estimation and genomic prediction.
- Λ, Σ_*G*_ and Σ_*E*_ were estimated using the approach outlined in Appendix A.2, using estimates 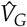 and 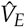 based on the estimation set, and using a topological ordering compatible with the PCgen reconstruction of the graph. 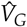 and 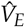 were obtained as in (Furlotte and Eskin 2015), i.e., by fitting a bivariate MTM for each pair of traits, using the replicates as well as the 100 SNPs. In case the resulting estimates 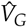 and 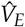 had negative eigenvalues we used the `nearPD` function to obtain the nearest positive-definite matrix. We recall that for the (unknown) true values of *V*_*G*_ and *V*_*E*_, the *LDL^t^* decomposition from Appendix A.2 gives exactly the matrices Λ, Σ_*G*_ and Σ_*E*_ used to simulate the data. However, given *estimates* 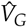 and 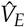, the resulting estimates 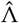 and 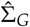 typically have small nonzero elements, which, according to the graph, should be zero (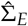 is always diagonal, by construction). We did not put these elements to zero, and used the original estimates 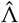 and 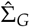.
- Let 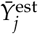 denote the accessions means of the *j*th trait on the estimation set (recall that, in contrast to PCgen, our genomic prediction is based on genotypic means, and uses the estimation set). Let *M*_est_ denote the *n*_est_ × *q* matrix of standardized SNP-scores for the *n*est accessions in the estimation set, and let *M*_test_ be the SNP-scores for the test set. When PCgen correctly reconstructs the edges *G* → *ϒ*_3_ ← *ϒ*_2_ and the absence of the edge *ϒ*_1_ → *ϒ*_3_, we perform genomic prediction with the SNP-BLUP, using the estimates 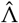, 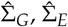 and the transformed phenotype (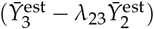), which captures the direct genetic effect on *ϒ*_3_:

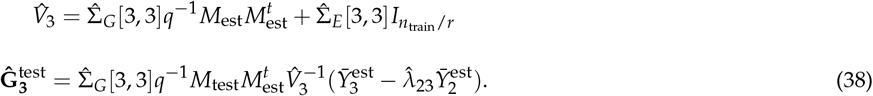 No genomic prediction was performed if the PCgen reconstruction contained undirected edges. When the PCgen reconstruction had no edge *G* → *ϒ*_3_, with *ϒ*_3_ being either isolated or having *ϒ*_2_ as only parent, the prediction was put to a constant.
- For comparison, we also performed genomic prediction with:

– The standard univariate GBLUP of the total genetic effects, which is the BLUP of the *q* = 100 SNP-effects (given 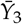 on the estimation set), multiplied by the SNP-scores for the test set (*M*_test_):

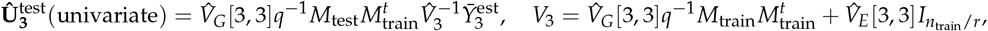

where we used all (1000) accessions from the training set (i.e., the PCgen set and the estimation set combined). Note that we use the notation 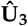 instead of 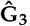, as the GBLUP predicts the *total* genetic effect, and not the direct effect.
– The conditional GBLUP for *ϒ*_3_, conditional on the observed *ϒ*_3_ and *ϒ*_2_ = 1:

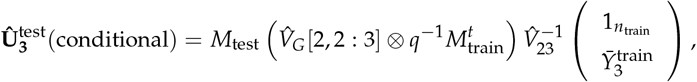

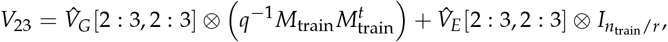

where 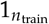 denotes a column vector of ones, 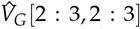 contains the second and third row and column of 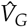, and 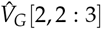, is the 1 × 2 vector with elements 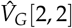 and 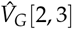. Again, we used the complete training set.

#### Results

First we discuss the results shown in Figure 12, obtained with separate parts of the training set used for PCgen and parameter estimation, and a test set of 307 accessions.

- In 182 of the 200 simulated datasets, PCgen identified the correct GSEM, and in 12 cases the reconstruction contained undirected edges. For two simulated datasets, the PCgen output was *G* → *ϒ*_1_ → *ϒ*_2_ → *ϒ*_3_, and the prediction for *ϒ*_3_ was put to a constant (3). In four datasets, one or both of the edges *G* → *ϒ*_1_ and *ϒ*_1_ → *ϒ*_2_ were not detected, but the relevant part was reconstructed correctly (*G* → *ϒ*_3_ → *ϒ*_2_, and no edge *ϒ*_1_ → *ϒ*_3_). Consequently, equation (38) could be used for predicting the test set after intervention, for 186 (93%) of the simulated datasets. Figure 12 compares the accuracy for these datasets, compared to the accuracy achieved with the conditional GBLUP (also after intervention) and the univariate GBLUP (before and after intervention).
- Looking at Figure 11, the prediction of **Y_3_** after intervention requires accurate prediction of the direct effect **G_3_**. This is exactly what is achieved with equation (38), where we obtain parameter estimates based on the ‘adjusted’ trait 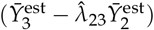 prior to intervention, i.e., controlling for the indirect path *G* → *ϒ*_1_ →*ϒ*_*2*_ → *ϒ*_3_. Consequently, the accuracy is high, as the ‘direct’ heritability of *ϒ*_3_ is close to 0.9 (Σ*G* [3, 3] = 4 and Σ_*E*_ [3, 3] = 1; the latter being twice as small for the accession means).
- We achieve lower accuracy when we use the univariate GBLUP to predict **Y3** after the intervention (Figure 12, middle), as we target the wrong quantity: instead of the direct effect **G_3_**, the GBLUP predicts the *total* genetic effect **U_3_**.
- For predicting **Y_3_***before* the intervention (Figure 12, right), **U_3_** is indeed the right quantity to predict, but accuracy is still lower. This is because the (line)heritability of **Y_3_** is only around 0.5 (recall from equation (37) that *V*_*G*_ [3, 3] = 17.5 and, for the accession means, the residual variance is 32.5/2).
- Also the conditional GBLUP (Figure 12, left) has suboptimal performance, the predictions being very similar to those obtained with univariate GBLUP (both have average accuracy of around 0.42). The conditioning on **Y_2_** = 1 might intuitively make sense, but has little impact and effectively shrinks the marker effects to the wrong point.

Finally, we simulated a new series of 200 datasets before and after intervention, but now defined training sets of 500 and test sets of 807, and used the training set both for causal inference and for parameter estimation. Also the univariate and conditional GBLUP used these training sets. The results (Figure 13) are very similar, suggesting that there is little overfitting. Again, PCgen inferred the correct graph for around 90% of the datasets.

**Figure 12.**
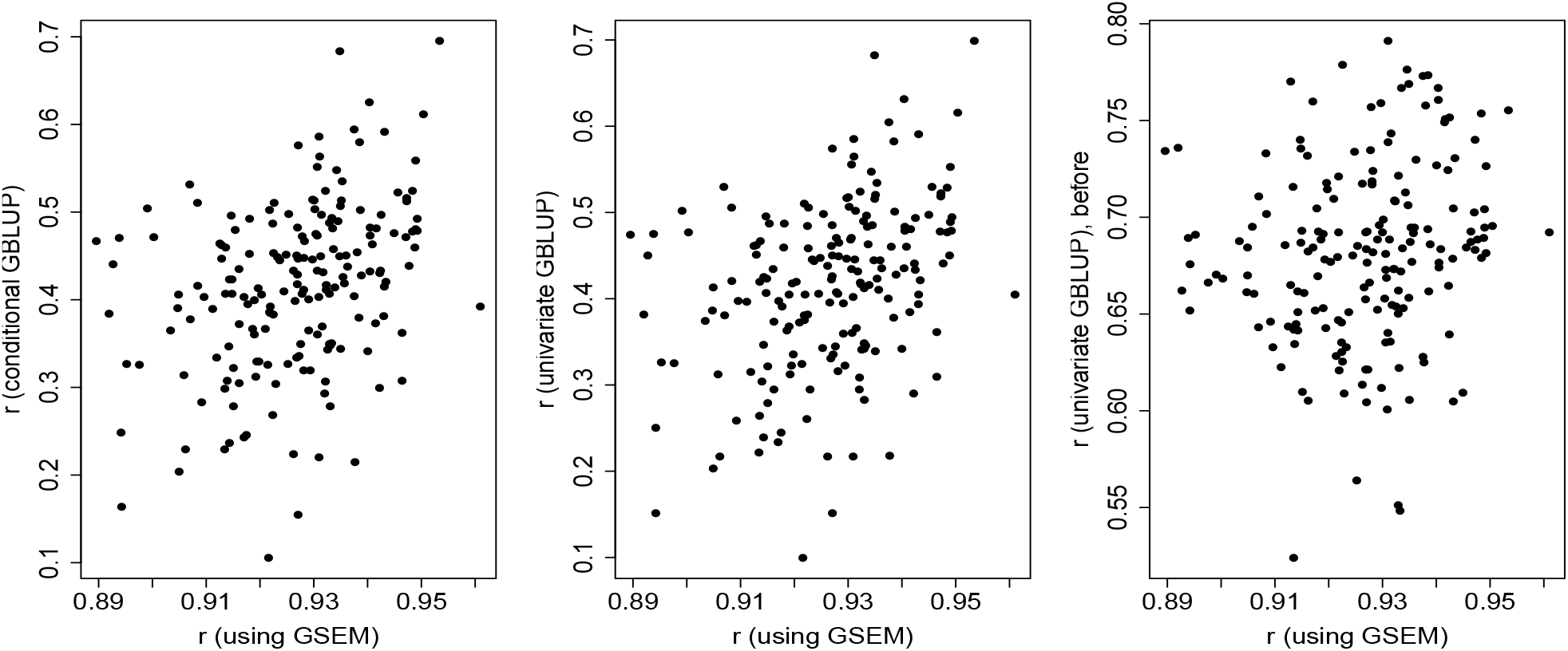
Accuracy (Pearson *r*) of the prediction of **Y_3_**, using PCgen and equation (38) (horizontal axis), and using the conditional or univariate GBLUP (vertical axis). Each training set contained 1000 accessions, observed prior to the intervention **Y_2_**:= 1; 500 accessions were used for inferring the network, and 500 for parameter estimation and genomic prediction for 307 accessions in the test set. The conditional and univariate GBLUP used all 1000 accessions for genomic prediction. Except for the GBLUP in the right panel, accuracy is evaluated using the correlation with **Y_3_** observed on the test set *after* the intervention. Each plot contains 186 points, corresponding to the datasets for which PCgen correctly reconstructed either the complete network, or at least the edges *G → ϒ_3_ ← ϒ_2_* and the absence of *ϒ_1_ → ϒ_3_*. In 12 of the remaining 14 cases, PCgen output contained undirected edges, and we could not define a predictor of **G3**. In two cases, **G3** was put to a constant, as the inferred network was wrong.

**Figure 13.**
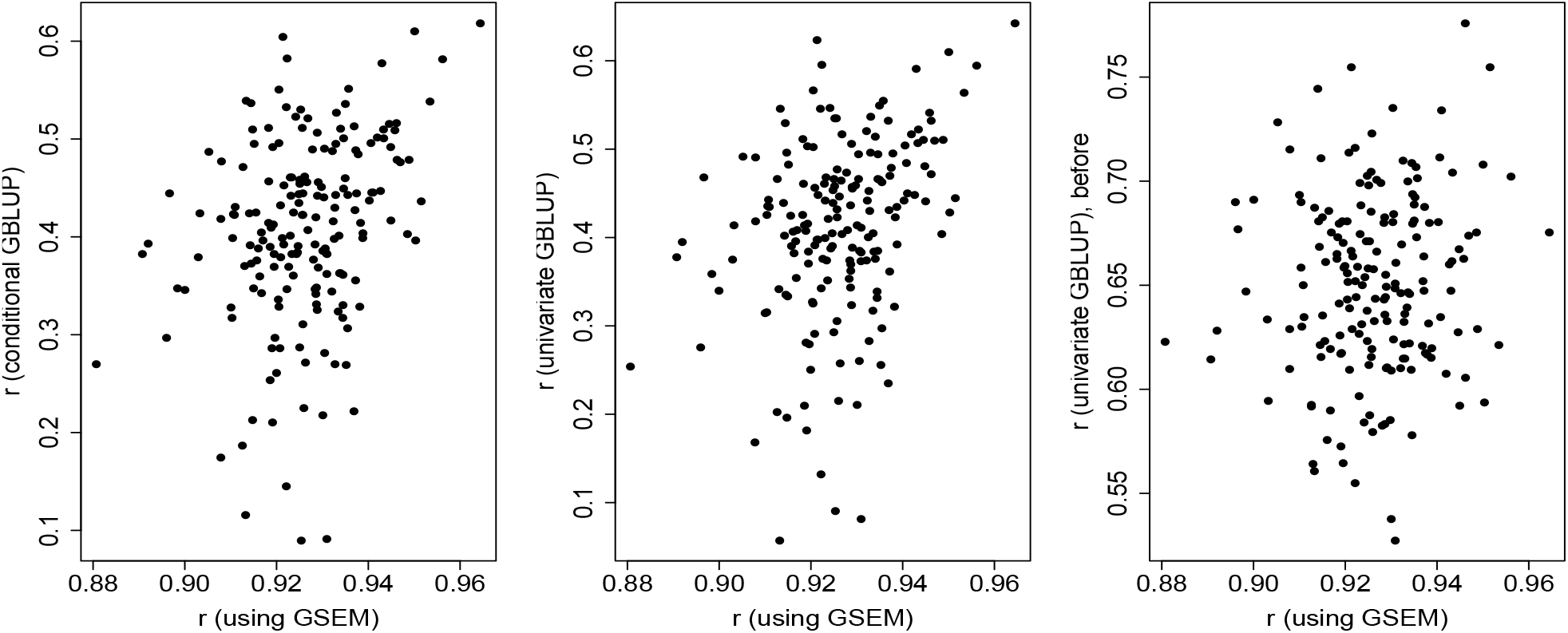
Accuracy (Pearson *r*) of the prediction of **Y_3_**, using PCgen and equation (38) (horizontal axis), and using the conditional or univariate GBLUP (vertical axis). Each training set contained 500 accessions, observed prior to the intervention **Y_2_**:= 1. Except for the GBLUP in the right panel, accuracy is evaluated using the correlation with **Y_3_** observed on the test set *after* the intervention. Each plot contains 181 points, corresponding to the datasets for which PCgen correctly reconstructed either the complete network, or at least the edges *G → ϒ_3_ ← ϒ_2_* and the absence of *ϒ_1_ → ϒ_3_*. In 14 of the remaining 19 cases, PCgen output contained undirected edges, and we could not define a predictor of **G3**. In 5 cases, the inferred network was wrong.

**Table S1.**
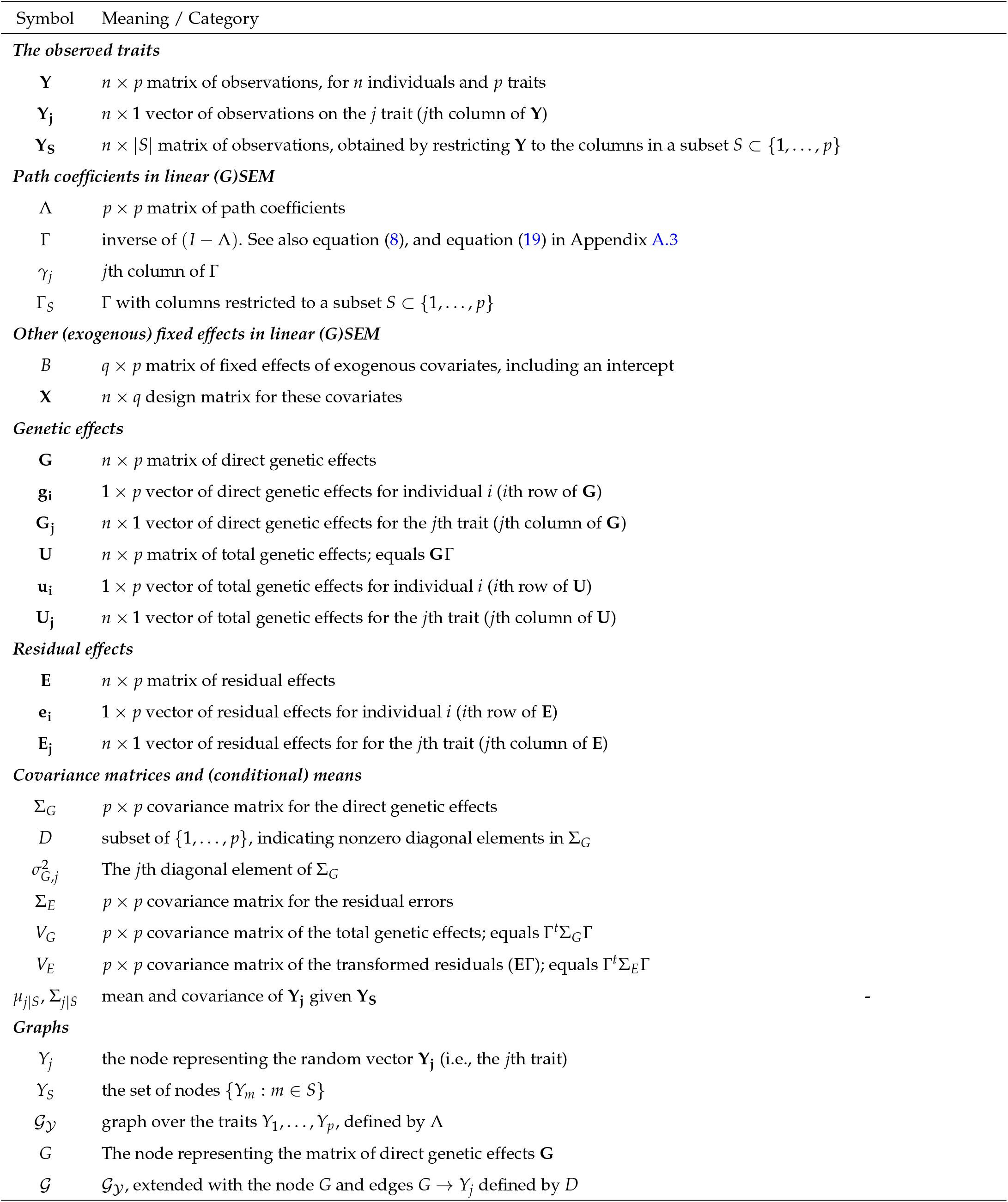
Overview of notation. Appendix A.1 provides additional graph-theoretic terminology.

**Table S2.** Performance of PCgen and residuals-based approaches, averaged over 500 simulated datasets, for scenarios 1 and 2. For ease of presentation we used in the main text the abbreviations ‘PCgen’, ‘PCres(means)’ and ‘PCres(replicates)’, to refer to respectively PCgen-screening, PCres-multi-A and PCres-uni-R. See also Tables 2 and 3 in File S2.

| | $\mathcal{G}_Y$ | | | | $G \rightarrow Y_j$ | | |
| --- | --- | --- | --- | --- | --- | --- | --- |
|  | TPR | FPR | TDR | SHD | TPR | FPR | TDR |
| Scenario 1 ($p = 4$) | $(p_t = 1/3)$ | | | | $(p_g = 0.5)$ | | |
| PCgen-RC | 0.652 | 0.009 | 0.964 | 0.454 | 0.982 | 0.026 | 0.995 |
| PCgen-RG-uni | 0.650 | 0.039 | 0.908 | 0.584 | 0.981 | 0.027 | 0.995 |
| PCgen-RC-screening | 0.647 | 0.006 | 0.981 | 0.442 | 0.982 | 0.026 | 0.995 |
| PCres-multi-R | 0.657 | 0.116 | 0.719 | 1.908 |  |  |  |
| PCres-uni-R | 0.650 | 0.039 | 0.908 | 1.410 |  |  |  |
| PCres-multi-RA | 0.658 | 0.088 | 0.779 | 1.728 |  |  |  |
| PCres-uni-RA | 0.653 | 0.026 | 0.932 | 1.374 |  |  |  |
| PCres-multi-A | 0.521 | 0.390 | 0.438 | 3.174 |  |  |  |
| PCres-uni-A | 0.640 | 0.083 | 0.758 | 1.760 |  |  |  |
| PC-GBLUP | 0.635 | 0.661 | 0.384 | 4.284 |  |  |  |
| Scenario 2 ($p = 4$) | $(p_t = 0.5)$ | | | | $(p_g = 0.5)$ | | |
| PCgen-RC | 0.818 | 0.035 | 0.980 | 1.238 | 0.976 | 0.031 | 0.995 |
| PCgen-RG-uni | 0.819 | 0.073 | 0.939 | 1.358 | 0.974 | 0.032 | 0.996 |
| PCgen-RC-screening | 0.804 | 0.033 | 0.986 | 1.246 | 0.976 | 0.031 | 0.995 |
| PCres-multi-R | 0.834 | 0.132 | 0.862 | 2.560 |  |  |  |
| PCres-uni-R | 0.819 | 0.073 | 0.939 | 2.320 |  |  |  |
| PCres-multi-RA | 0.831 | 0.098 | 0.884 | 2.430 |  |  |  |
| PCres-uni-RA | 0.821 | 0.058 | 0.947 | 2.266 |  |  |  |
| PCres-multi-A | 0.672 | 0.364 | 0.659 | 3.628 |  |  |  |
| PCres-uni-A | 0.782 | 0.124 | 0.863 | 2.688 |  |  |  |
| PC-GBLUP | 0.793 | 0.709 | 0.563 | 4.634 |  |  |  |

**Table S3.**
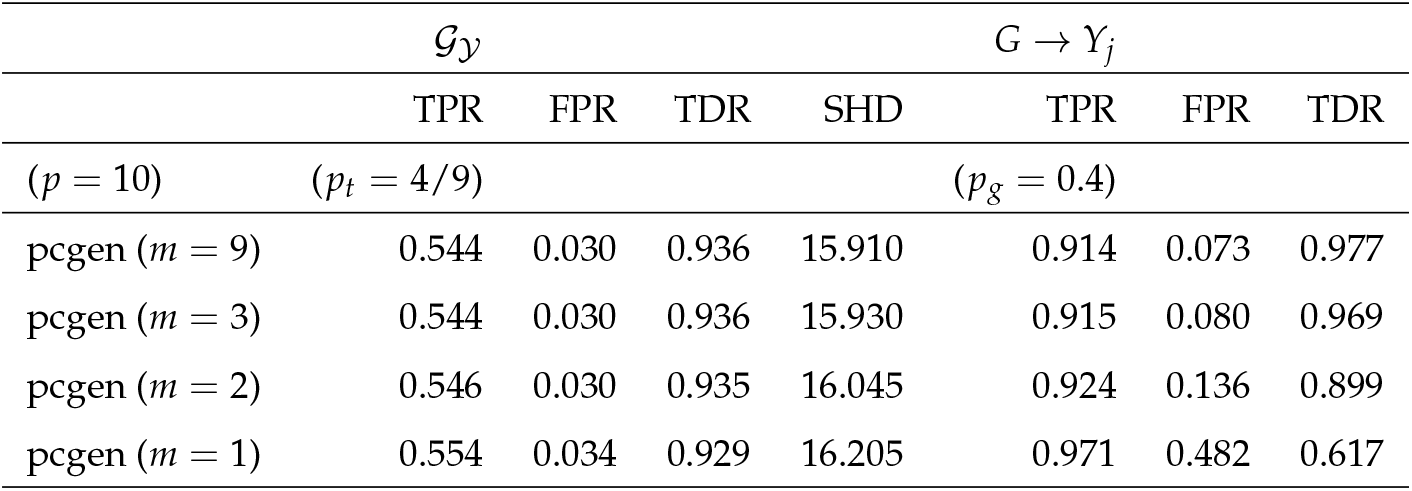
The effect of limiting the maximum size of the conditioning sets: performance of PCgen, averaged over 200 simulated datasets with *p* = 10 traits. We considered maximum sizes (*m*) of 1, 2, 3 and 9 (i.e., no limit).

| | $\mathcal{G}_Y$ | | | | $G \rightarrow Y_j$ | | |
| --- | --- | --- | --- | --- | --- | --- | --- |
|  | TPR | FPR | TDR | SHD | TPR | FPR | TDR |
| $(p = 10)$ | $(p_t = 4/9)$ | | | | $(p_g = 0.4)$ | | |
| pcgen ($m = 9$) | 0.544 | 0.030 | 0.936 | 15.910 | 0.914 | 0.073 | 0.977 |
| pcgen ($m = 3$) | 0.544 | 0.030 | 0.936 | 15.930 | 0.915 | 0.080 | 0.969 |
| pcgen ($m = 2$) | 0.546 | 0.030 | 0.935 | 16.045 | 0.924 | 0.136 | 0.899 |
| pcgen ($m = 1$) | 0.554 | 0.034 | 0.929 | 16.205 | 0.971 | 0.482 | 0.617 |

**Table S4.** Performance of PCgen on the graph *G → ϒ_1_ → ϒ_2_*, averaged over 200 simulated datasets. The path-coefficient associated with *ϒ_1_ → ϒ_2_* was 0.25, and Σ_*G*_ and Σ_*E*_ were diagonal, with elements (1, 0) and (3, 3), respectively. For each trait *j* = 1, 2 we added independent Gaussian measurement errors with variance *αm VE* [*j*, *j*], where *αm* is a constant and *VE* [*j*, *j*] the *j*th element on the diagonal of *VE*, containing the total residual variance of trait *j*, before adding measurement error (see e.g., equation (7) in the main text). Since the graph 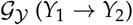 does not contain gaps, the FPR is always defined to be 1.

| | $\mathcal{G}_Y$ | | | | $G \rightarrow Y_j$ | | |
| --- | --- | --- | --- | --- | --- | --- | --- |
|  | TPR | FPR | TDR | SHD | TPR | FPR | TDR |
| pcgen ($r = 2, n = 250, \alpha_m = 0$) | 0.905 | 1.000 | 1.000 | 1.000 | 0.890 | 0.115 | 0.888 |
| pcgen ($r = 2, n = 250, \alpha_m = 0.2$) | 0.740 | 1.000 | 1.000 | 0.995 | 0.785 | 0.215 | 0.785 |
| pcgen ($r = 2, n = 250, \alpha_m = 1$) | 0.535 | 1.000 | 1.000 | 1.000 | 0.400 | 0.615 | 0.398 |
| pcgen ($r = 4, n = 250, \alpha_m = 1$) | 0.710 | 1.000 | 1.000 | 0.995 | 0.980 | 0.025 | 0.978 |
| pcgen ($r = 2, n = 500, \alpha_m = 1$) | 0.780 | 1.000 | 1.000 | 0.995 | 0.740 | 0.265 | 0.738 |

**Table S5.** Performance of PCgen on the graph *ϒ_1_ → ϒ_2_* with direct genetic effects on *ϒ_1_* and *ϒ_2_* (*ϒ*_1_ ← *G* → *Y*_2_), averaged over three series of 200 simulated datasets. The path-coefficient associated with *ϒ*_1_ → *ϒ*_2_ was 0.25, and Σ_*E*_ was diagonal with elements (3, 3). Σ_*G*_ had diagonal (1, 1), the off-diagonal element being respectively 0, 0.5 and 0.95 in the different datasets.

| | $\mathcal{G}_Y$ | | | | $G \rightarrow Y_j$ | | |
| --- | --- | --- | --- | --- | --- | --- | --- |
|  | TPR | FPR | TDR | SHD | TPR | FPR | TDR |
| pcgen ($\Sigma_G[1, 2] = 0$) | 0.960 | 1.000 | 1.000 | 0.950 | 0.940 | 0.000 | 0.995 |
| pcgen ($\Sigma_G[1, 2] = 0.5$) | 0.980 | 1.000 | 1.000 | 0.920 | 0.852 | 0.000 | 0.980 |
| pcgen ($\Sigma_G[1, 2] = 0.95$) | 0.975 | 1.000 | 1.000 | 0.935 | 0.595 | 0.000 | 0.910 |

**Table S6.**
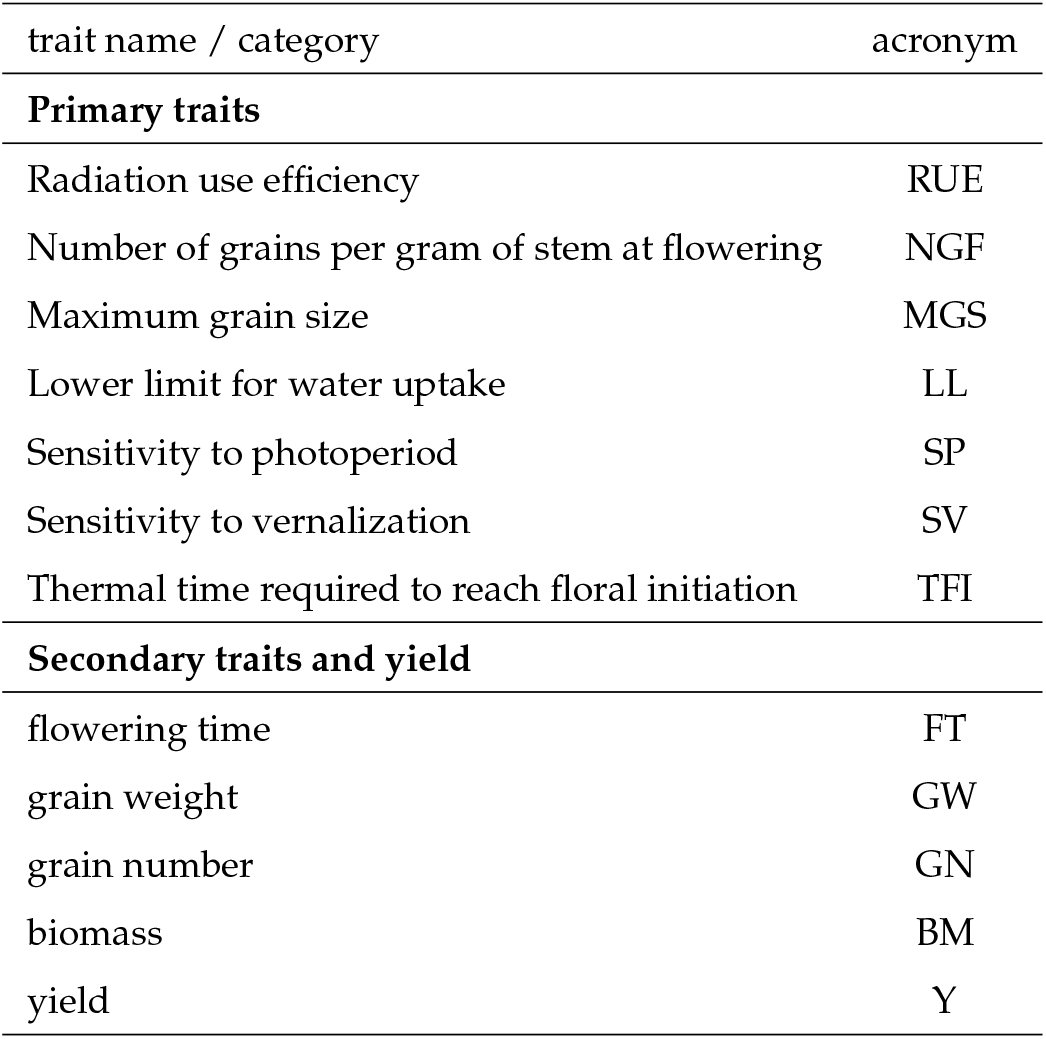
Acronyms for 12 traits simulated using APSIM.

| trait name / category | acronym |
| --- | --- |
| **Primary traits** |  |
| Radiation use efficiency | RUE |
| Number of grains per gram of stem at flowering | NGF |
| Maximum grain size | MGS |
| Lower limit for water uptake | LL |
| Sensitivity to photoperiod | SP |
| Sensitivity to vernalization | SV |
| Thermal time required to reach floral initiation | TFI |
| **Secondary traits and yield** |  |
| flowering time | FT |
| grain weight | GW |
| grain number | GN |
| biomass | BM |
| yield | Y |

**Table S7.** Maize data: trait acronyms

| trait name / category | acronym |
| --- | --- |
| **Flowering** |  |
| anthesis | A |
| silking | Sk |
| **Height** |  |
| ear height | EH |
| plant height | PH |
| tassel height | TH |
| **Yield related** |  |
| grain weight | GW |
| yield | Y |

**Table S8.**
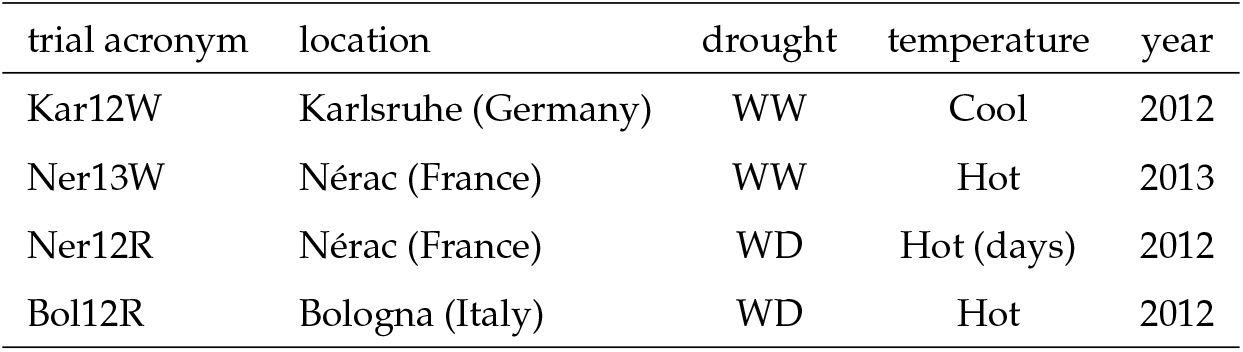
Maize data: overview of the trials. Details on the environmental classification are given in the main text, and in *Millet et al.* (2016).

| trial acronym | location | drought | temperature | year |
| --- | --- | --- | --- | --- |
| Kar12W | Karlsruhe (Germany) | WW | Cool | 2012 |
| Ner13W | Nérac (France) | WW | Hot | 2013 |
| Ner12R | Nérac (France) | WD | Hot (days) | 2012 |
| Bol12R | Bologna (Italy) | WD | Hot | 2012 |

**Table S9.** PCgen-reconstruction for the 4 maize trials, with *α* = 0.01: p-values. For each pair of nodes, the table contains the highest p-value observed across all conditional independence tests performed for that pair.

| **Kar12W** | G | PH | TH | EH | A | Sk | Y | GW |
| --- | --- | --- | --- | --- | --- | --- | --- | --- |
| G | 1.00E+00 | 1.75E-08 | 1.62E-04 | 4.06E-10 | 4.83E-09 | 3.98E-18 | 3.76E-14 | 3.44E-26 |
| PH | 1.75E-08 | 1.00E+00 | 0.00E+00 | 2.17E-04 | 6.27E-02 | 3.06E-01 | 1.81E-01 | 3.66E-01 |
| TH | 1.62E-04 | 0.00E+00 | 1.00E+00 | 3.60E-01 | 3.10E-02 | 1.08E-02 | 1.65E-01 | 1.00E-01 |
| EH | 4.06E-10 | 2.17E-04 | 3.60E-01 | 1.00E+00 | 1.25E-01 | 2.07E-01 | 2.58E-01 | 1.09E-01 |
| A | 4.83E-09 | 6.27E-02 | 3.10E-02 | 1.25E-01 | 1.00E+00 | 0.00E+00 | 3.01E-01 | 9.68E-01 |
| Sk | 3.98E-18 | 3.06E-01 | 1.08E-02 | 2.07E-01 | 0.00E+00 | 1.00E+00 | 5.26E-01 | 1.08E-02 |
| Y | 3.76E-14 | 1.81E-01 | 1.65E-01 | 2.58E-01 | 3.01E-01 | 5.26E-01 | 1.00E+00 | 2.22E-01 |
| GW | 3.44E-26 | 3.66E-01 | 1.00E-01 | 1.09E-01 | 9.68E-01 | 1.08E-02 | 2.22E-01 | 1.00E+00 |
| **Ner13W** | G | PH | TH | EH | A | Sk | Y | GW |
| G | 1.00E+00 | 1.79E-07 | 4.49E-10 | 8.16E-13 | 5.80E-15 | 1.21E-34 | 3.81E-34 | 2.83E-41 |
| PH | 1.79E-07 | 1.00E+00 | 0.00E+00 | 3.71E-03 | 5.53E-01 | 4.79E-02 | 7.80E-01 | 4.06E-01 |
| TH | 4.49E-10 | 0.00E+00 | 1.00E+00 | 1.41E-02 | 5.74E-01 | 4.21E-02 | 4.05E-03 | 8.43E-01 |
| EH | 8.16E-13 | 3.71E-03 | 1.41E-02 | 1.00E+00 | 3.70E-01 | 1.17E-02 | 2.65E-02 | 5.91E-01 |
| A | 5.80E-15 | 5.53E-01 | 5.74E-01 | 3.70E-01 | 1.00E+00 | 1.60E-12 | 3.20E-01 | 1.94E-02 |
| Sk | 1.21E-34 | 4.79E-02 | 4.21E-02 | 1.17E-02 | 1.60E-12 | 1.00E+00 | 3.59E-01 | 1.87E-01 |
| Y | 3.81E-34 | 7.80E-01 | 4.05E-03 | 2.65E-02 | 3.20E-01 | 3.59E-01 | 1.00E+00 | 3.62E-02 |
| GW | 2.83E-41 | 4.06E-01 | 8.43E-01 | 5.91E-01 | 1.94E-02 | 1.87E-01 | 3.62E-02 | 1.00E+00 |
| **Ner12R** | G | PH |  | EH | A | Sk | Y | GW |
| G | 1.00E+00 | 8.68E-14 |  | 1.47E-25 | 1.39E-68 | 1.16E-50 | 5.26E-62 | 7.03E-59 |
| PH | 8.68E-14 | 1.00E+00 |  | 4.65E-13 | 1.70E-02 | 7.46E-01 | 8.86E-03 | 5.00E-01 |
| EH | 1.47E-25 | 4.65E-13 |  | 1.00E+00 | 4.53E-01 | 3.52E-01 | 1.60E-01 | 1.66E-02 |
| A | 1.39E-68 | 1.70E-02 |  | 4.53E-01 | 1.00E+00 | 6.56E-03 | 2.25E-01 | 3.43E-01 |
| Sk | 1.16E-50 | 7.46E-01 |  | 3.52E-01 | 6.56E-03 | 1.00E+00 | 3.55E-09 | 2.59E-02 |
| Y | 5.26E-62 | 8.86E-03 |  | 1.60E-01 | 2.25E-01 | 3.55E-09 | 1.00E+00 | 1.79E-02 |
| GW | 7.03E-59 | 5.00E-01 |  | 1.66E-02 | 3.43E-01 | 2.59E-02 | 1.79E-02 | 1.00E+00 |
| **Bol12R** | G | PH | TH | EH | A | Sk | Y | GW |
| G | 1.00E+00 | 8.58E-11 | 2.92E-11 | 1.80E-51 | 4.40E-25 | 5.92E-37 | 4.55E-32 | 3.17E-61 |
| PH | 8.58E-11 | 1.00E+00 | 0.00E+00 | 5.09E-07 | 7.89E-02 | 3.25E-03 | 1.32E-01 | 9.86E-01 |
| TH | 2.92E-11 | 0.00E+00 | 1.00E+00 | 1.52E-05 | 1.23E-01 | 3.36E-01 | 1.53E-02 | 9.71E-01 |
| EH | 1.80E-51 | 5.09E-07 | 1.52E-05 | 1.00E+00 | 1.78E-01 | 2.29E-01 | 7.36E-02 | 2.71E-01 |
| A | 4.40E-25 | 7.89E-02 | 1.23E-01 | 1.78E-01 | 1.00E+00 | 1.47E-10 | 2.54E-03 | 9.30E-02 |
| Sk | 5.92E-37 | 3.25E-03 | 3.36E-01 | 2.29E-01 | 1.47E-10 | 1.00E+00 | 3.99E-09 | 2.99E-02 |
| Y | 4.55E-32 | 1.32E-01 | 1.53E-02 | 7.36E-02 | 2.54E-03 | 3.99E-09 | 1.00E+00 | 2.65E-04 |
| GW | 3.17E-61 | 9.86E-01 | 9.71E-01 | 2.71E-01 | 9.30E-02 | 2.99E-02 | 2.65E-04 | 1.00E+00 |

**Table S10.** Estimates of total genetic correlation in the four maize trials, with significance between parentheses.

| **Kar12W** | PH | TH | EH | A | Sk | Y | GW |
| --- | --- | --- | --- | --- | --- | --- | --- |
| PH | NA (NA) | 0.91 (0) | 0.59 (1.1e-11) | 0.53 (2.6e-12) | 0.47 (2.1e-10) | 0.32 (0.00035) | -0.099 (0.23) |
| TH | 0.91 (0) | NA (NA) | 0.61 (4.9e-11) | 0.49 (4.7e-10) | 0.48 (7.5e-10) | 0.39 (4.2e-05) | -0.15 (0.074) |
| EH | 0.59 (1.1e-11) | 0.61 (4.9e-11) | NA (NA) | 0.56 (3.2e-12) | 0.51 (1.2e-10) | 0.16 (0.11) | -0.19 (0.029) |
| A | 0.53 (2.6e-12) | 0.49 (4.7e-10) | 0.56 (3.2e-12) | NA (NA) | 0.96 (0) | 0.31 (0.00019) | -0.4 (6.2e-08) |
| Sk | 0.47 (2.1e-10) | 0.48 (7.5e-10) | 0.51 (1.2e-10) | 0.96 (0) | NA (NA) | 0.28 (0.00076) | -0.44 (1.3e-09) |
| Y | 0.32 (0.00035) | 0.39 (4.2e-05) | 0.16 (0.11) | 0.31 (0.00019) | 0.28 (0.00076) | NA (NA) | 0.13 (0.15) |
| GW | -0.099 (0.23) | -0.15 (0.074) | -0.19 (0.029) | -0.4 (6.2e-08) | -0.44 (1.3e-09) | 0.13 (0.15) | NA (NA) |
| **Ner13W** | PH | TH | EH | A | Sk | Y | GW |
| PH | NA (NA) | 0.92 (0) | 0.7 (1.1e-16) | 0.59 (1.3e-15) | 0.49 (5.9e-11) | 0.48 (1.3e-09) | 0.0095 (0.9) |
| TH | 0.92 (0) | NA (NA) | 0.67 (0) | 0.66 (0) | 0.57 (5.6e-16) | 0.51 (8.4e-12) | 0.02 (0.8) |
| EH | 0.7 (1.1e-16) | 0.67 (0) | NA (NA) | 0.66 (0) | 0.58 (1.4e-15) | 0.31 (0.00012) | -0.042 (0.61) |
| A | 0.59 (1.3e-15) | 0.66 (0) | 0.66 (0) | NA (NA) | 0.94 (0) | 0.52 (1.9e-14) | -0.11 (0.11) |
| Sk | 0.49 (5.9e-11) | 0.57 (5.6e-16) | 0.58 (1.4e-15) | 0.94 (0) | NA (NA) | 0.37 (1.1e-07) | -0.24 (0.00062) |
| Y | 0.48 (1.3e-09) | 0.51 (8.4e-12) | 0.31 (0.00012) | 0.52 (1.9e-14) | 0.37 (1.1e-07) | NA (NA) | 0.15 (0.043) |
| GW | 0.0095 (0.9) | 0.02 (0.8) | -0.042 (0.61) | -0.11 (0.11) | -0.24 (0.00062) | 0.15 (0.043) | NA (NA) |
| **Ner12R** | PH |  | EH | A | Sk | Y | GW |
| PH | NA (NA) |  | 0.29 (0.0034) | 0.039 (0.65) | -0.09 (0.31) | 0.26 (0.0037) | 0.16 (0.083) |
| EH | 0.29 (0.0034) |  | NA (NA) | 0.57 (2.7e-15) | 0.21 (0.0058) | 0.21 (0.0055) | -0.21 (0.0088) |
| A | 0.039 (0.65) |  | 0.57 (2.7e-15) | NA (NA) | 0.68 (0) | 0.12 (0.081) | -0.16 (0.026) |
| Sk | -0.09 (0.31) |  | 0.21 (0.0058) | 0.68 (0) | NA (NA) | -0.44 (2.2e-11) | -0.23 (0.00097) |
| Y | 0.26 (0.0037) |  | 0.21 (0.0055) | 0.12 (0.081) | -0.44 (2.2e-11) | NA (NA) | 0.38 (7.9e-08) |
| GW | 0.16 (0.083) |  | -0.21 (0.0088) | -0.16 (0.026) | -0.23 (0.00097) | 0.38 (7.9e-08) | NA (NA) |
| **Bol12R** | PH | TH | EH | A | Sk | Y | GW |
| PH | NA (NA) | 0.89 (0) | 0.66 (0) | 0.42 (3.4e-08) | 0.38 (5.9e-07) | -0.046 (0.58) | 0.02 (0.8) |
| TH | 0.89 (0) | NA (NA) | 0.53 (1.4e-12) | 0.37 (1.7e-06) | 0.34 (9.2e-06) | -0.051 (0.54) | 0.12 (0.13) |
| EH | 0.66 (0) | 0.53 (1.4e-12) | NA (NA) | 0.44 (1.9e-10) | 0.36 (4.8e-07) | -0.025 (0.74) | -0.12 (0.11) |
| A | 0.42 (3.4e-08) | 0.37 (1.7e-06) | 0.44 (1.9e-10) | NA (NA) | 0.9 (0) | -0.43 (3.3e-09) | -0.11 (0.15) |
| Sk | 0.38 (5.9e-07) | 0.34 (9.2e-06) | 0.36 (4.8e-07) | 0.9 (0) | NA (NA) | -0.61 (0) | -0.1 (0.16) |
| Y | -0.046 (0.58) | -0.051 (0.54) | -0.025 (0.74) | -0.43 (3.3e-09) | -0.61 (0) | NA (NA) | 0.0068 (0.93) |
| GW | 0.02 (0.8) | 0.12 (0.13) | -0.12 (0.11) | -0.11 (0.15) | -0.1 (0.16) | 0.0068 (0.93) | NA (NA) |

**Table S11.** Rice data: trait acronyms, following the description of Kadam *et al.* (2017), with derived traits removed.

| Traits | Trait | acronym | Unit |
| --- | --- | --- | --- |
| (A) Shoot morphological traits |  |  |  |
|  | Plant height | PHT | cm |
| | Tiller number | TN | $plant^{-1}$ |
| | Total leaf area | TLA | $m^2 plant^{-1}$ |
| (B) Physiological traits |  |  |  |
| | Cumulative water transpiration | CWT | $kg plant^{-1}$ |
| (C) Root anatomical traits |  |  |  |
|  | Maximum root length | MRL | cm |
| | Surface area | SA | $cm^2 plant^{-1}$ |
| | Root volume | RV | $cm^3 plant^{-1}$ |
|  | Average root thickness | ART | mm |
| | Total root weight density | TRWD | $g cm^{-3}$ |
|  | Root length (RL) with diameter class |  |  |
| | RL_0-0.5 | RL005 | $m plant^{-1}$ |
| | RL_0.5-1.0 | RL0510 | $m plant^{-1}$ |
| | RL_1.0-1.5 | RL1015 | $m plant^{-1}$ |
| | RL_1.5-2.0 | RL1520 | $m plant^{-1}$ |
| | RL_2.0-2.5 | RL2025 | $m plant^{-1}$ |
| | RL_2.5-3.0 | RL2530 | $m plant^{-1}$ |
| | RL_3.0-3.5 | RL3035 | $m plant^{-1}$ |
| | RL_3.5 | RL35 | $m plant^{-1}$ |
| (D) Root anatomical traits |  |  |  |
| | Root diameter | RD | $\mu m$ |
| | Cortex diameter | CD | $\mu m$ |
| | Stele diameter | SD | $\mu m$ |
| | Late metaxylem diameter | LMXD | $\mu m$ |
| | Late metaxylem number | LMXN | $\mu m$ |
| (E) Dry matter traits |  |  |  |
| | Leaf weight | LW | $g plant^{-1}$ |
| | Stem weight | SW | $g plant^{-1}$ |
| | Root weight | RW | $g plant^{-1}$ |

**Figure S1.**
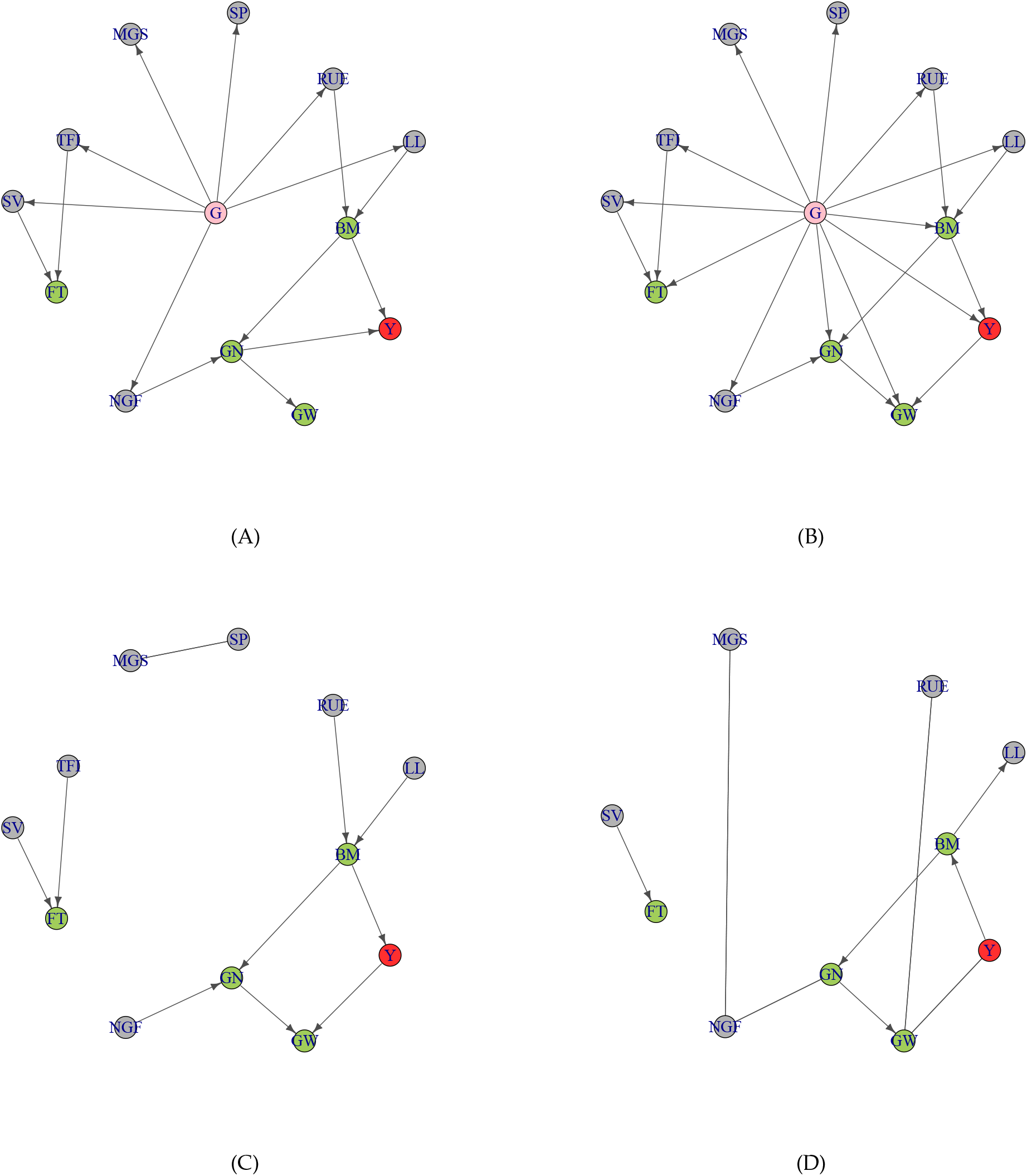
Networks of 12 APSIM traits in the dry environment Emerald 1993 (severe drought), including seven primary traits (grey), four secondary traits (green) and yield (red). (A) Summary graph, (B) PCgen, (C) pcRes and (D) PC-stable, using the 300 simulated QTLs. Trait acronoyms are given in Table S6.

**Figure S2.**
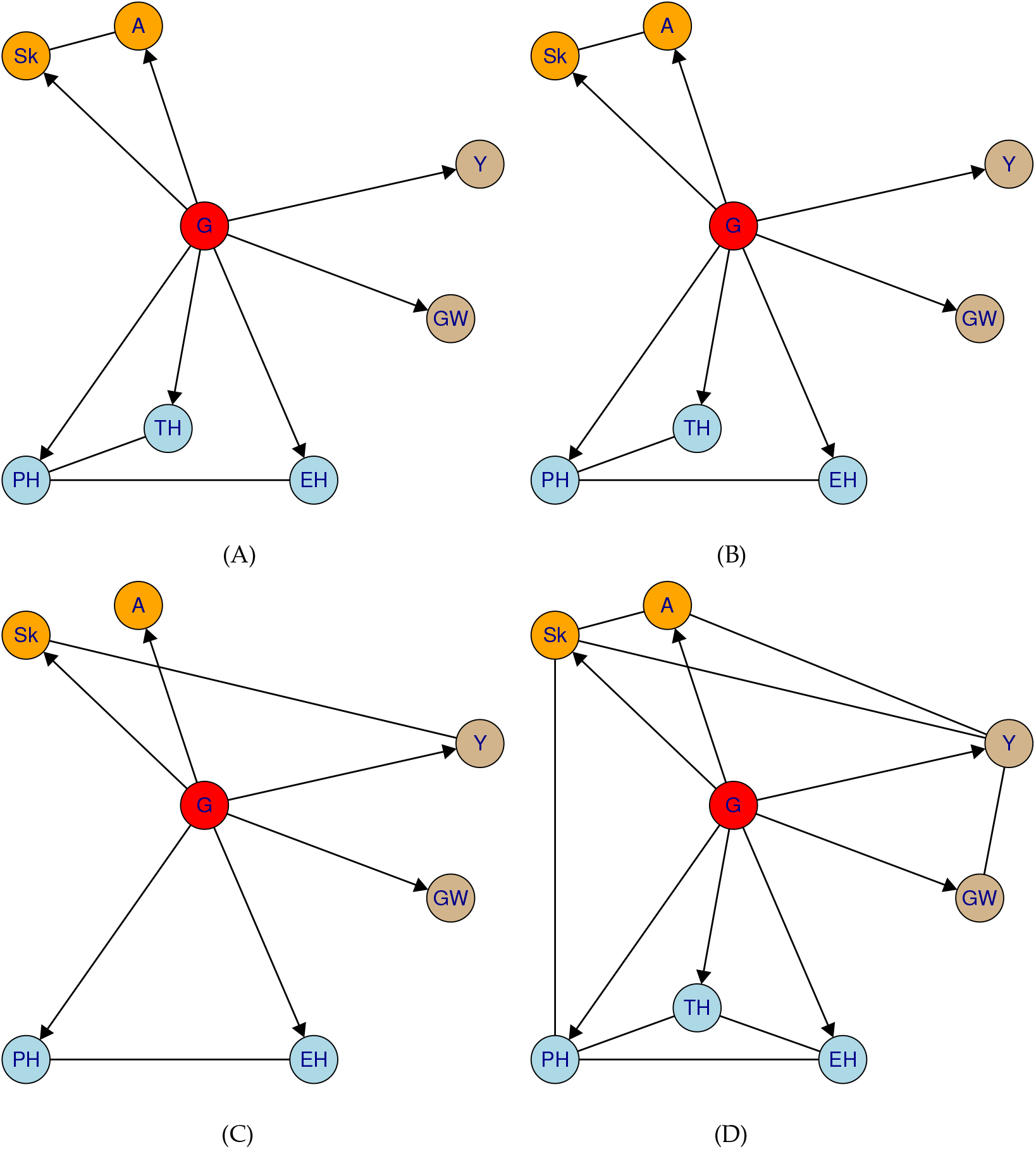
Estimated networks with *α* = 0.001, for four of the DROPS field trials: (A) Kar12W, (WW, Cool), (B) Ner13W (WW, Hot), (C) Ner12R (WD, Hot (days)) and (D) Bol12R (WD, Hot). Trait categories are flowering (orange), height (blue) and yield (brown). Each trial represents a different environmental scenario, being well-watered (WW) or water-deficit (WD), and different temperatures (see text). Trait acronyms are explained in Table S7.

**Figure S3.**
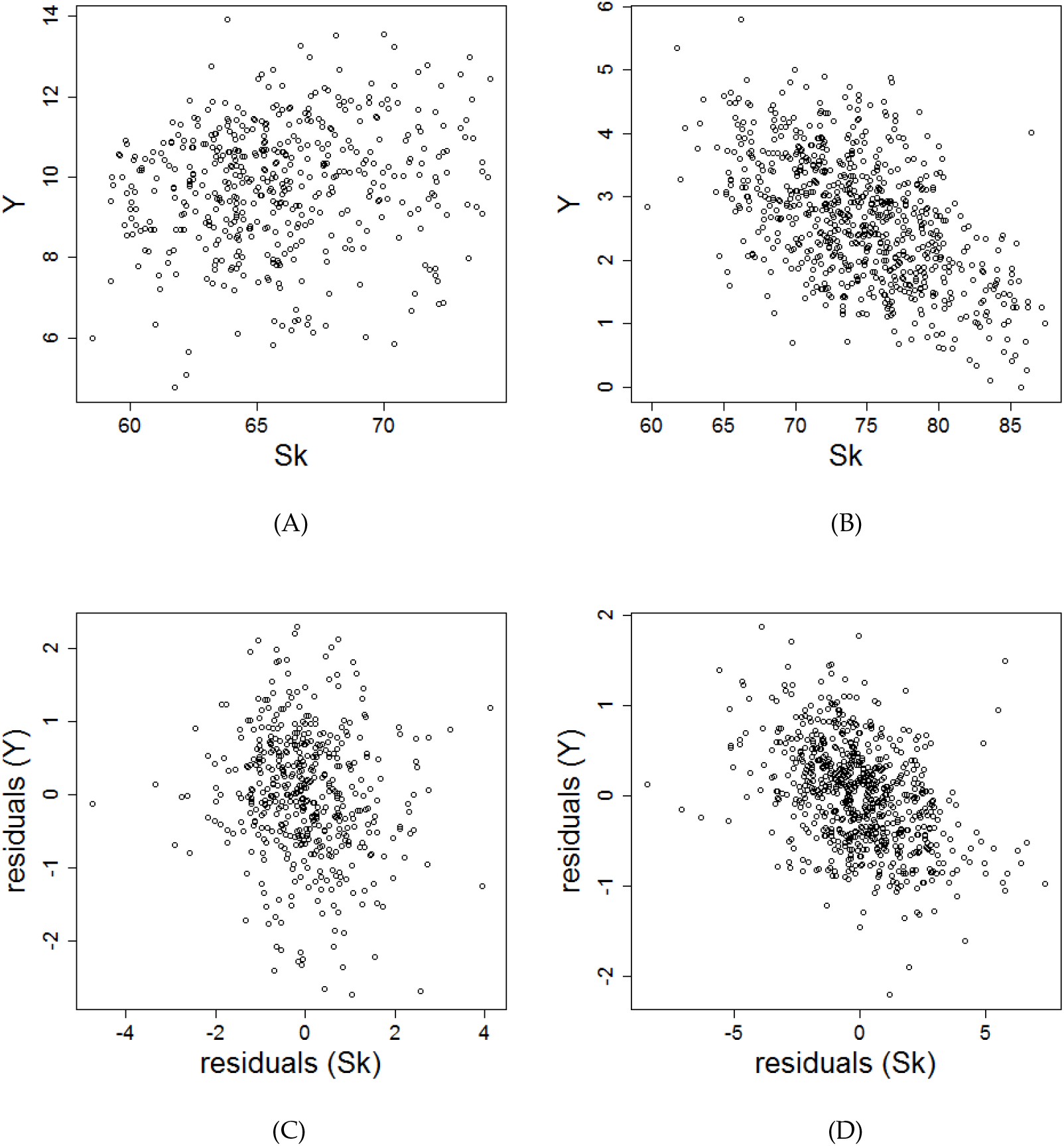
Yield (*ϒ*) versus silking (*Sk*), in a trial without heat and drought stress (Kar12W, (A)), and in a trial with severe stress (Bol12R, (B)). Panels (C) and (D) show the corresponding residuals from a bivariate multi-trait model.

**Figure S4.**
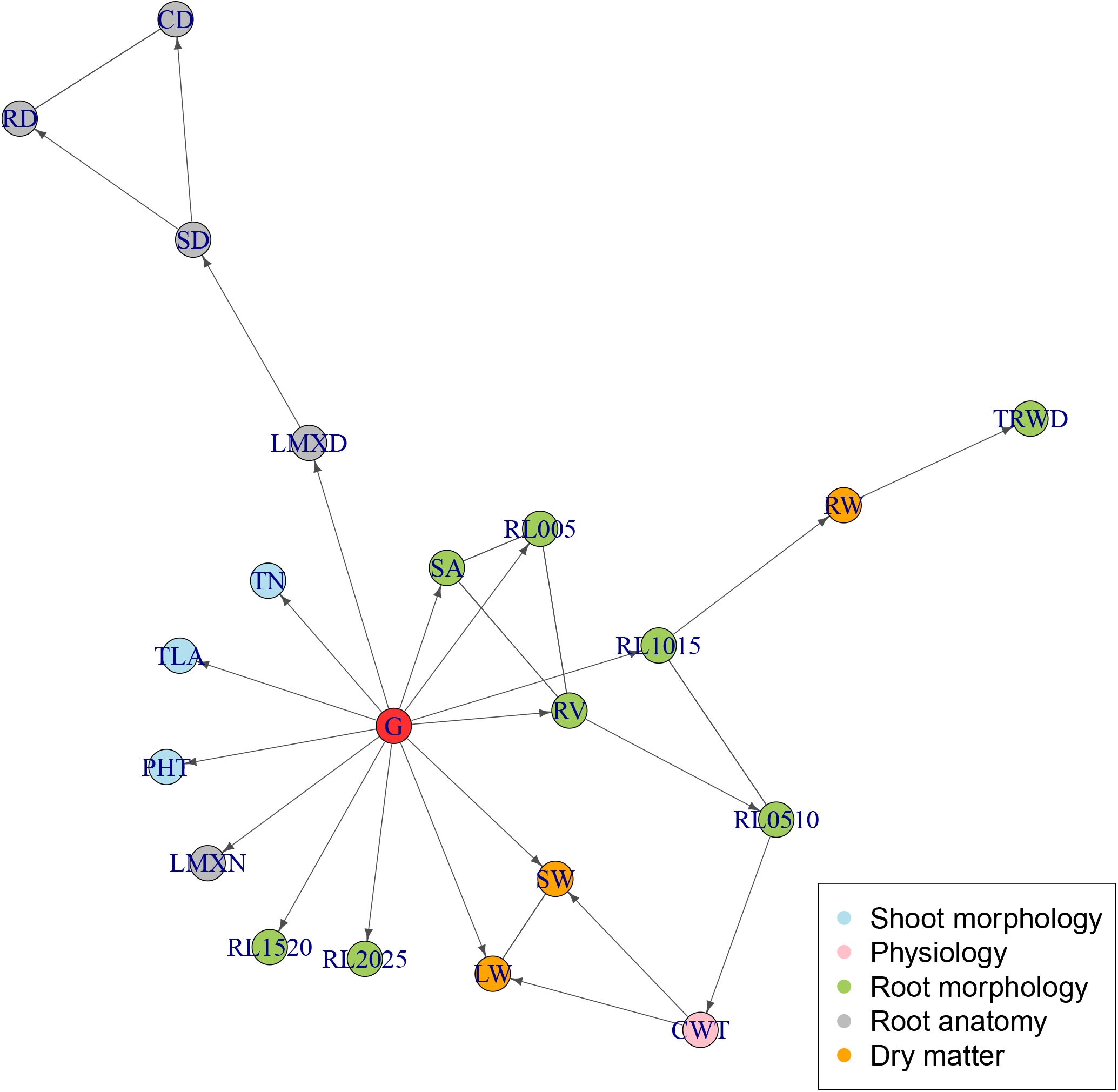
PCgen-reconstruction for the rice-data from Kadam *et al.* (2017), with *α* = 0.001. Five traits (MRL, ART, RL2530, RL3035 and RL35) are not shown, as they were completely isolated in the graph, without any connections to other traits or *G*.

**Figure S5.**
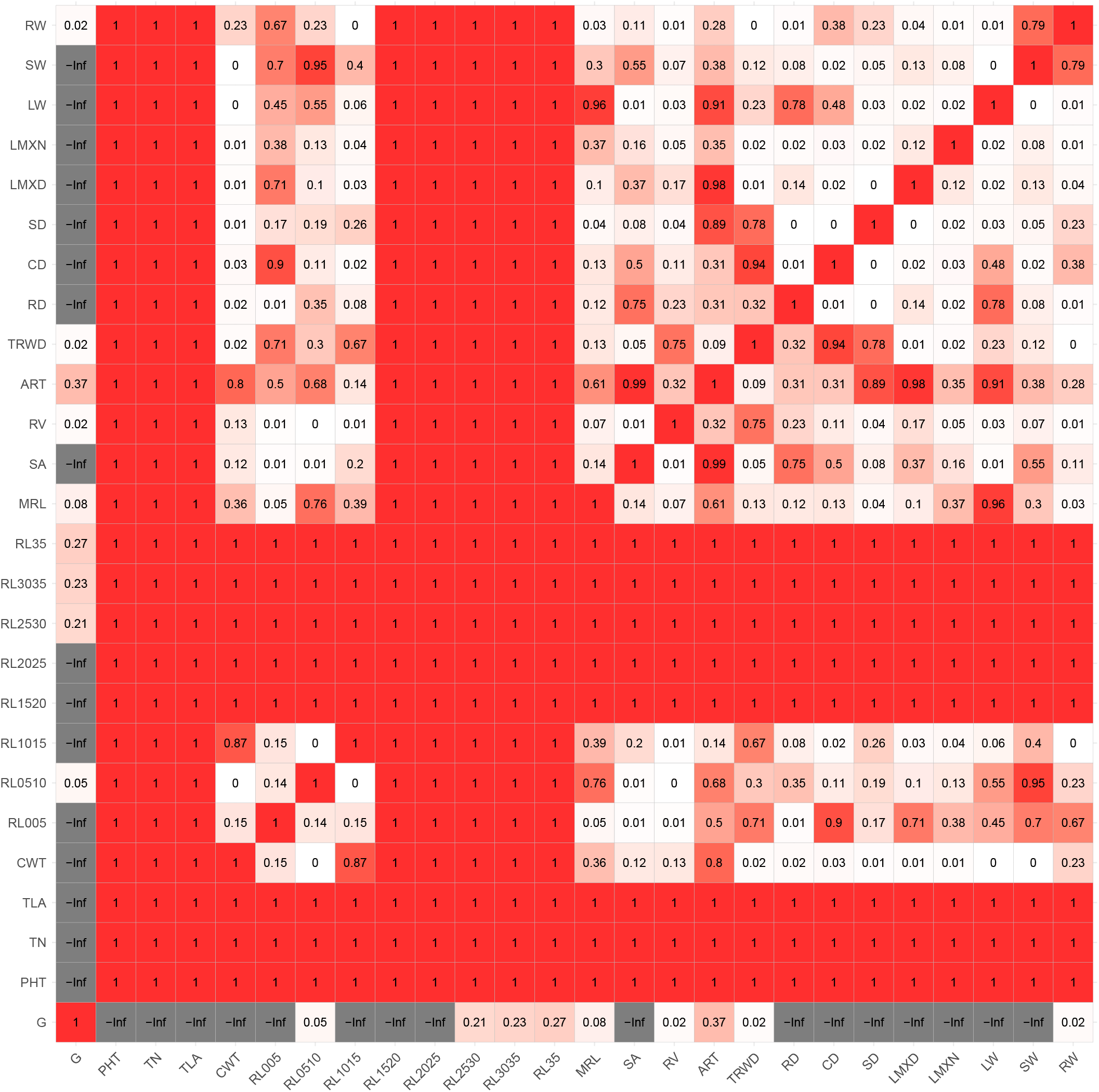
PCgen-reconstruction for the rice-data from Kadam *et al.* (2017), with *α* = 0.01: p-values. For each pair of nodes, the figure displays the highest p-value observed across all conditional independence tests performed for that pair.

## Footnotes

3 This is without restrictions on Σ_*E*_. In Appendix A.2 we conjecture an identifiability result for the case when Σ_*E*_ is diagonal.

4 We will see below that Γ can be computed from sums of products of path coefficients, without inverting Λ. Note that in the residual and genetic covariances in equation (5), we get Γ_*t*_ in front instead of Γ, as we *post*-multiply with Γ.

5 In this particular example, Γ = (*I* − Λ)^−1^ = (*I* + Λ).

6 alternatively, we could test the residual correlation

7 This does not directly describe the joint distribution of the traits at *t* = *T* (obtained by marginalizing over *t* = 0, … , *T* − 1, and typically represented by an ancestral graph (Richardson *et al.* 2002)), but for simplicity we nevertheless investigate to which extent we can reconstruct the summary graph, with observations taken at *t* = *T*.

8 For better interpretability we report correlations, although we test for zero genetic covariance (see File S1.4).

9 For statement (B) we will focus on the RC-test. The RG-test requires that the GBLUP **U**^*^ is close to the true matrix of genetic effects (**U**). Apart from the difficulty of obtaining good estimates of genetic and residual covariances, the quality of this approximation can be easily assessed using expressions for the prediction error variance (see e.g., Henderson (1975)).

10 The pcalg-package (Kalisch *et al.* 2012) has the addBgKnowledge option to add orientations (’background knowledge’) in the estimated CPDAG. This is however only done *after* running PC or a related algorithm, and is only allowed if compatible with the CPDAG.

11 Alternatively, conflicts can be represented with a bi-directed edge (↔), but the arrowheads do not have a causal interpretation

